# Charting the Heterogeneity of Colorectal Cancer Consensus Molecular Subtypes using Spatial Transcriptomics

**DOI:** 10.1101/2023.01.23.525135

**Authors:** Alberto Valdeolivas, Bettina Amberg, Nicolas Giroud, Marion Richardson, Eric J.C. Gálvez, Solveig Badillo, Alice Julien-Laferrière, Demeter Turos, Lena Voith von Voithenberg, Isabelle Wells, Amy A. Lo, Emilio Yángüez, Meghna Das Thakur, Michael Bscheider, Marc Sultan, Nadine Kumpesa, Björn Jacobsen, Tobias Bergauer, Julio Saez-Rodriguez, Sven Rottenberg, Petra C. Schwalie, Kerstin Hahn

## Abstract

The heterogeneity of colorectal cancer (CRC) contributes to substantial differences in patient response to standard therapies. The consensus molecular subtypes (CMS) of CRC is the most widely-used gene expression-based classification and has contributed to a better understanding of disease heterogeneity and prognosis. Nevertheless, CMS intratumoral heterogeneity restricts its clinical application, stressing the necessity of further characterizing the composition and architecture of CRC. Here, we used Spatial Transcriptomics (ST) in combination with single-cell RNA sequencing (scRNA-seq) to decipher the spatially resolved cellular and molecular composition of CRC. In addition to mapping the intratumoral heterogeneity of CMS and their microenvironment, we identified cell communication events in the tumor-stroma interface of CMS2 carcinomas. This includes tumor growth-inhibiting as well as -activating signatures, such as the potential regulation of the ETV4 transcriptional activity by DCN or the PLAU-PLAUR ligand-receptor interaction. Our data show the power of ST to bring the CMS-based classification of CRC to another level and thereby gain useful molecular insights for personalized therapy.

## 1. Introduction

CRC is a leading cause of cancer-related death worldwide with over 1.85 million diagnosed cases and 850 000 deaths annually^1^. CRC mortality rates have decreased in recent years as a result of treatments tailored to the molecular and pathological features of the different groups of patients^2^. However, the inter-patient and intra-tumor heterogeneity of CRC entails different responses to standard treatments, such as chemotherapy or immunotherapy, and provides a profound clinical hurdle^3^. CRC heterogeneity encompasses differences at the genomic, epigenomic and transcriptomic level as well as variations of the stroma and immune landscape, i.e. the composition of the tumor microenvironment (TME)^4^.

In 2015, the CRC subtyping consortium performed an integrative analysis on different large-scale gene expression datasets encompassing over 4000 CRC patients. Their study resulted in a gene expression-based subtyping classification of CRC into four CMS with distinguishing features^5^. CMS1 is hypermutated, microsatellite unstable and characterized by strong immune activation. CMS2 and CMS3 are epithelial subtypes, with CMS2 displaying marked WNT and MYC signaling activation, whereas CMS3 presents noticeable metabolic dysregulations. CMS4 features a prominent TGFβ activation, stromal invasion and angiogenesis^2^. The CMS classification framework is widely used and contributed to a better understanding of the diversity of CRC and disease prognosis. Nevertheless, its clinical impact on decision-making for CRC patients is still limited for several reasons^6, 7^. First, the CMS classification system fails to assign up to 13% of the CRC tumors, which are thought to display mixed or transitioning CMS phenotypes^8^. Moreover, it relies on bulk sequencing of CRC tumors, which lacks the resolution to comprehensively define the cell content and disentangle the heterogeneity of CRC tumors and their intricate TME^9^. Indeed, the CMS classification displays large intra-tumor heterogeneity as revealed by the assignment of different subtypes to samples extracted from the same CRC patient^10, 11^. Recently, several studies applied scRNA-seq on CRC samples to further reveal the diversity of and the dynamic relationships between cellular components of CRC tumors and their TME^7, 9, 12–14^. These analyses disclosed CMS features at the individual tumor cell level and stressed the high prevalence of multiple CMS phenotypes in the same patient^7, 9, 12^. Nevertheless, the spatial distribution and complex network of cellular interactions between the different CMS and their respective TMEs are still poorly understood.

Recent technological advances in next-generation sequencing- and imaging-based approaches have established the power of ST to systematically measure gene expression levels throughout tissue space^15^. In oncology, this technology adds another dimension to the classical histological readouts by enabling the integration of morphology, spatial localization and transcriptomic profile. Accordingly, ST paves the way towards a better understanding of cancer heterogeneity, TME composition, and complex cellular interactions. In this context, ST has been employed to study breast cancer^16^, prostate cancer^17^, melanoma^18^ and CRC. Concerning CRC, Wu et al.^19^ used ST to support their results obtained with scRNA-seq, describing immune pressure-driven evolution of metastasis and response to neoadjuvant chemotherapy. The ST data generated in that study were integrated with bulk transcriptomics of CRC patients by Peng et al.^20^ to explore the crosstalk between cancer-associated fibroblast and other components of the TME. In line with this, Qi et al^21^. revealed the interaction between FAP^+^ fibroblast and SPP1^+^ macrophages by using scRNA-seq and supporting their results with ST. In addition, Zhang et al.^22^ applied ST to study inflammatory patterns in proficient mismatch repair CRC. In these publications, the use of ST was mainly intended to support the results obtained with other technologies and did not specifically address the CMS of CRC.

In this work, we intend to improve our understanding of the spatial properties and heterogeneity of the CMS of CRC by applying ST on 14 samples from a heterogeneous cohort of seven CRC patients. Using a deconvolution-based approach, we first spatially characterized the cell type composition of the CRC tumors and their microenvironment. We associated the different CMS with distinctive molecular and morphological features and demonstrated the power of ST to dissect tumor heterogeneity. When we explored cell-to-cell communication events at the tumor-stroma interface in CMS2 carcinomas, we revealed well characterized and novel interactions including tumor growth-inhibiting as well as -activating signatures. Importantly, we supported our findings by analyzing an external ST CRC dataset. Overall, our results pave the way for a better understanding of CRC heterogeneity that builds on the current CMS characterization. We anticipate that future studies can take advantage of the power of ST to stratify treatments tailored to individual patients and thereby help the use of personalized and/or combinatorial therapy in CRC.

## 2. Results

### 2.1. ST and scRNA-seq-based deconvolution reliably reveal CRC cell type composition

We processed fresh-frozen (FF) resection samples obtained from seven CRC patients for ST using 10x Genomics VISIUM aiming at exploring spatial molecular heterogeneity in CRC (Fig. 1a, Table 1). We considered two serial sections per patient to generate technical replicates. Overall, quality control displayed favorable metrics with a median number of genes per spot ranging from 1233 to 5457 (Supplementary Fig. S1 and Methods). We evaluated the similarity between technical replicates and the heterogeneity among samples from different patients at the morphologic and transcriptomic level (Fig. 1b and Methods). The pathologists examined and annotated the samples regarding tissue type and cellular morphology.

**Figure 1:**
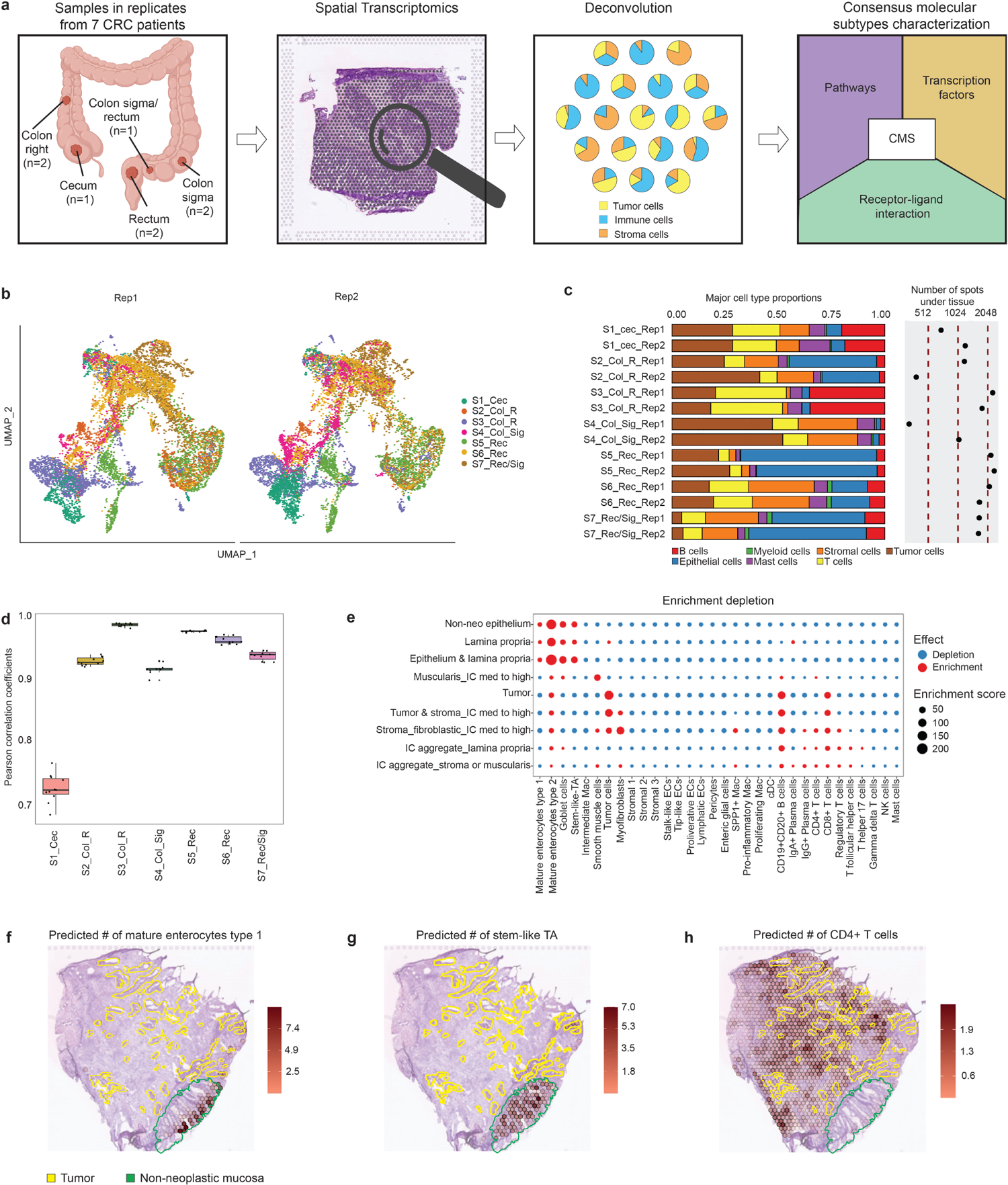
Study outline and deconvolution results matching histopathological annotations with high correlation between replicates. (a) Study outline displaying the anatomical localization of our set of CRC samples, their spatial transcriptomics processing and the deconvolution-based approach to characterize spatial features of CMS. (b) UMAP embedding of the gene expression measurements per spot split by technical replicates. Colors represent the different patients. (c) Proportions of major cell classes per sample as estimated by the results of the deconvolution approach. The right hand side of the plot displays the number of analyzed spots per sample. (d) Pearson’s correlation coefficients of the cell subtype abundance in small anatomical regions of variable size that were considered equivalent between technical replicates for all the patients. (e) Enrichment/depletion plot describing the association between cell type abundance as predicted by the deconvolution (x-axis) and the different anatomical regions as annotated by the pathologists (y-axis). The dot size represents the enrichment score (Methods), while the color represents enrichment (red) or depletion (blue). (f-h) Spatial mapping of the predicted number of mature enterocytes type I, stem-like TA and CD4+ T cells per spot matching the pathologists’ tissue annotation and expected cell type localization as illustrated for sample S6_Rec_Rep2

**Table 1:** Selected clinical information for the samples included in this study. *sample contains non neoplastic tissue; na: not applicable; - mutation profile was not assessed, IC: Immune cell content.

| Sample number | Localization | Diagnosis | Pre-treatment | Lymph node/liver metastasis | Mutation | Growth pattern and Immune cells |
| --- | --- | --- | --- | --- | --- | --- |
| S1_Cec<br>A551763 | Cecum | Adenocarcinoma, mucinous, moderately differentiated | no | yes/yes | BRAF V600E | Mucinous, IC low |
| S2_Col_R<br>A595688 | Colon (right) | Adenocarcinoma, moderately differentiated | no | yes/yes | no KRAS mutations | Tubular to cribriform, IC low |
| S3_Col_R<br>A416371 | Colon (right) | Adenocarcinoma, areas with moderate and poor differentiation | no | yes/no | - | 2 tumor types: I) tubular, IC low; II) extended solid, IC high |
| S4_Col_Sig<br>A120838 | Colon (Sigma) | Adenocarcinoma, moderately differentiated | no | yes/no | no KRAS mutations | Tubular to cribriform, IC low |
| S5_Rec<br>A121573 | Rectum | Adenocarcinoma, moderately differentiated* | no | yes/yes | - | Tubular to cribriform, IC low |
| S6_Rec<br>A938797 | Rectum | Adenocarcinoma, moderately differentiated* | no | no/no | - | Tubular, IC medium |
| S7_Rec/Sig<br>A798015* | Sigma/Rectum | non-neoplastic tissue | na | na | - | na |

To spatially map cellular composition per spot in our set of CRC samples, we applied the Cell2Location deconvolution method^23^ using as reference a recently published scRNA-seq dataset^12^ (Methods and Supplementary Table 1). In their study, Lee and colleagues explored the cellular landscape of different CRC subtypes, characterizing in detail cellular composition and suggesting intercellular interactions. They independently analyzed samples from a Korean cohort (23 patients) and a Belgian cohort (6 patients). Here, we focused on results obtained using the more comprehensive Korean cohort, which contains 65,362 well-annotated non-neoplastic-, CMS-classified neoplastic, immune-, stromal-, and endothelial cells. We note that the data and annotations from the Belgian cohort led to overall comparable results (Supplementary Note 1).

We then assessed the results of the deconvolution by exploring the similarity of the estimated cell type abundances per spot between technical replicates. First, we found highly comparable proportions between replicates when considering the major cell types present across samples (Fig. 1c and Supplementary Table 1). In contrast, proportions greatly differed across individuals: for instance, samples from patient S7_Rec/Sig comprised <= 5% tumor cells. This is consistent with the histology profile of this patient mainly containing non-neoplastic tissue, as described in Table 1. We next examined the correlation of cell subtype abundances in anatomical structures of variable size that were considered equivalent between replicates based on their gene expression profiles (Fig. 1d, Supplementary Table 1 and Methods). Excluding a single low quality sample (Methods), Pearson’s correlation coefficients were above 0.9, highlighting the similarity between technical replicates in the deconvolution results.

To further determine the accuracy of the deconvolution, we evaluated whether the predicted cell types are located at their corresponding anatomic tissue compartment. To achieve this, the pathologists manually assigned a category to each spot based on the tissue type and composition (Methods). Then, we computed proportions of the major cell types abundances in these different tissue categories (Supplementary Fig. S2). As expected, non-neoplastic epithelial cells were the most abundant in the non-neoplastic epithelium (89%), whereas T and B cells were the prevalent types in the immune cell aggregates (IC) located at the lamina propria (83%) and at the stromal or muscularis region (68%). In tumor-annotated spots, the most predominant categories were tumor cells (36%), T cells (26%) and B cells (25%). At the cell subtype level, non-neoplastic mucosal cells, such as mature enterocytes type 1 and 2, goblet cells and stem-like transiently amplifying (TA) cells, were significantly enriched in spots labeled as non-neoplastic epithelium, lamina propria or mixed (Fig. 1e and Methods). In contrast, tumor cells, CD19^+^CD20^+^ B cells and CD8^+^ T cells were mainly enriched in spots classified as tumor or tumor-stroma mixed. Other immune cells, including CD4^+^ T-cells, were mostly enriched in spots identified as immune-cell rich in stromal regions and/or IC. To frame these global results in our individual samples, we visualized the estimated number of different cell subtypes, overlaid with the pathologists’ tissue annotations (Figs. 1f-h and Supplementary Figs. S3-S9).

In summary, the estimated cell type abundances were highly comparable between technical replicates and their spatial distribution was in line with the pathologists’ assessment for all analyzed samples, demonstrating the reliability of the ST data and our deconvolution results. Consequently, we further used these to spatially characterize the CMS signatures and the TME in our set of CRC samples.

### 2.2. Spatially resolved consensus molecular subtyping of CRC and their key molecular features

Deconvolution-based estimates of CMS tumor cell proportions revealed a predominance of CMS2 cells in the S2_Col_R (94%), S4_Col_Sig (98%), S5_Rec (81%) and S6_Rec (90%) patients (Fig. 2a); hereafter referred to as CMS2 tumors. A mixed abundance of CMS1 and CMS2 tumors was identified in the S1_Cec (49% and 41% respectively) and S3_Col_R (65% and 29% respectively) patients; hereafter designated mixed CMS1-CMS2 tumors. Of note, S1_Cec harbored a *BRAF^V600E^* mutation (see Table 1), in line with previous findings linking this mutation to the CMS1 phenotype^5^. In addition, we detected CMS3 tumor cell signatures in the S1_Cec (10%) and S5_Rec (16%) patients. In the non-neoplastic S7_Rec/Sig sample, the few spots displaying a tumorigenic signal were mainly classified as CMS3 (60%) and, to a lesser extent, as CMS1 (19%). This is in agreement with the study by Lee et al^12^, in which CMS3 tumor cells were also observed to co-occur with CMS1 or CMS2. The CMS4 signatures were minor and multifocally distributed in our samples, but overlapped with anatomical regions displaying an invasive phenotype, suggesting an accurate spatial mapping. Supplementary Figs. S10-S16 show the overlay of the pathologists’ tissue annotations with the estimated abundance of the different CMS tumor cells across our set of samples.

**Figure 2:**
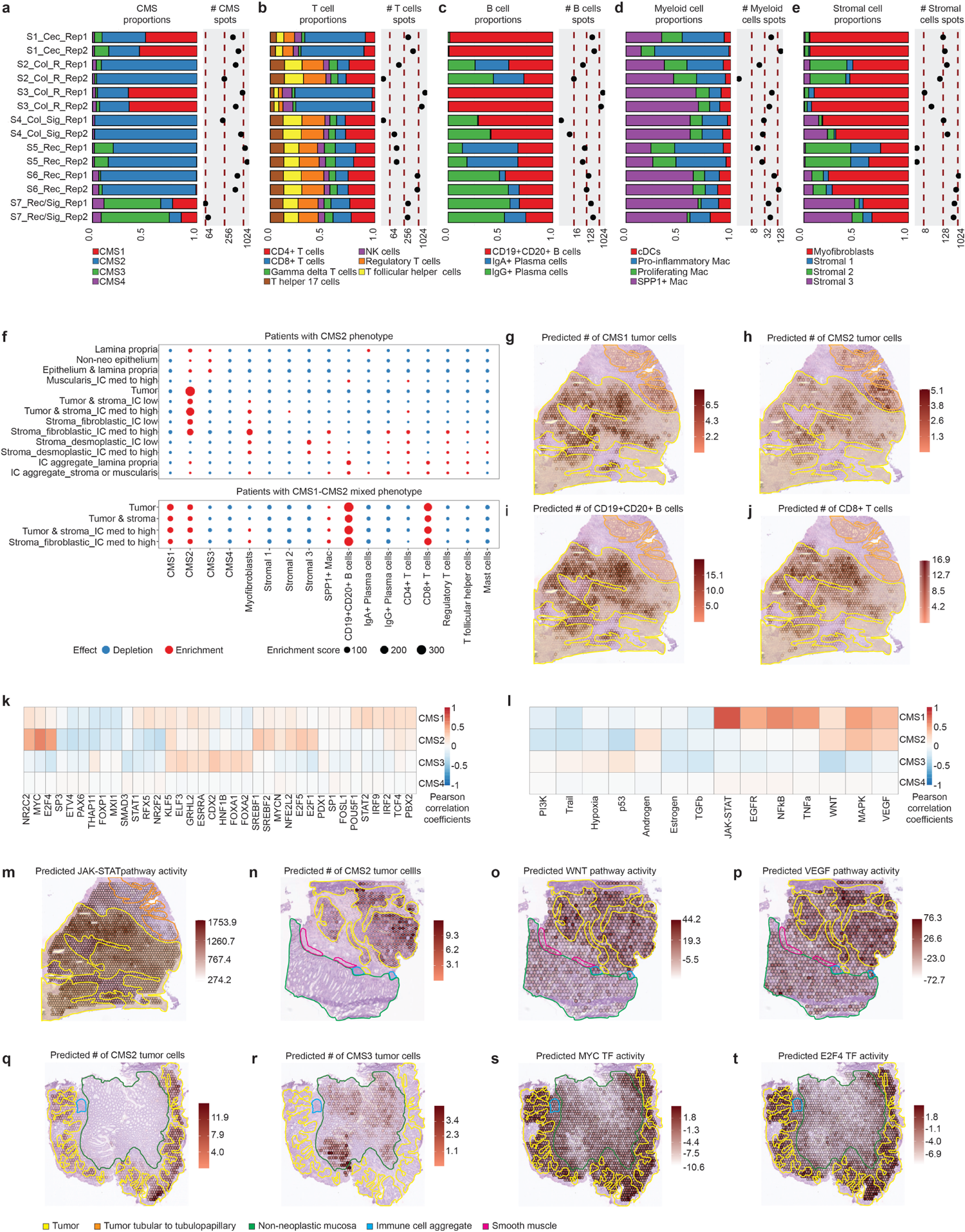
Consensus molecular subtyping of our set of CRC samples, characterization of their TME and spatially resolved mapping of their histological and molecular features. (a-e) Cell type proportions per sample as estimated by the results of the deconvolution. The number of spots containing an abundance of at least 20% of the specified cell types is also displayed. (f) Enrichment/depletion assessment of selected cell types (x-axis) in CMS2 and mixed CMS1-CMS2 tumors in the different tissue compartments defined by the pathologists’ spot classification (y-axis). (g-j) Spatial mapping of the predicted abundance of CMS1, CMS2, CD19^+^CD20^+^ B cells and CD8^+^ T-cells abundances overlaid with the pathologists’ tissue annotation in the S3_Col_R_Rep1 sample. Note the absence of CD19^+^CD20^+^ B cell and CD8^+^ T cells and the minor CMS1 signatures in regions of the tubulopapillary tumor (see Supplementary Fig. S12). (k) Per spot Pearson’s cross-correlation across all the samples between TF activities and CMS cell abundances. For visualization purposes, the 10 most highly correlated TFs in absolute value per CMS are shown. (l) Per spot Pearson’s cross-correlation across all the samples between pathway activities and CMS cell abundances. (m) Spatial mapping of the JAK-STAT pathway activity in sample S3_Col_R_Rep1 sample. Note the co-localization with the CMS1 signature. (n-p) Overlay of the predicted spatial CMS2 cell abundance, WNT pathway activity and VEGF pathway activity with the pathologists’ tissue annotations in the S2_Col_R_Rep1 sample. (q-t) Overlay of the predicted spatial CMS2 cell abundance, CMS3 cell abundance, MYC and E2F4 TF activities with the pathologists’ tissue annotations in the S5_Rec_Rep1 sample.

As an alternative approach towards CMS classification, we applied CMScaller^24^ on pseudo-bulk RNA-seq generated by either pooling together all the spots or only the tumor-annotated spots for each sample (Supplementary Fig. S17 and Methods). The patients with mixed CMS1-CMS2 tumors were both labeled as CMS1, suggesting that their large content of T and B cells (see Fig. 1c) was driving their classification towards CMS1 rather than the immune-deprived CMS2. Surprisingly, the S6_Rec patient was classified as CMS4 despite containing almost only CMS2-like tumor cells (90% of the total tumorigenic abundance) according to the deconvolution. The non-neoplastic S7_Rec/Sig sample was also categorized as CMS4. The high stromal content of these two patients may be driving these results, as suggested by previous studies reporting that CMS4 classification is highly influenced by marker genes of cancer-associated fibroblast and other stromal cells^9, 25, 26^. From the remaining patients classified as CMS2 by the deconvolution, uniquely S5_Rec was consistently classified as CMS2 by CMScaller. S2_Col_R and S4_Col_Sig were categorized as diverse subtypes in different replicates. These results underline how adjacent tissue components influence CMS classification approaches based on bulk transcriptomics, highlighting the importance of relying on scRNA-seq and ST to improve the characterization of CRC tumors.

In order to characterize the TME composition in our samples, we examined their immune and stromal cell proportions (Figs. 2b-e). As described above, mixed CMS1-CMS2 contained larger proportions of T and B cells than the other samples, as expected from the immune-rich phenotype associated with CMS1^5^. Their most abundant subtypes were CD8^+^T and CD19^+^CD20^+^ B cells, whereas they contained lower proportions of regulatory T cells (Tregs) as compared to CMS2 samples. Tregs inhibit antitumor immunity^27^ and therefore their presence in the surroundings of the CMS2 carcinomas may prevent immune infiltration. Myofibroblasts were the predominant stromal cell type in mixed CMS1-CMS2 tumors, whereas stromal cell types in CMS2 neoplasms were more heterogeneous. These results are in line with Lee et al.^12^ and Khaliq et al.^9^ who also reported a dominance of myofibroblast in CMS1 and CMS4 tissues.

Next, to associate these results with histological and spatial features, we computed the enrichment or depletion of the different cell subtypes in the tissue compartments defined by pathologists’ spot annotations (Fig. 2f and Methods). This analysis revealed the association of CMS1 and CMS2 signatures with the tumor-annotated spots. Of note, CMS3 signatures were confined to the non-neoplastic mucosa in all of our samples. This result can be attributed to the normal-like gene expression patterns of CMS3 tumors described in Guiney et al.^5^. Immune cells were predominantly associated with the stroma in CMS2, whereas in mixed CMS1-CMS2 tumors, CD19^+^CD20^+^ B cells and CD8^+^ T cells were also found in the neoplastic tissue. These results were further supported by an integrative co-localization analysis of the different cell subtypes based on their abundance maps (Supplementary Fig. S18 and Methods).

The overlay of the pathologists’ tissue annotations with the deconvolution results further revealed co-localization of CMS1 and CMS2 signatures in the S1_Cec and S3_Col_R samples (Figs. 2g-h). In S3_Col_R, a stronger CMS2 or CMS1 signature was associated with tubular or solid growth pattern respectively, as described in Thanki et al.^28^ (see Supplementary Fig. S12). As expected, immune cells, such as CD8^+^ T and CD19^+^CD20^+^ B cells, were abundant in the CMS1- and devoid in the CMS2-predominant region (Figs. 2i-j). In the non-neoplastic S7_Rec sample, the CMS1 signature was confined to the rectal gland, whereas CMS3 was associated with the mucosa (see Supplementary Fig. S16). We also delineated stromal 2 signatures as spatially adjacent to tumor lobes. Larger stroma bundles displayed a myofibroblast and a minor stromal 3 signature (Supplementary Fig. S19). Selected features of the TME of individual tumors and semiquantitative pathologists’ gradings are detailed in Supplementary Table 2.

To identify further molecular features associated with the different CMS in a spatially resolved manner, we performed an integrative analysis across all samples and explored the per spot correlation between tumor abundance and transcription factor (TF) and pathway activities (Figs. 2k-l and Methods). For the CMS1 tumor cells, we captured the expected correlation with JAK-STAT^29^ (Fig. 2m) and immune-related pathways, such as the TNF⍺^30^ and NFkB. In addition, a correlation with the EGFR and MAPK pathway was identified. The activation of the MAPK pathway is well known in the hypermutated CMS1^31^. For the CMS2 tumor cells, we found the expected correlation with the WNT and VEGF pathways^32^ (Figs. 2n-p). At the TF activity level, CMS2 abundance was associated with high expression of MYC-regulated genes^5^, whereas lower transcriptional MYC activities were detected in the non-neoplastic mucosa containing CMS3 signatures (Figs. 2q-s). Noteworthy, E2F4 and MYC TF activity maps displayed similar patterns suggesting an interplay between these TFs in the regulation of target genes implicated in CMS2^33^ (Fig. 2t and Supplementary Fig. S20). Hence, our deconvolution-based approach spatially mapped the different CMS and TME cell types, revealing their association to key molecular and histological features. In addition, we showed the ability of ST to detect and characterize spatially heterogeneous CMS phenotypes.

### 2.3. ST maps the inter-patient and intra-patient heterogeneity of CMS2 tumors

ST enables the exploration of the transcriptomic diversity of tumors and their TME at an unprecedented level. While the integrative analysis of our samples captured the core molecular features of the different CMS subtypes, we subsequently performed an in-depth assessment of individual samples to delineate the inter-patient and intra-tumor heterogeneity of our four CMS2 carcinomas (S2_Col_R; S4_Col_Sig; S5_Rec; S6_Rec).

In order to depict inter-patient heterogeneity between CMS2 tumors, we first extracted all tumor-annotated spots (Supplementary Fig. S21 and Methods). These spots possessed CMS2-dominated transcriptomes as their CMS2 cell abundance ranged from 65% to 84% of the total estimated number of cells (Supplementary Fig. S22). Nevertheless, there were noticeable differences between patients, as revealed by differential gene expression, pathway and TF activity analyses (Figs. 3a-d, Methods and Supplementary Table 3). This observation is in accordance with previous studies suggesting that CMS2 tumors are highly heterogeneous^9, 34^. In tumors obtained from the S4_Col_Sig and S5_Rec patients, genes involved in mTORC1 signaling, a known player in the progression of normal to neoplastic cells in CRC at early stages of the tumorigenesis^35^, were overrepresented. However, genes participating in the mTORC1 pathway were differentially expressed between these two patients (Supplementary Table 3), suggesting alternative signaling cascades. For instance, *NUPR1*, which was shown to promote metastasis in CRC by activating the PTEN/AKT/mTOR signaling pathway^36^ was only highly expressed in CMS2 tumor cells derived from the S4_Col_Sig patient (Fig. 3b). At the pathway level, we identified lower EGFR signaling activity in the tumor spots originating from the S2_Col_R and S4_Col_Sig patients (Fig. 3c and Supplementary Fig. S23). KRAS mutations were screened and not detected in these patients (see Table 1), in line with the assumption that the EGFR signaling pathway is usually activated in CMS2 tumors at the expense of KRAS mutations^37^. FOXM1 displayed higher transcriptional activity in S6_Rec, when compared to the tumor spots derived from the other patients (Fig. 3d and Supplementary Fig S24). This signal might be related to residual stromal cells in spots annotated as tumor and is consistent with the pseudo-bulk classification of this tumor as CMS4.

**Figure 3:**
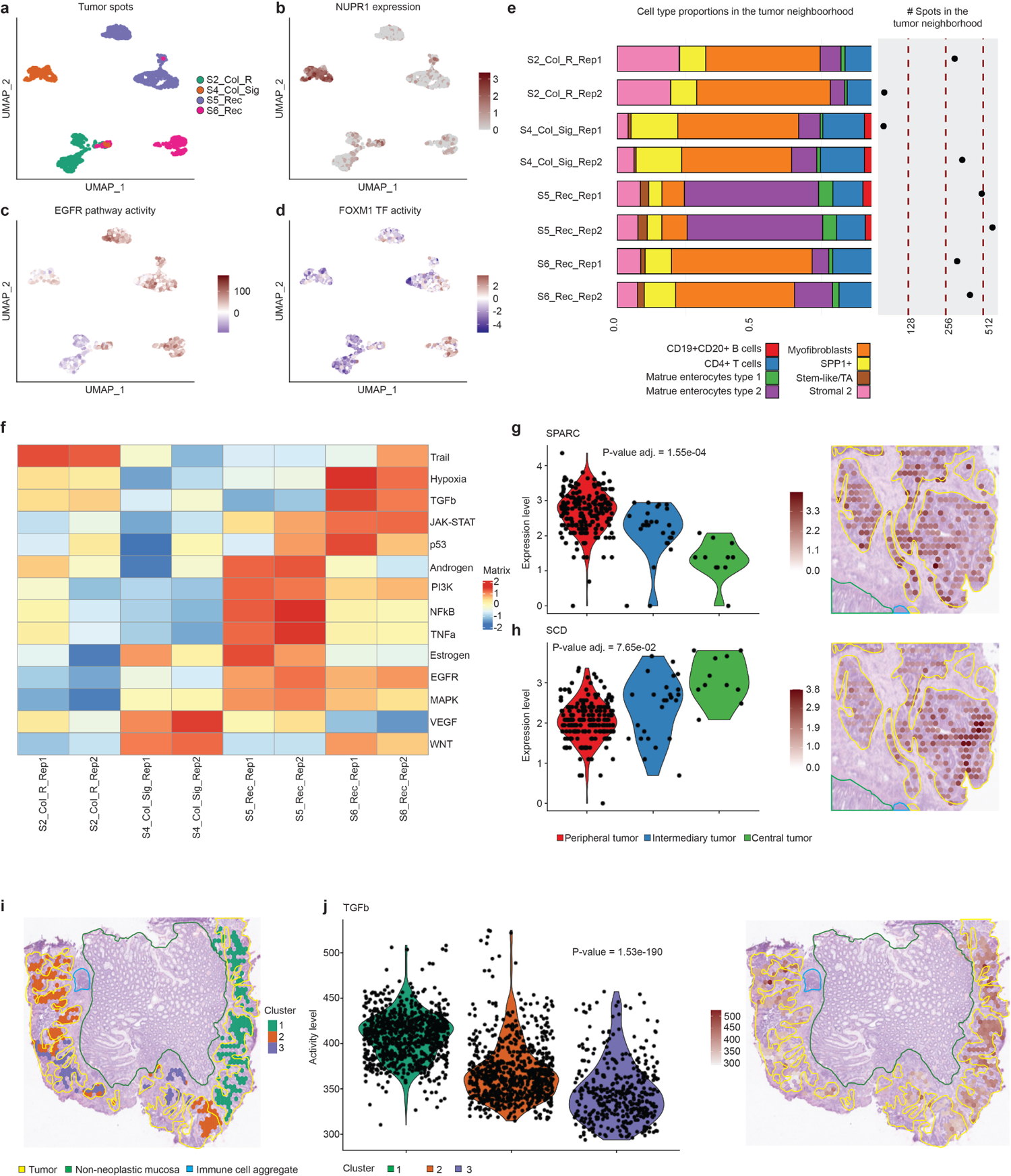
Inter- and intra-patient heterogeneity in CMS2 tumors and their TME in terms of cell composition and different molecular features. (a-d) UMAP embeddings of the gene expression measurements in tumor annotated spots which were colored by different criteria: a) per patient, b) per the expression of the NUPR1 gene, c) per activity of the EGFR pathway and d) per activity of the FOXM1 TF. (e) Cell type proportions in the tumor-surrounding spots per sample as estimated by the results of the deconvolution approach. The number of tumor-surrounding spots for the different samples is also displayed. (f) Differential pathway activity computed on pseudo-bulk RNA-seq generated from the tumor-surrounding spots for the different samples. (g-h) Gene expression gradients of SPARC and SCD in the different anatomical regions of tumor-annotated spots in the S2_Col_R_Rep1 sample. A Wilcoxon rank sum test was conducted to assess the significance of the gene expression variation (p-value adjusted). (i) Overlay of the spatial mapping of the clustering at subspot enhanced resolution of the tumor-annotated spots with the pathologists’ tissue annotations in the S5_Rec_Rep1 sample. (j) Spatial mapping and violin plots per group of the TGFb pathway activity at the enhanced subspot resolution in the S5_Rec_Rep1 sample. A Kruskal-Wallis statistical test was performed to assess whether the pathway activities in the different subclusters originated from the same distribution (p-value).

Multiple factors, such as the inherent heterogeneity of CMS2 tumor cells or their different anatomical origin can account for these inter-patient transcriptomic differences. The composition and spatial organization of the TME may also have a major impact on their transcriptomic profile. ST enables assessing the latter in a unique manner. Towards this end, we selected the spots surrounding CMS2 tumors and explored their cell type abundance profiles for each sample (Fig. 3e and Methods). Then, using these spots, we generated pseudo-bulk RNA-seq data and evaluated differential pathway activity among patients (Fig. 3f and Methods). Interestingly, we observed a depletion in the number of myofibroblasts for the S5_REC patient (Supplementary Fig. S25a). This might explain the lower activity of the TGFβ pathway^38^ in the tumor and surrounding regions and be indicative of earlier tumor stages with reduced stromal content. We also detected an enrichment of mature enterocytes type 2, highlighting the close morphological and spatial association of non-neoplastic and neoplastic cells in this sample (Supplementary Fig. S25b), and underlining the potential of ST to assess surgical margins. On the other hand, S4_Col_Sig displayed a higher proportion of SPP1^+^ macrophages (Supplementary Fig. S25c), which are key to creating an immunosuppressive TME^39^. This could relate to the lower activities of immune response associated pathways such as NFkB and TNFa signaling observed in the tumor of this patient.

CMS2 tumors can also exhibit a large degree of heterogeneity within the same patient. To illustrate and characterize this intra-tumor heterogeneity, we first selected the S2_Col_R_Rep1 sample and categorized its tumor annotated spots in three different regions based on their distance to non-tumor annotated spots (Methods). The regions were termed as peripheral-, intermediate -and central tumor. In this manner, we established a zonation model allowing us to investigate the genes and processes that are more active in the tumor boundary or in its internal solid area. Differential gene expression analysis between these zones revealed the anticipated overrepresentation of genes involved in EMT and angiogenesis in the peripheral tumor^40^ (Supplementary Fig. S26a, Supplementary Table 4 and Methods). In this region, we detected high expression of several fibroblast-specific genes, such as *FBLN1* or *COL3A1* (Supplementary Fig. S27), which could originate from few reminiscent stromal cells located in tumor-annotated spots. A more intriguing result is the upregulation of *SPARC* (Fig. 3g), a gene whose expression in cancer cells (not in stromal cells) was recently shown to control tumor progression and prognosis in CRC^41^. In the central solid tumor, we identified several upregulated genes known to be involved in hypoxic response and cholesterol homeostasis (Supplementary Fig. 26b and Supplementary Table 4), in accordance with the low oxygen conditions expected in this region^42^. Fig. 3h shows the spatial expression pattern of *SCD*, which we consider of particular interest as its upregulation in hypoxic tumors is linked to the metabolic reprogramming required to promote growth and metastasis of cancer cells^43^, including CRC^44^. Additional examples of upregulated genes in the central tumor were *INSIG1* and *MELTF* (Supplementary Fig. S28), whose role in cancer is not yet clearly defined.

We then selected the S5_Rec_Rep1 sample and sub-clustered the tumor-annotated spots using gene expression at enhanced resolution (Fig. 3i and Methods). The clustering revealed three different regions defined by various differentially expressed genes (Supplementary Fig. S29), biological processes (Supplementary Fig. S30 and Supplementary Table 5) and pathway activities (Methods). Of note, some tumor-associated pathways, such as EGFR and MAPK, displayed marked differences between subclusters (Supplementary Figs. S31a-b). On the other hand, the activity patterns of WNT and VEGF pathways, hallmarks of CMS2 tumors, presented a more homogeneous distribution among subclusters (Supplementary Figs. S31c-d). We further detected increased TGFβ pathway activity in subcluster number 1 (Fig. 3j), pointing to the tumor regions that most likely proliferate and undergo metastatic processes^45^.

Together, our results illustrate the ability of ST to delineate the inter- and intra-tumor heterogeneity of CMS2 carcinomas and to define the composition of their TME. This is crucial to better understand differential patient response to treatments such as immunotherapy. In addition, the differential spatial patterns of key molecular processes involved in cancer progression, such as high TGFβ pathway activity, can help designing tailored treatments or new combination therapies.

### 2.4. ST charts cell-to-cell communication processes modulating CMS2 tumor progression

ST reveals the cellular organization within tissues, providing a unique opportunity to study cell communication events. Accordingly, we explored such processes occurring in the tumor-stroma interface and their potential role in CMS2 tumor progression. In addition, we independently used the scRNA-seq data from Lee et al.^12^ to support and refine our results.

In the previous section, we used ST to highlight and characterize the heterogeneity of CMS2 tumors and their TME at the transcriptome level. To study common biological processes across our CMS2 tumor samples, we hypothesized that some transcriptional programs modulating tumor progression may display higher similarity than individual gene expression patterns. We consequently merged the spots from our four samples displaying an unequivocal CMS2 phenotype (S2_Col_R; S4_Col_Sig; S5_Rec; S6_Rec), and clustered them based on their TF activity profiles (Methods). Indeed, the UMAP embedding and clustering revealed higher similarity than that of gene expression-based results (Fig. 4a and Supplementary Fig. S32). Cluster 0, hereafter referred to as the tumor cluster, contained spots mainly annotated as *tumor* (49%) and *tumor&stroma_IC med to high* (26%) across replicates and patients (Fig. 4d and Supplementary Figs. S33-S34). In the same line, cluster 1, hereafter referred to as the TME cluster, included spots predominantly annotated as stromal regions (63% as *stroma_fibroblastic_IC med to high* and 20% as *tumor&stroma_IC med to high*), which were lying in the neighborhood of the tumor in every sample (Fig. 4d, Supplementary Figs. S33-S34). As expected, we found MYC and E2F4 among the most differentially activated TFs in the tumor cluster (Fig. 4b, Supplementary Fig. S35 and Methods). In the TME cluster, we identified several TFs known to play a pivotal role in cancer progression such as JUN^46^ and ETS1^47^ (Fig. 4c and Supplementary Fig. S35). Of note, cluster 6 also presented higher transcriptional activity of MYC and E2F4 (Supplementary Figs. S34-S35) and spots mainly annotated as *tumor* (50%) and *tumor&stroma_IC med to high* (30%) (Fig. 4d). However, almost all the spots belonging to cluster 6 come from the S6_Rec patient (see Supplementary Figs. S32-S34). It was therefore not considered for subsequent analysis.

**Figure 4:**
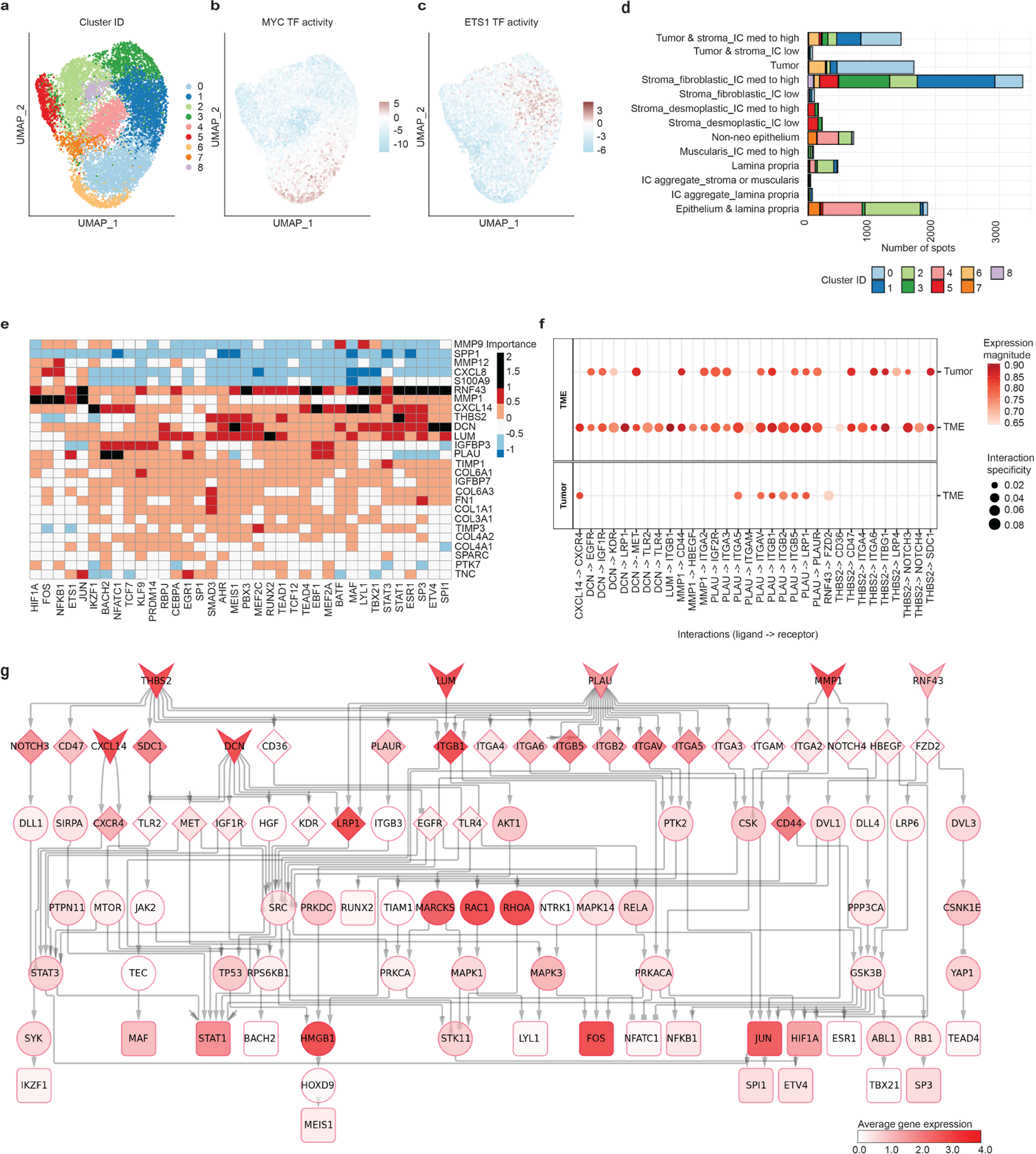
Clustering based on TF activities to study cell communication events at the tumor-stroma interface of CMS2 tumors. The signaling cascades triggered by those events and leading to transcriptional activities related to tumor progression were also investigated. (a-c) UMAP embedding of the TF activity profiles for our set of CMS2 samples. The spots were colored following different criteria: a) per cluster group, b) per activity of the MYC TF, and c) per activity of the ETS1 TF. d) Number of spots belonging to the different categories of pathologist’s annotations and clusters as inferred from the TF activity profiles. (e) Misty results showing the potential importance of ligands (rows) expression on TF (columns) activity. The ligand-TFs relationships with an importance score over 1 are represented as black slots and were further investigated. (f) Top ligand-receptor interactions at the tumor stroma interface predicted by LIANA. The left panel shows the source of the interaction (ligands) and the right the target (receptors). (g) Signaling cascades potentially linking ligands (V shape) to their downstream TF targets (triangles) according to Misty predictions. The downstream signaling cascades go first through the top predicted receptors by LIANA and then to intermediary signaling proteins (ellipses). The color of the nodes indicates the average expression of these genes in the TME cluster. Network edges can represent stimulatory (arrows) or inhibitory (squares) interactions.

To study cell communication events, we first selected highly expressed ligands in the tumor and TME clusters (Methods). We then used Misty^48^ to estimate the potential influence of the expression of these ligands on modulating the activity of TFs that are operating in the TME, such as the aforementioned JUN and ETS1 (Fig. 4e and Methods). To place these results into a mechanistic context, we inferred the most likely signaling cascades connecting the top predicted ligand-TF associations (black squares in Fig. 4e). To do so, we first investigated inter-cellular ligand-receptor interactions between the tumor and the TME clusters (Fig. 4f and Methods). Then, using a network-based approach, we connected the top predicted ligand-receptor interactions with our set of active TFs in the TME (Fig. 4g and Methods). To define the cell types involved in these processes, we independently computed TF activity and inferred ligand-receptor interactions on the patients classified as CMS2 in the scRNA-seq dataset published by Lee et al.^12^ (Figs. 5a-d, Supplementary Fig. 36 and Methods).

We predicted that the stroma-secreted DCN may modulate the transcriptional activity of ETV4, MEIS1 and SPI1 (Figs. 4e, 5e-f and Supplementary Figs. S37a-c). ETV4 is known to promote tumor invasion in CRC by regulating the expression of metalloproteinases^49^. As revealed by our network-based analysis (Fig. 4g), this regulation occurs in response to the MAPK signaling pathway, which is in turn activated by the binding of EGF to its putative EGFR receptor^50^. Substantial abundance of DCN can obstruct that interaction, as it directly binds to EGFR and downregulates its expression, preventing tumor progession^51^. Our ligand-receptor analysis identified the DCN-EGFR interaction targeting stromal or CMS2 tumor cells (Figs. 4f, 5c-d). In line with these findings, the average transcriptional activity of ETV4 appears to be lower in cell types targeted by the interaction, such as myofibroblasts, than in non-targeted cells, like macrophages (Fig. 5a). The family of MEIS TFs can act as tumor suppressors or oncogenes under different cellular conditions and cancer types and their target genes are widely misregulated in CRC^52^. Our results suggest that DCN may influence the transcriptional activity of MEIS1 through the downstream signaling of the SRC kinase family (Fig. 4g), which is known to promote metastasis and cause chemotherapeutic drug resistance in CRC^53^. Finally, SPI1 participates in the transcription of several genes involved in immune cell differentiation and tumor progression^54^. Our network-based analysis revealed that DCN may modulate the activity of SPI1 via the well-characterized STAT-mediated EGFR signaling axis^55^ (Fig. 4g). Of note, we identified additional interactions where DCN is known to play a protective role in tumor progression. Namely, DCN interacts with the MET receptor to inhibit tumor growth and angiogenesis and promotes inflammation via interactions with TLR2 and TLR4 receptors^56^ (Supplementary Fig. 37d). Given its eminent protective role, high expression levels of DCN in stromal cells around CMS2 carcinomas could be expected to be indicative of non-proliferative tumor regions. On the other hand, DCN expression levels may also be associated with a protective response against tumor progression associated events, such as intense metastatic activities (Fig 5g). In sum, our results provide mechanistic insights about how DCN expression may modulate signaling cascades involved in CRC tumor progression.

**Figure 5:**
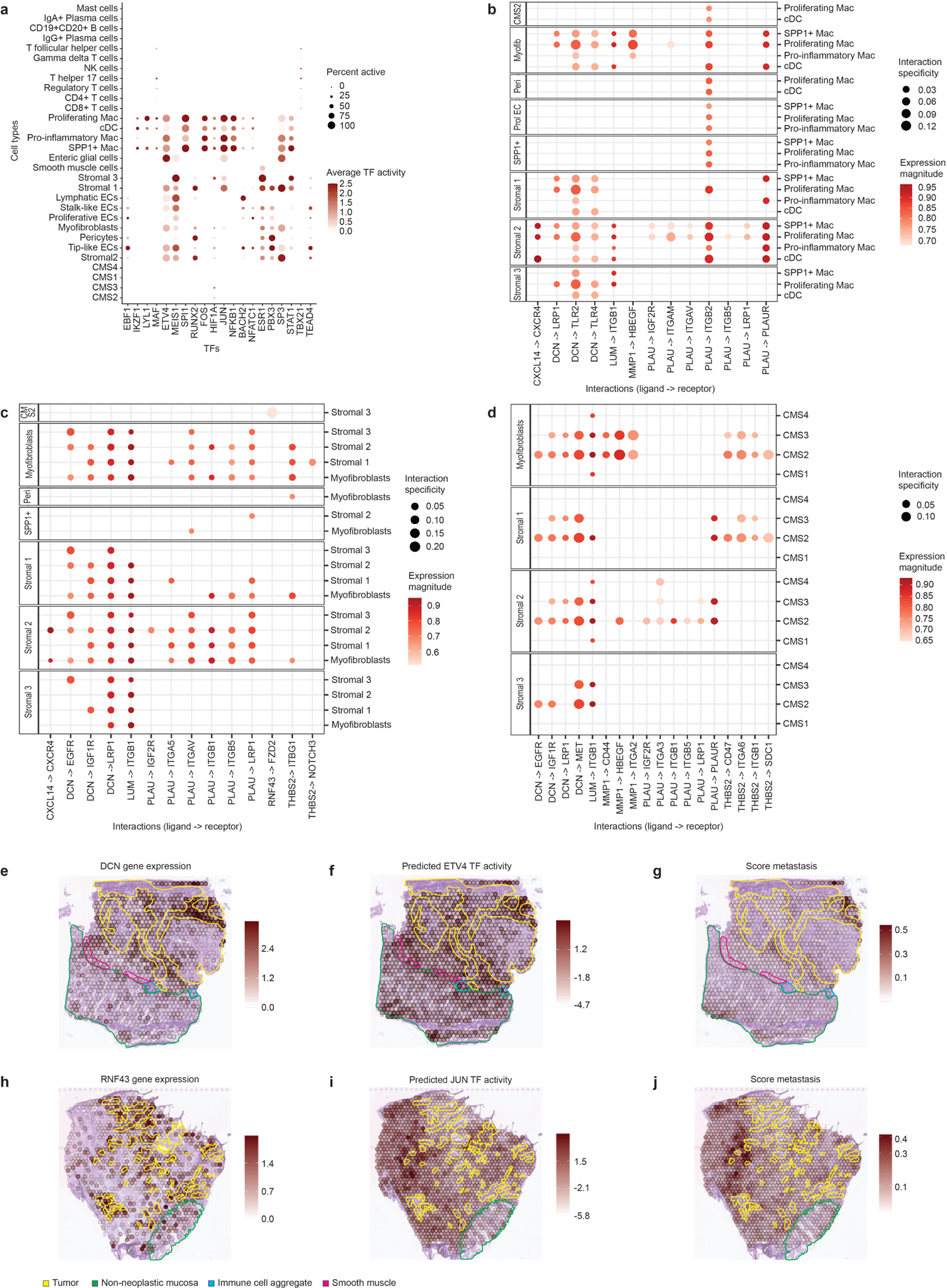
Transcription factor activity and ligand-receptor interactions in the scRNA-seq from Lee et al. Spatial maps showing gene expression, TF activity and a score for selected tumor-associated processes. (a) Average TF activity per cell type. The percentage of cells of a given type where the TF is active is represented by the size of the circle. (b-d) Ligand-receptor interactions between the different cell types overlapping with the interactions predicted in our ST data. The left panel shows the source of the interaction (ligands) and the right the target (receptors): b) target cell types are myeloid cells, c) target cell types are the major stromal cell populations, and d) target cell types are the different CMS tumor cell types. (e-g) Overlay of the DCN gene expression, the predicted ETV4 TF activity and the metastasis score with the pathologists’s tissue annotations in the S2_Col_R_Rep1 sample. (h-j) Overlay of the RNF43 gene expression, the predicted JUN TF activity and the metastasis score with the pathologists’s tissue annotations in the S6_Rec_Rep2 sample.

We also found that the expression of the transmembrane protein RNF43 may modulate the activity of several TFs operating in the TME, including JUN and TEAD4 (Figs. 4e, 5h-i and Supplementary Figs. S38a-b). Previous studies in colorectal and pancreatic cancer suggest that RNF43 possesses a tumor suppressor role through the inhibition of the WNT signaling pathway. Indeed, overexpressed RNF43 was shown to target the WNT receptors of the Frizzled family^57–59^ for degradation. JUN plays a well-known role in neoplastic transformation and is activated via the non-canonical WNT pathway^60^. The TEAD family (TEAD1-4) of TFs are key components of the Hippo signaling pathway and bind to YAP to promote the transcription of genes involved in cell migration and angiogenesis^61^. Interestingly, YAP is also a negative regulator of the WNT pathway^62^. Our network-based analysis revealed the interaction between YAP1 and TEAD4 and also other crucial components of the WNT pathway, such as the disheveled family of proteins (DVL-1 and DVL-3) (Fig. 4g). Interestingly, we consistently predicted an interaction between RNF43 and FZD2 in both the ST (Fig. 4f and Supplementary Fig. S38c) and scRNA-seq data targeting stromal 3 cell populations (Fig. 5c and Supplementary Fig. 38d). However, this interaction is well-documented to occur in the intracellular domain of RNF43 in tumor cells^63^, with few studies reporting a potential extracellular interaction^64^. It is therefore likely that the ligand-receptor analysis is capturing indirect expression associations. Taken together, higher expression levels of *RNF43* may lead to increased degradation of the receptors of the WNT pathway. As a consequence, the transcriptional activity of TFs downstream of these receptors is affected and may be indicative of tumor regions with lower metastatic activity (Fig. 5j). Noteworthy, these particular biological processes may not directly result from cellular interactions at the tumor-stroma interface.

We further predicted other ligand-TF associations with potential protective roles against tumor progression. For instance, the THBS2 secreted ligand was predicted to have an influence on the activity of STAT1 (Fig. 4e and Supplementary Figs. S39a-b). THBS2 is known to have anti-tumor progression properties by interacting with CD36 to promote anti-angiogenic processes (Fig. 4f and Supplementary Figs. S36, S39c-d)^65^. On the other hand, we found other modulations suggestive to promote tumor cell growth and migration. For example, the expression of *MMP1,* a matrix metalloproteinase involved in cancer progression through degradation of the extracellular matrix^66^, was predicted to have an effect on the activity of the FOS TF (Fig. 4e and Supplementary Figs. S40a-b). Another interesting result is the predicted interaction of the secreted ligand PLAU with its putative receptor PLAUR (Fig. 4f and Supplementary Figs. S41a-b). The binding of PLAU and PLAUR is known to trigger the degradation of extracellular matrix components, promoting tumor invasiveness^67^. We predicted this interaction to be occurring between myofibroblast cells (source of the interaction) and macrophages or conventional dendritic cells (cDCs) (target of the interaction) (Fig. 5b and Supplementary Figs. S41c-d). This is in line with a study in prostate cancer, associating PLAU-PLAUR interaction with macrophage infiltration^68^. Finally, we also found that the CXCL14 chemokine may potentially have a downstream impact in the transcriptional activity of MAF (Fig. 4e), which has been shown to regulate the immunosuppressive function of tumor-associated macrophages^69^. Interestingly, a stabilized dimeric peptide containing part of CXCL14 amino acid residues has been proposed as an anticancer treatment^70^. Taken together, our analysis revealed ligands potentially modulating the activity of TFs known to play a pivotal role in CMS2 tumor progression, highlighting the power of ST to study cell communication processes in CRC.

### 2.5. External ST CRC data confirms deconvolution-based subtyping, heterogeneity patterns and predicted cell communication events

To further validate our results, we used independent CRC ST data from a recent publication with morphological features suggestive of CMS2^19^. This ST dataset contains four samples from primary CRC tumors and their corresponding four liver metastases. Two of the patients were untreated (*Unt*) and the other two were treated with neoadjuvant chemotherapy (*Tre*).

We first applied our deconvolution-based approach to characterize this set of samples. Of note, the deconvolution was also applied to the samples originating from the liver. Consequently, some cell types from the scRNA-seq reference are not expected to match liver tissue, mainly the non-neoplastic intestinal epithelial cells. The proportions of the major cell types present in the tissues (Fig. 6a) revealed a very reduced tumor content in the ST-colon2_Unt, ST-colon3_Tre and ST-liver3_Tre samples, in accordance with their histology. For these samples, only around 4% percent of the total cell abundance was mapped to tumor cells. All the samples, including the liver metastases, presented a dominant CMS2 phenotype with over 80% of the total tumor cells mapped to this subtype (Figs. 6b-c and Supplementary Figs. S42-S45). In agreement with our previous results, CMS3 signatures were confined to the non-neoplastic mucosa and CMS4 signals were minor and multifocally distributed. The abundance of CMS1 tumor cells was almost negligible in these samples. Interestingly, CMS2 signals were substantial and overlapped with the histology of the liver tumors suggesting a conservation of the CMS phenotype in metastasis. (Fig. 6d and Supplementary Figs. S46-S49). To further characterize these samples, we explored their relative content of the different types of T cells, B cells, myeloid cells and the main stromal cells (Supplementary Figs. S50-53).

**Figure 6:**
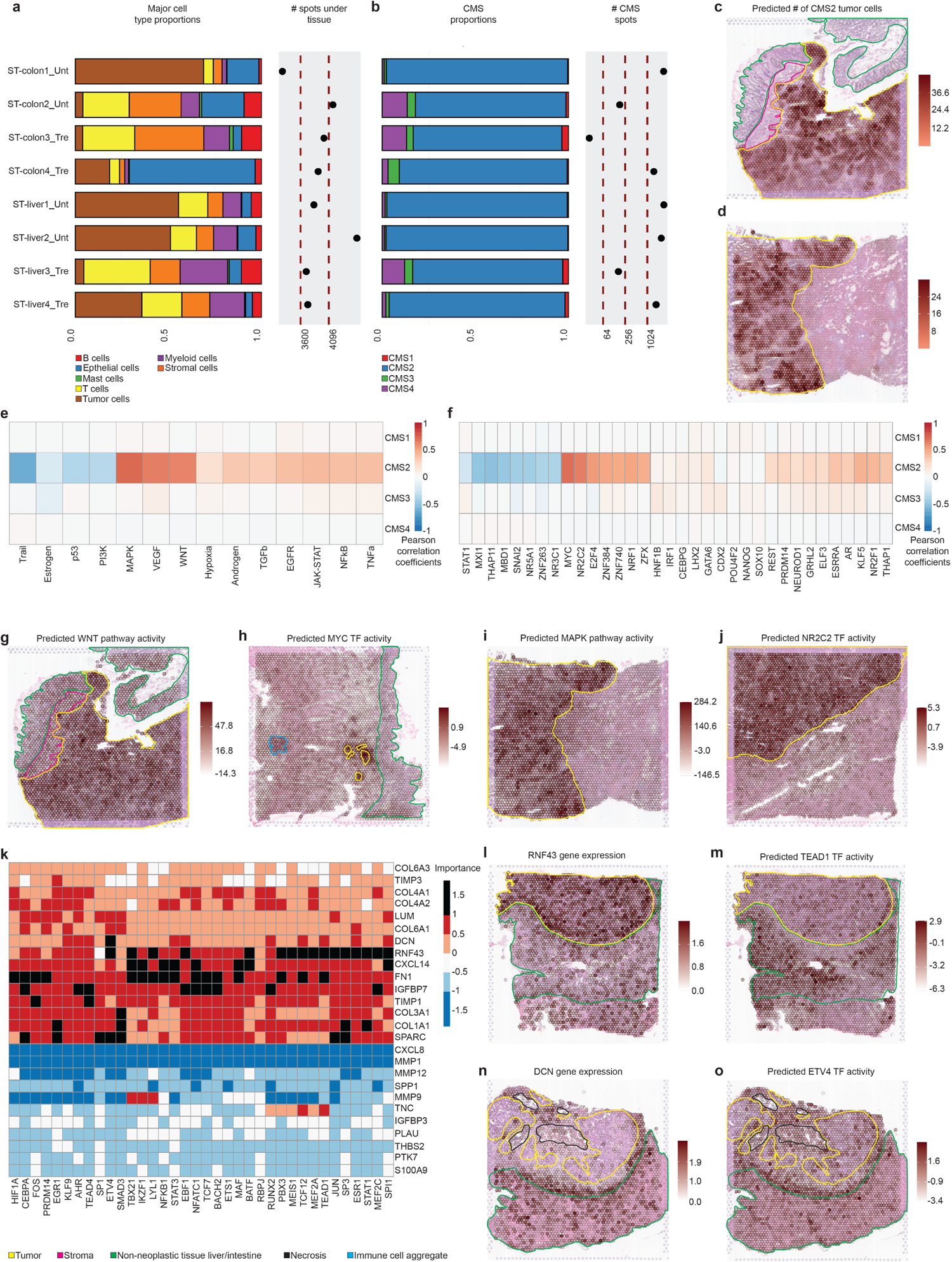
Characterization and analysis of an external ST CRC dataset to support the results in our internal set of samples. (a) Proportions of major cell classes per sample as estimated by the results of the deconvolution. The right hand side of the plot displays the number of analyzed spots per sample. (b) CMS tumor cell type proportions per sample as estimated by the results of the deconvolution approach. The number of spots containing an abundance of at least 20% of tumor cells subtypes is also displayed. (c-d) Overlay of the spatial mapping of the predicted CMS2 tumor cell abundance with the pathologists’s tissue annotations in the ST-colon1_Unt and ST-liver1_Unt samples. (e) Per spot Pearson’s cross-correlation across all the samples between pathway activities and CMS cell abundances. (f) Per spot Pearson’s cross-correlation across all the samples between TF activities and CMS cell abundances. For visualization purposes, the 10 most highly correlated TFs in absolute value per CMS are shown. (g) Overlay of the spatial mapping of the predicted WNT pathway activity with the pathologists’s tissue annotations in the ST-colon1_Unt sample. (h) Overlay of the spatial mapping of the predicted MYC TF activity with the pathologists’s tissue annotations in the ST-colon2_Unt sample. (i) Overlay of the spatial mapping of the predicted MAPK pathway activity with the pathologists’s tissue annotations in the ST-liver1_Unt sample. (j) Overlay of the spatial mapping of the predicted NR2C2 TF activity with the pathologists’s tissue annotations in the ST-liver2_Unt sample. (k) Misty results showing the potential importance of ligands (rows) expression on TF (columns) activity when considering the samples from primary CRC tumors. The ligand-TFs relationships with an importance score over 1 are represented as black slots and were considered as relevant. (l-m) Overlay of the spatial mapping of the RNF43 gene expression and the predicted TEAD1 TF activity with the pathologists’s tissue annotations in the ST-colon4_Tre sample. (n-o) Overlay of the spatial mapping of the DCN gene expression and the predicted ETV4 TF activity with the pathologists’s tissue annotations in the ST-liver4_Unt.

Next, we additionally classified these samples by applying CMScaller^24^ on pseudo-bulk RNA-seq generated by pooling together all their spots (Supplementary Fig. S54 and Methods). Interestingly, only ST-colon1-Unt, ST-liver1-Unt and ST-liver2-Unt were classified as CMS2. The colon samples with minor tumor content, ST-colon2-Unt and ST-colon3-Tre, were labeled as CMS4, suggesting that their stromal content was driving their classification. The liver sample with reduced tumor content, ST-liver3-Tre, was not assigned to any CMS. The ST-colon4-Tre and ST-liver4-Tre samples were respectively classified as CMS3 and CMS1, despite their large abundance of CMS2 tumor cells revealed by the deconvolution results. For ST-colon4-Tre, we hypothesized that its large content of non-neoplastic intestinal epithelium is driving the classification towards CMS3 (see Supplementary Fig. S45). The high content of resident macrophages in the liver tissue may be associated with the classification of ST-liver4-Tre as CMS1 (Supplementary Fig. S55). In summary, we emphasized again the lack of resolution of bulk-transcriptomics based classification systems to describe CRC heterogeneity.

We then spatially mapped the main CRC associated molecular features and examined their correlations with CMS cell abundance jointly considering primary and hepatic metastatic tumors (Figs. 6e-f and Methods). We focused our analysis on CMS2 given the limited number of cells estimated for the remaining subtypes. At the pathway level, we confirmed the activation of the WNT and VEGF pathways in CMS2 rich regions (Fig. 6g and Supplementary Fig. S56a). Regarding transcriptional activity, we corroborated the activation of the MYC and E2F4 TFs in CMS2 tumors (Fig. 6h and Supplementary Fig. S56b). Besides these well-known CMS2 features, we also found correlations between the number of estimated CMS2 cells and the activity of the MAPK pathway and the NR2C2 TF (Figs. 6i-j), in line with the findings in our set of samples (see Figs. 2k-l). These results are particularly interesting as their role in CMS2 tumors is not clearly defined.

We finally used this external dataset to corroborate selected cell-to-cell communication processes of those described above, i.e. the previously predicted ligand-TF regulations. For this purpose, we first computed Misty scores on the primary CRC tumors (Fig. 6k and Methods). The results supported the modulation of the transcriptional activity of JUN and members of the TEAD family by the expression of *RNF43* (Figs. 6l-m). Moreover, we captured again the potential influence of *DCN* expression on the transcriptional activity of ETV4 (Supplementary Figs. S57a-b). We also confirmed that the CXCL14 chemokine may potentially have a downstream impact in the transcriptional activity of MAF (Supplementary Figs. S57c-d). Next, we computed Misty scores on the liver metastatic tumors (Supplementary Fig. S58 and Methods). Our results indicate that the modulation of the transcriptional activity of ETV4 and JUN by DCN and RNF43, respectively, are preserved after metastasis (Figs. 6n-o and Supplementary Fig. S59). These findings are in accordance with and can provide new mechanistic insights to a recent study describing the protective role of DCN in hepatic metastasis of CRC^71^.

Overall, we confirmed the main results of our study in an independent ST CRC dataset. The deconvolution of these external samples validated the spatial molecular and morphological features of the CMS, particularly of CMS2. We also corroborated a part of the previously inferred cell communication processes involved in tumor progression. Moreover, we depicted the conservation of the CMS2 phenotype and of some ligand-TF regulations in CRC liver metastasis.

## 3. Discussion

The clinical need for accurate CRC patient stratification led to the development of several gene expression-based classification systems, such as the CMS^5^ or the CRC intrinsic subtypes (CRIS)^72^. The CMS classification system is broadly used and has helped to understand the different molecular mechanisms underlying CRC and disease prognosis^73^. Nevertheless, CMS intra-tumor heterogeneity hampers its clinical application, underlining the necessity of further characterizing the cellular composition and architecture of CRC and its microenvironment.

To complement our understanding of CRC CMS, we used ST combined with scRNA-seq through cell type deconvolution to delineate subtype inherent transcriptomic and morphological features. Our spatial alignment of CMS signatures with pathologists’ annotations, distinctly confined CMS1 and CMS2 cells to the neoplastic areas. Interestingly, the CMS1-CMS2 mixed S3_Col_R sample revealed a co-localization of CMS1-CMS2 cells indicating their coexistence. CMS1 signals were predominant in the diffuse-growing tumor, whereas CMS2 signatures were accentuated in a defined region showing a tubular growth pattern, in line with morphological features described by Thanki et al.^28^. These results stress the ability of ST to characterize mixed or transitioning CMS phenotypes and to reveal features that cannot be described using bulk- or scRNA-seq. In all the analyzed samples, CMS3 signatures were exclusively detected in the non-neoplastic mucosa, which can be associated with the previously described normal-like expression patterns of CMS3^5^. The EMT-associated CMS4 signals were minor, but overlapped with tumor regions displaying an invasive phenotype. Their limited abundance across the analyzed samples is in line with the very low number of tumor-like epithelial cells showing a CMS4 phenotype reported in previous publications^7^ and in our scRNA-seq reference^12^. Indeed, previous studies have proposed that CMS4 defines a transcriptional state or a stromal cell signature, rather than tumor cells^74^. Larger datasets are required to confirm whether ST can provide new insights into the nature of CMS4 tumors or can be used to delimitate the most invasive parts of other CRC subtypes.

The discrepancies in the CMS categorization between the deconvolution- and the bulk-based approaches, underline the large influence of adjacent tissue components on tumor classification. This is illustrated by both replicates of the S6_Rec patient, which were classified as CMS2 by the deconvolution and assigned to CMS4 by CMScaller^24^. Histologically, this sample is composed of small tumor islands surrounded by large stroma bundles. This morphology hampers the separation of the tumor components for bulk RNA sequencing, whereas ST can provide a detailed assessment of them. The CMS4 classification of stroma-rich tumors is in accordance with previous studies reporting that the CMS4 signature is highly influenced by marker genes of cancer-associated fibroblasts and other stromal cells^9, 25, 26^. The external ST-colon4_Tre sample, which was categorized as CMS2 by deconvolution and classified as CMS3 by CMScaller^24^, represents another example. This case raises concern about the potential influence of the non-neoplastic mucosa, containing CMS3 signals as described above, in bulk-based CMS classification systems.

Overall, our results underline the potential of ST in CRC characterization, enabling the spatial correlation of morphological tumor, stroma and non-neoplastic tissue patterns with corresponding transcriptomic features. Nevertheless, limitations inherent to our deconvolution-based approach should be acknowledged. The deconvolution partially failed to map stromal cells in the regions annotated as such by the pathologists. This effect is pronounced in the S3_Col_R sample and was observed regardless of the scRNA-seq dataset used as reference. This might be related to the lack of this specific stroma cell type in the scRNA-seq reference datasets. It can also correspond to a more general issue related with a potential loss of sensitivity in the deconvolution results in anatomical regions where the number of transcripts per spot was lower due to tissue inherent properties or technical and processing variabilities, as illustrated in Supplementary Figs. S60-S62. Another aspect of the deconvolution that requires critical consideration is the impact of the scRNA-seq dataset used as reference. In our study, we compared the results yielded by two reference datasets that were annotated using the same criteria^12^. As described in Supplementary Note 1, the overall deconvolution results were similar between references, but notable differences for particular cell types, such as CMS1, were observed. We hypothesized that the most comprehensive Korean dataset could lead to more accurate cell type specific signatures for deconvolution purposes. Additionally, the differences in the genetic background between both cohorts can contribute to these discrepancies.

We also explored the power of ST to explore inter- and intra-tumor heterogeneity and cell communication processes in CMS2 carcinomas. Regarding CMS2 heterogeneity, our results defined patient specific signaling cascades for the mTORC1 and EGFR pathway, and suggested specific features in stromal, immune-rich and tumor regions that might be relevant for personalized treatment approaches. For cell communication processes, we explored ligand-receptor interactions at the tumor-stroma interface potentially involved in tumor progression. To support our results and to identify the specific cell types involved in these processes, we estimated in parallel ligand-receptor interactions and TF activity on the scRNA-seq dataset from the Lee et al. publication^12^. By using this approach, we revealed signaling cascades modulating the interplay between CMS2 tumor cells and their TME (mesenchymal or vascular stroma components and immune cells), which are crucial for tumor progression and immune phenotyping. Interestingly, some of our results suggested a protective response mitigating tumor growth, such as the potential effect of *DCN* expression on the transcriptional activity of ETV4 through its binding to the EGFR receptor.. On the other hand, we inferred modulations that seem to promote tumor growth and invasiveness. For instance, myofibroblast cells secreted the PLAU ligand that was predicted to target the PLAUR receptor in SPP1^+^ macrophages, which are known to be associated with EMT in CRC^21, 75^. Overall, our results revealed several well characterized and novel cell-to-cell interactions, highlighting the potential of ST to delineate potential therapeutic targets for specific CMS subtypes. Nevertheless, the ligand-TF modulations predicted in this study need to be further investigated. They were inferred by modeling the spatial maps of TF activity using gene expression of ligands as a proxy. Therefore, some of the results may not represent direct causal regulation as a consequence of the complex network of cellular interactions and alternative signaling cascades in cancers. In a similar line, the ligand-receptor interaction analysis can also capture indirect gene expression associations, as may be the case for the predicted RNF43-FZD2 interaction, which is mostly reported as intracellular in the literature.

We finally supported our key findings using an independent ST CRC dataset comprising four CRC samples from primary tumors and their corresponding liver metastatic samples. Interestingly, our deconvolution approach delineated the primary, but also the metastatic carcinomas, as CMS2. In these liver tumors, we captured the CMS2 main molecular features, like the activation of the WNT pathway or high expression of MYC-regulated genes, and preserved cell communication events as the modulation of the transcriptional activity of ETV4 by DCN. This suggested that the CMS2 phenotype was recovered to some extent after migration of the primary CRC cells to sites of metastasis. Systematic and organ specific assessment of the active common pathways in primary and metastatic carcinomas might support treatment strategies for stage IV tumors.

In conclusion, our study highlights how ST, coupled with scRNA-seq, provides a novel dimension to explore patient conserved and specific molecular features of CRC and its CMS by characterizing the spatial arrangement of the different cell types composing tumors and their TME. We acknowledge that the limited number of samples and patients in our study hampers a comprehensive analysis of CRC heterogeneity. Moreover, we are working with 2-dimensional sections of tumors, whereas a 3-dimensional view is required to fully describe and characterize them. However, we envision that our proof-of-concept work delineates the potential of ST to contribute to patient-specific treatment approaches. On the one hand, more refined patient stratification strategies can be designed by taking into account the composition and spatial distribution of the cells composing the tumor and the TME, and its integration with morphological features extracted from the corresponding histological images. On the other hand, intra-tumor spatial heterogeneity can reveal tumor progression related processes that are anatomically restricted or more pronounced in a particular region. This can fuel the development of new treatment strategies, such as combinational therapies or spatially restricted medication, potentially leading to improvements in drug-efficacy and dosage reductions. In this regard, the foreseeable improvements in the resolution, reduction of processing costs per sample and clinical validation of ST technologies will facilitate the detailed analysis of larger CRC cohorts towards personalized oncology.

## 4. Methods

### 4.1 Collection of CRC samples

Human CRC tissues (<8 months storage) and annotated data were obtained and experimental procedures were performed within the framework of the non profit foundation HTCR, including the informed patient’s consent^76^. Tissues were cut on a Cryostat (CryoStar NX70, Thermo Scientific) at 10 um. Pathologists performed quality and comparability assessment of FF material using an H&E stained slide.

### 4.2 Sample preparation

RNA from all samples was extracted using the Arcturus® PicoPure® RNA Isolation Kit (Applied Biosystems™, KIT0204). For cell lysis, a 10 um section of the sample was resuspended in a 200 ul extraction buffer. Total RNA was extracted following the instructions of the manual. RNA integrity number (RIN) was assessed using the 2100 Bioanalyzer system (Agilent Technologies, Inc.) with a Agilent RNA 6000 Pico Kit (Agilent Technologies, Inc., 5067-1513). Samples with RIN above 7.0 were used.

Tissue optimization was carried out according to the manufacturer’s instructions (VISIUM Spatial Tissue Optimization User Guide_RevC). Image acquisition was performed on the Hamamatsu NanoZoomer S 360 C13220 series at 40x magnification and the coverslip was removed afterwards by immersing the slide in a 3x Saline-Sodium Citrate buffer. The stained tissue sections were permeabilized using a time course to test for the optimal permeabilization time. After performing a fluorescent cDNA synthesis, the tissue was removed. Finally the fluorescent cDNA was imaged using a Zeiss Axio Scan.Z1 with a Plan Apochromat 20x/0.8 M objective, a ET-Gold FISH filter (ex 538-551 nm/em 556-560 nm) and 100 ms exposure time.

For the gene expression analysis, 10 um thick sections of the samples were placed with a random distribution over four chilled 10x Genomics VISIUM Gene Expression slides containing four capture areas each. The sections were similarly stained with H&E and subsequently imaged as described above. To release the mRNA, the sections were permeabilized for 30 min as defined by tissue optimization. For further processing, the cDNA was amplified according to the manufacturer’s protocol (CG000239_VisiumSpatialGeneExpression_UserGuide_RevC). Double indexed libraries were prepared. The libraries were quality controlled using a 2100 Bioanalyzer system with Agilent High Sensitivity DNA Kit (Agilent Technologies, Inc., 5067-4626) and quantified with Qubit™ 1X dsDNA HS Assay Kit (Invitrogen, Q33230) on a Qubit 4 Fluorometer (Invitrogen, Q33238). The libraries were loaded onto the NovaSeq 6000 (Illumina) at a concentration of 250 pM. A NovaSeq S1 v 1.5 or SP v 1.5 Reagent Kit (100 cycles) (Illumina, 20028319 and 20028401) was used. For paired end-dual indexed sequencing, the following read protocol was used: read 1: 28 cycles; i7 index read: 10 cycles; i5 index read: 10 cycles; and read 2: 90 cycles. All libraries were sequenced at a minimum of 50000 reads per covered spot.

Raw sequencing data were demultiplexed using the *mkfastq* function from Space Ranger (v. 1.2.0). Demultiplexed data were mapped to the human reference GRCh38 with *spaceranger count*. Spots under tissue folds, artifacts and at the tissue boundary were manually removed using the 10X Loupe browser (v. 5.1.0).

### 4.3 Histopathological annotations and spot categorization

H&E stained tissue sections were annotated by the pathologists using Q-Path software (v. 0.2.3)^77^. Spot categorization was performed by the pathologists using the 10X Loupe browser (v. 5.1.0). Categories and corresponding criteria are listed in Supplementary table 6.

### 4.4 Grading of CMS signatures

Grading of CMS signatures in the tumor tissue was performed semi-quantitatively according to the number of spots with positive signature and the percentage of positive cells per spot. This grading was done in an individual replicate per patient (S1_Cec_Rep1, S2_Col_R_Rep1, S3_Col_R_Rep1, S4_Col_Sig_Rep1, S5_Rec_Rep1, S6_Rec_Rep2 and S7_Rec/Sig_Rep1) according to the scheme detailed in Supplementary table 2.

### 4.5 Bioinformatic analysis

#### 4.5.1 ST data pre-processing

We used the Seurat^78^, Scanpy^79^ and SingleCellExperiment^80^ packages to load the output of the Space Ranger pipeline and process the ST data. We evaluated the quality of the ST data by determining the average number of reads, UMIs and genes per spot covered by tissue and compared it with those from spots non covered by tissue. We found substandard quality for the S1_Cec_Rep2 sample as revealed by its low numbers of unique molecular identifier (UMI) counts and genes in spots covered by tissue (Supplementary Fig. S1). Consequently, this sample was either treated carefully or excluded from integrative analysis. For each individual sample, we filtered out spots for which the number of UMI counts detected were below 500 or above 45000. In addition, spots containing a fraction of more than 0.5 mitochondrial genes were not considered in the analysis. We normalized the UMI counts from the remaining spots using SCTransform^81^.

#### 4.5.2 Sample integration, batch correction and dimensionality reduction

To jointly represent the CRC samples in the same low dimensional space (UMAP embedding), correct from batch effects and integrate samples and technical replicates for downstream analysis, we used Harmony, which is a robust and efficient algorithm designed to integrate scRNA-seq datasets^82^. We ran Harmony with default parameters allowing a maximum number of 20 interactions (*max.iter.harmony = 20*) and correcting per individual samples. Of note, Harmony was either applied to batch-correct for all the spots derived from all the samples or to batch-correct only the tumor annotated spots from a subset of samples (CMS2 tumor samples).

#### 4.5.3 Deconvolution of the ST datasets

ST datasets derived from 10x Genomics VISIUM technology currently lack single cell resolution. Therefore, the gene expression values detected per spot originate from a variable number of different cells, i.e. every spot can be considered as a mini-bulk RNAseq dataset. Consequently, a deconvolution approach is required to estimate the different cell types and their proportions across spots.

To this end, we used the recently proposed Cell2Location (v 0.0.5)^23^ method. Cell2location first creates gene expression signatures of cell types from a scRNA-seq reference. We adopted as scRNA-Seq reference a comprehensive dataset from a recent publication exploring the cellular landscape of the different CRC subtypes and their microenvironment^12^. The annotations from the original publication at the cell subtype level (Supplementary Table 1) were used to generate the signature using the *run_regression* function with the following parameters: *n_epochs*=*100*, *minibatch_size*=*1024*, *learning_rate*=*0.01* and *train_proportion*=0.9. These signatures are subsequently used to assess cell type abundances in the ST data using the *run_cell2location* with *selection_specificity*=*0.20*. This parameter determines the number of genes used to establish the signature per cell type (Supplementary Table 1). Additional parameters were set as follows: *n_iter=40000, cells_per_spot=8, factors_per_spot=9, combs_per_spot: 5, mean= 1/2 and sd= 1/4*.

#### 4.5.4 Consistency of deconvolution results between technical replicates

To evaluate the consistency of the deconvolution between technical replicates, we batch-corrected their transcriptomic profiles using Harmony^82^ as described above. Then, we clustered the Harmony embeddings using the Louvain algorithm as encoded in the *FindClusters* function from the Seurat package. We chose a series of large resolution parameters (ranging from 1 to 2 increasing by 0.1 steps) to obtain fine-grain clusters that can match with anatomical regions displaying similar cell type distribution patterns across replicates. Finally, we computed the mean number of UMIs estimated by Cell2Location per cell type and cluster, and applied Pearson’s correlation to evaluate their similarity between technical replicates.

#### 4.5.5 Enrichment/depletion of cell types in different anatomical regions

The enrichment (depletion) in the abundance of the deconvolution-estimated cell types in different pathologist-assigned tissue categories was assessed following a similar procedure to be one described in Andersson et al.^16^. Briefly, the estimated cell type proportions per spot were 10 000 times randomly shuffled with respect to their spatial location. Then, we computed the average cell type proportions per permutation and tissue type. The mean value of differences between the real and the permuted average proportions divided by the standard deviation of these differences was used as the enrichment score for the different tissue categories.

#### 4.5.6 Cell-type colocalization analysis

Deconvolution results were subsequently used to assess cell-colocalization using non-negative matrix factorization as described in Cell2Location tutorials (https://cell2location.readthedocs.io/en/latest/notebooks/cell2location_tutorial.html#Identifying-cellular-compartments-/-tissue-zones-using-matrix-factorisation-(NMF)). As suggested by the authors, we examined results for different numbers of factors. We chose *R=3* and *R=7* as good representatives between strong co-localization signals and distinctive anatomical regions.

#### 4.5.7 Pathway activity

We estimated pathway activity per spot and at subspot resolution (see section 4.5.13) using PROGENy^83, 84^. PROGENy computes pathway activity by accounting for the expression of genes which are more responsive to perturbations on those pathways. The PROGENy model comprises 14 pathways, namely: Wnt, VEGF, Trail, TNFα, TGFβ, PI3K, p53, NFkB, MAPK, JAK/STAT, Hypoxia, Estrogen, Androgen and EGFR. In our setup, we ran PROGENy using the top 500 most responsive genes per pathway.

In addition, we also computed pathway activities in pseudo-bulk generated from our ST samples (see section 4.5.11). We again used the top 500 most responsive genes per pathway. In this case, we set the *scale* parameter to *TRUE* to allow direct comparison of pathway activities between samples.

#### 4.5.8 Transcription factor activity

We computed TF activity per spot using the Viper^85^ algorithm coupled with regulons extracted from DoRothEA^86^. In DoRothEA, every TF–target interaction is assigned a confidence score based on the reliability of its source, which ranges from A (most reliable) to E (least reliable). In this study, we selected interactions with confidence scores A, B and C and computed the activity for TFs with at least four different targets expressed per spot.

The activity profiles of the different TFs were additionally used to cluster the spots from our four CMS2 tumor samples. To do so, the TF activity scores from these samples were first merged and subsequently scaled and centered. Then, the standard procedure to compute clustering using the Seurat package was followed. Briefly, we computed a Principal Component Analysis (PCA) dimensionality reduction on the scaled TF activities per spot followed by the computation of the 20 nearest neighbors. Finally, we applied the Louvain algorithm with a resolution parameter of 0.5 to group the spots into different clusters according to their TF activity profile. We identified TF with a differential activity profile among the different clusters using Receiver Operating Characteristic (ROC) analysis as implemented in the Seurat’s *FindAllMarkers* function. We only considered TF whose activity was computed in at least 25% of the spots per cluster and with a log_2_ fold-change greater than 1. Of note, we used the same procedure to compute TF activity per cell on the scRNA-seq dataset from Lee et al.^12^.

#### 4.5.9 Canonical correlation analysis

We used the *cc* function from the CCA package^87^ to compute canonical correlation between the cell type proportions per spot and pathway or TF activity per spot. This canonical correlation analysis was first performed for every individual CRC sample. To capture global correlations across samples, we performed an integrative analysis by merging spots coming from all the different samples (excluding S1_Cec_Rep2) into matrices and computing the canonical correlation on them.

#### 4.5.10 Selection of tumor surrounding spots

We applied the *GetTissueCoordinates* function from the Seurat package to get the spatial coordinates of the spots in the different CRC samples. We subsequently computed the Euclidean distance between every pair of spots. Finally, we selected as tumor surrounding spots those lying within a distance smaller or equal to 2 from a tumor annotated spot. Spots fulfilling these criteria but annotated as tumors were removed.

#### 4.5.11 Pseudo-bulk generation

We generated pseudo-bulk from the ST samples using the *sumCountsAcrossCells* function from the Scater package^88^. Here, counts were normalized by the total number of reads (counts per million normalization). We used the *filterByExpr* function from the edgeR package^89^ to filter out genes with less than 50 counts per sample.

#### 4.5.12 Definition of different anatomical regions in tumor annotated spots

The distance between every tumor annotated spot and non-tumor annotated spots was calculated as described in section 4.5.10. We then defined the different tumor anatomical regions for the S2_Col_R_Rep1 sample based on the following criteria:

- *Peripheral Tumor*: tumor spots in direct contact with at least a non-tumor annotated spot. Their Euclidean distance to a non-tumor annotated spot is smaller than 2.
- *Central Tumor*: tumor spots in the most solid and internal region of the tumor. Their Euclidean distance to a non-tumor annotated spot is greater than 2.5.
- *Intermediary Tumor*: tumor spots that we consider as a transition region between the inner and outer tumor. Their Euclidean distance to a non-tumor annotated spot is greater or equal to 2 and smaller than 2.5.

#### 4.5.13 Clustering and enhanced gene expression at the sub spot level

We applied BayesSpace^90^ to cluster at the subspot level and increase the gene expression resolution of our CMS2 tumor annotated spots in the S5_Rec_Rep1 sample. To do so, BayesSpace uses the neighborhood structure in spatial transcriptomic data. Of note, the preprocessing of the ST raw data was conducted following the recommendations of BayesSpace authors. This procedure is slightly different from the one described in previous sections. Briefly, the ST data was processed using the SingleCellExperiment package and raw counts were log normalized using the *logNormCounts* function from the Scuttle package^88^. Then, the Scran^91^ package was used to model the variance of the log-expression profiles for each gene and select the 2000 most variable genes. We performed a PCA using the Scater ^88^ package.

Using BayesSpace, we subsequently computed the spatial clustering and the enhanced clustering with default parameters, excepting the *jitter_scale* parameter which was set to 3. Finally, we enhanced the gene expression of all the genes expressed in the considered spots using the *enhanceFeatures* function with default parameters.

#### 4.5.14 Differential gene expression analysis

The CMS2 tumor regions extracted from the different samples were integrated into the same Seurat^78^ object. We used the Wilcoxon Rank Sum test to identify differentially expressed genes between the groups of spots coming from different patients as implemented in the Seurat’s *FindAllMarkers* function. We set a log_2_ fold-change threshold of 0.25 and only positive markers were retrieved. Some specific criteria were followed for the analyses conducted in section 2.3:

- To describe inter-patient heterogeneity, the differential gene expression analysis was performed between the different patients (two replicates per patient considered). We filtered results by only considering genes that are overexpressed in tumor annotated spots versus non-tumor annotated spots. To do so, we took advantage of the pathologists’ annotations and used the Seurat’s *FindMarkers* with the same parameters described above for the *FindAllMarkers* function. Ribosomal and mitochondrial genes were removed due to the fact that they can be overrepresented in tumor necrotic regions.
- To describe intra-tumor heterogeneity, the differential expression analysis was carried out between the different anatomical regions of the tumor in the S2_Col_R_Rep1 sample (see section 4.5.12) with no further considerations.
- Another differential gene expression analysis was conducted on the enhanced gene expression between the different enhanced clusters generated by BayesSpace (see section 4.5.13) on the S5_Rec_Rep1 sample. We selected for further analysis genes with an adjusted p-value smaller than 0.01 in the Wilcoxon Rank Sum test. Ribosomal and mitochondrial genes were excluded from the analysis.

#### 4.5.15 Gene set overrepresentation analysis

Differentially expressed genes were subsequently used for gene set overrepresentation analysis using the Hallmark annotations from MSigDB^92^. The Hallmark gene sets contain 50 well-defined biological states or processes. We used the *enricher* function from the clusterProfiler^93^ package to carry out the analysis. We set a minimal size of the genes annotated for testing to five, excepting for the analysis between different patients where it was set to three. Background genes were adjusted accordingly to the global set of genes expressed in the different contexts.

#### 4.5.16 Ligand modulation of TF activity

As a first step and taking as reference the TF activity-based clustering, we selected ligands which are overexpressed in the tumor and TME with respect to the other anatomical regions across all our CRC samples. To do so, we applied the Seurat’s *FindMarkers* function with a log_2_ fold-change threshold of 0.5 and only positive markers were retrieved. We matched our set of overexpressed genes against the set of proteins annotated as ligands in the Omnipath^94^ database. Additionally, we filtered out ligands that are not detected in at least 10% of the tumor and TME spots in every individual sample.

In the second place, we chose TFs with a higher differential activity profile in the TME regions across all the samples according to the clustering approach described in section 4.5.8. In particular, we selected those TFs that are considered as markers of the TME cluster when using the Seurat’s *FindAllMarkers* function (AUC ≥ 0.75).

We then applied Misty^48^ to investigate the potential effect of the expression of the selected ligands in modulating the transcriptional activity of the chosen TFs. Specifically, we created an intrinsic view (*intraview*) describing ligand gene expression and a local niche view (*juxtaview*) using TF activity with a *neighbor.thr =2* aiming at capturing effects in the direct neighborhood of each spot. This criteria is based on the fact that many cancer relevant ligands are membrane bound and that the majority of secreted ligands cannot travel long distances. Following this approach, Misty was first individually applied to every sample. Then, the individual results were collected and aggregated using Misty’s *collect_results* function in order to obtain the most robust common signals across samples. Ligand-TF associations with an aggregated importance greater than 1 were considered for further analysis. Of note, when running Misty on the external dataset, the ST-colon3-Tre and ST-liver3-Tre samples were excluded from the analysis due to their very reduced tumor content.

#### 4.5.17 Prediction of Ligand-Receptor interactions

We used LIANA^95^ to estimate the most likely ligand-receptor interactions between the different spatial clusters defined by their TF activity profiles. It is to note that the interactions were computed for every pair of clusters, but for subsequent analysis and visualization we focused on the interactions between the clusters labeled as 0 (Tumor) and 1 (TME). LIANA computes an aggregated score for every potential ligand-receptor interaction based on the results of different methods. In our particular case, we ran LIANA with default settings and used OmniPath^94^ as a source of prior knowledge in human ligand-receptor interactions. For further analysis, we considered interactions involving Misty’s predicted ligands with an aggregated rank smaller than 0.01, as this value can be seen as analogous to a p-value ^96^. We also ran LIANA on the scRNA-seq dataset from Lee et al.^12^ using the same procedure.

#### 4.5.18 Inference of signaling networks

We used a network-based approach to infer the most likely signaling cascades linking LIANA’s predicted ligand-receptor interactions to their targeted TFs according to Misty’s predictions. To do so, we first built an intra-cellular signaling network by retrieving protein-protein interactions from Omnipath^94^. Then, for every ligand, we selected their predicted receptors and targeted TFs. We subsequently connected every receptor to every corresponding TF by selecting the shortest path between them in the signaling network. All the resultant shortest paths were merged into a network together with the previously predicted ligand-receptor interactions. Finally, for every gene in the predicted network, we computed its average expression in the TME cluster, as defined by TF activity profiles (see section 4.5.8), across all the CMS2 samples. Cytoscape^97^ was utilized for the visualization of the network.

#### 4.5.19 Metastasis Score

We retrieved a list of genes linked to metastatic processes from CancerSEA^98^ and computed their score per spot using the Seurat’s *AddModuleScore* function. We set the *ctrl* parameter, i.e. number of control features selected from the same bin per analyzed feature, to 20.

## 5. Data availability

The output of Space Ranger, including processed count data matrices and histological images, for the ST data generated in this study is available at https://doi.org/10.5281/zenodo.7551712. In addition, this repository also contains the spot categorization made by the pathologists. The processed scRNA-seq and metadata used for the deconvolution and for further characterization of the cell communication processes are available via the GEO database under the accession codes <u>GSE132465</u> and <u>GSE144735</u>^12^. The processed data from the external ST CRC dataset used to support our findings was downloaded from http://www.cancerdiversity.asia/scCRLM19.

## 6. Code availability

The scripts containing all the code used to generate the results presented in this study are available at https://github.com/alberto-valdeolivas/ST_CRC_CMS. Their associated notebooks containing additional results and information about the versions of the different packages used are available at https://doi.org/10.5281/zenodo.7440182. Finally, Intermediary object files to reproduce the analysis are available at https://doi.org/10.5281/zenodo.7551712.

## Acknowledgements

This work was supported by The Roche Postdoctoral Fellowship (RPF) programme. We thank Daniel Dimitrov, Ricardo Omar Ramirez Flores and Dario Zimmerli for productive scientific discussions around the topics covered in this manuscript.

## 7. Conflict of interests

AV, BA, EG, NG, MR, SB, IW, LV, EY, MB, MS, NK, BJ, PS, TB and KH are currently employed by F. Hoffmann-La Roche Ltd. AJL and DT were previously employed by F. Hoffmann-La Roche Ltd. AJL is currently employed by Idorsia Pharmaceuticals Ltd. DT is currently employed by University of Bern. AL is currently employed by Genentech, Inc. MDT was previously employed by Genentech, Inc and is currently employed by Gilead Sciences, Inc. JSR has received funding from GSK and Sanofi and fees from Travere Therapeutics and Astex Pharmaceuticals. The authors declare that they have no other competing interests.

## 8. Authors contributions

AV, KH, TB, BJ and PS planned and designed the study. AV, BA and KH wrote the manuscript with the input and feedback from the remaining authors. BA, NG, MR and NK conducted the sample preparation and laboratory experiments. AV, AJL and EG carried out the data analysis. KH, AL and MDT performed pathology assessments and assisted bioinformatics data interpretation. DT, LV, SB, IW, EY, MDT, MB, SR, JSR and MS provided guidance on the data analysis direction and the biological findings. All authors read and approved the final manuscript.

## Notes

### Summary of Updates

We have introduced some minor changes into the original version as a result of feedback upon publication of the first version. In particular, we have detailed that one of the ligand-receptor interactions described in the first version may actually correspond to an intra-cellular effect rather than a cell communication effect.

## References

1. Biller, L. H. & Schrag, D. Diagnosis and Treatment of Metastatic Colorectal Cancer: A Review. JAMA 325, 669–685 (2021).

2. Wang, W. et al. Molecular subtyping of colorectal cancer: Recent progress, new challenges and emerging opportunities. Semin. Cancer Biol. 55, 37–52 (2019).

3. Okita, A. et al. Consensus molecular subtypes classification of colorectal cancer as a predictive factor for chemotherapeutic efficacy against metastatic colorectal cancer. Oncotarget 9, 18698–18711 (2018).

4. Chan, D. K. H. & Buczacki, S. J. A. Tumour heterogeneity and evolutionary dynamics in colorectal cancer. Oncogenesis 10, 1–9 (2021).

5. Guinney, J. et al. The consensus molecular subtypes of colorectal cancer. Nat. Med. 21, 1350–1356 (2015).

6. Koulis, C. et al. Personalized Medicine-Current and Emerging Predictive and Prognostic Biomarkers in Colorectal Cancer. Cancers 12, (2020).

7. Chowdhury, S. et al. Implications of Intratumor Heterogeneity on Consensus Molecular Subtype (CMS) in Colorectal Cancer. Cancers 13, (2021).

8. Mooi, J. K. et al. The prognostic impact of consensus molecular subtypes (CMS) and its predictive effects for bevacizumab benefit in metastatic colorectal cancer: molecular analysis of the AGITG MAX clinical trial. Ann. Oncol. 29, 2240–2246 (2018).

9. Khaliq, A. M. et al. Refining colorectal cancer classification and clinical stratification through a single-cell atlas. Genome Biol. 23, 1–30 (2022).

10. Sirinukunwattana, K. et al. Image-based consensus molecular subtype (imCMS) classification of colorectal cancer using deep learning. Gut 70, 544–554 (2021).

11. Househam, J. et al. Phenotypic plasticity and genetic control in colorectal cancer evolution. Nature 611, 744–753 (2022).

12. Lee, H.-O. et al. Lineage-dependent gene expression programs influence the immune landscape of colorectal cancer. Nat. Genet. 52, 594–603 (2020).

13. Joanito, I. et al. Single-cell and bulk transcriptome sequencing identifies two epithelial tumor cell states and refines the consensus molecular classification of colorectal cancer. Nat. Genet. 54, 963–975 (2022).

14. Cañellas-Socias, A. et al. Metastatic recurrence in colorectal cancer arises from residual EMP1^+^ cells. Nature 611, 603–613 (2022).

15. Rao, A., Barkley, D., França, G. S. & Yanai, I. Exploring tissue architecture using spatial transcriptomics. Nature 596, 211–220 (2021).

16. Andersson, A. et al. Spatial deconvolution of HER2-positive breast cancer delineates tumor-associated cell type interactions. Nat. Commun. 12, 1–14 (2021).

17. Berglund, E. et al. Spatial maps of prostate cancer transcriptomes reveal an unexplored landscape of heterogeneity. Nat. Commun. 9, 1–13 (2018).

18. Hunter, M. V., Moncada, R., Weiss, J. M., Yanai, I. & White, R. M. Spatially resolved transcriptomics reveals the architecture of the tumor-microenvironment interface. Nat. Commun. 12, 1–16 (2021).

19. Wu, Y. et al. Spatiotemporal Immune Landscape of Colorectal Cancer Liver Metastasis at Single-Cell Level. Cancer Discov. 12, 134–153 (2022).

20. Peng, Z., Ye, M., Ding, H., Feng, Z. & Hu, K. Spatial transcriptomics atlas reveals the crosstalk between cancer-associated fibroblasts and tumor microenvironment components in colorectal cancer. J. Transl. Med. 20, 302 (2022).

21. Qi, J. et al. Single-cell and spatial analysis reveal interaction of FAP fibroblasts and SPP1 macrophages in colorectal cancer. Nat. Commun. 13, 1742 (2022).

22. Zhang, R. et al. Spatial transcriptome unveils a discontinuous inflammatory pattern in proficient mismatch repair colorectal adenocarcinoma. Fundamental Research (2022) doi:10.1016/j.fmre.2022.01.036.

23. Kleshchevnikov, V. et al. Cell2location maps fine-grained cell types in spatial transcriptomics. Nat. Biotechnol. (2022) doi:10.1038/s41587-021-01139-4.

24. Eide, P. W., Bruun, J., Lothe, R. A. & Sveen, A. CMScaller: an R package for consensus molecular subtyping of colorectal cancer pre-clinical models. Sci. Rep. 7, 1–8 (2017).

25. Herrera, M. et al. Cancer-associated fibroblast-derived gene signatures determine prognosis in colon cancer patients. Mol. Cancer 20, 73 (2021).

26. Isella, C. et al. Stromal contribution to the colorectal cancer transcriptome. Nat. Genet. 47, 312–319 (2015).

27. Ohue, Y. & Nishikawa, H. Regulatory T (Treg) cells in cancer: Can Treg cells be a new therapeutic target? Cancer Sci. 110, 2080–2089 (2019).

28. Thanki, K. et al. Consensus Molecular Subtypes of Colorectal Cancer and their Clinical Implications. Int Biol Biomed J 3, 105–111 (2017).

29. Mevizou, R., Sirvent, A. & Roche, S. Control of Tyrosine Kinase Signalling by Small Adaptors in Colorectal Cancer. Cancers 11, (2019).

30. Nunez, S. K. et al. Identification of Gene Co-Expression Networks Associated with Consensus Molecular Subtype-1 of Colorectal Cancer. Cancers 13, (2021).

31. García-Aranda, M. & Redondo, M. Targeting Receptor Kinases in Colorectal Cancer. Cancers 11, (2019).

32. Rebersek, M. Consensus molecular subtypes (CMS) in metastatic colorectal cancer - personalized medicine decision. Radiol. Oncol. 54, 272–277 (2020).

33. Bhawe, K. & Roy, D. Interplay between NRF1, E2F4 and MYC transcription factors regulating common target genes contributes to cancer development and progression. Cell. Oncol. 41, 465–484 (2018).

34. Orouji, E. et al. Chromatin state dynamics confers specific therapeutic strategies in enhancer subtypes of colorectal cancer. Gut 71, 938–949 (2022).

35. Francipane, M. G. & Lagasse, E. mTOR pathway in colorectal cancer: an update. Oncotarget 5, 49–66 (2014).

36. Martin, T. A. et al. NUPR1 and its potential role in cancer and pathological conditions (Review). Int. J. Oncol. 58, (2021).

37. Albuquerque, C. & Pebre Pereira, L. Wnt Signalling-Targeted Therapy in the CMS2 Tumour Subtype: A New Paradigm in CRC Treatment? Adv. Exp. Med. Biol. 1110, 75–100 (2018).

38. Shi, X., Young, C. D., Zhou, H. & Wang, X. Transforming Growth Factor-β Signaling in Fibrotic Diseases and Cancer-Associated Fibroblasts. Biomolecules 10, (2020).

39. Lin, Y., Xu, J. & Lan, H. Tumor-associated macrophages in tumor metastasis: biological roles and clinical therapeutic applications. J. Hematol. Oncol. 12, 76 (2019).

40. Brabletz, T. et al. Invasion and Metastasis in Colorectal Cancer: Epithelial-Mesenchymal Transition, Mesenchymal-Epithelial Transition, Stem Cells and β-Catenin. CTO 179, 56–65 (2005).

41. Naito, T. et al. Mesenchymal stem cells induce tumor stroma formation and epithelial-mesenchymal transition through SPARC expression in colorectal cancer. Oncol. Rep. 45, (2021).

42. Hapke, R. Y. & Haake, S. M. Hypoxia-induced epithelial to mesenchymal transition in cancer. Cancer Lett. 487, 10–20 (2020).

43. Kikuchi, K. & Tsukamoto, H. Stearoyl-CoA desaturase and tumorigenesis. Chem. Biol. Interact. 316, 108917 (2020).

44. Ran, H. et al. Stearoyl-CoA desaturase-1 promotes colorectal cancer metastasis in response to glucose by suppressing PTEN. J. Exp. Clin. Cancer Res. 37, 54 (2018).

45. Syed, V. TGF-β Signaling in Cancer. J. Cell. Biochem. 117, 1279–1287 (2016).

46. Wang, H., Birkenbach, M. & Hart, J. Expression of Jun family members in human colorectal adenocarcinoma. Carcinogenesis 21, 1313–1317 (2000).

47. Dittmer, J. The Biology of the Ets1 Proto-Oncogene. Mol. Cancer 2, 1–21 (2003).

48. Tanevski, J., Flores, R. O. R., Gabor, A., Schapiro, D. & Saez-Rodriguez, J. Explainable multiview framework for dissecting spatial relationships from highly multiplexed data. Genome Biol. 23, 97 (2022).

49. Deves, C. et al. Analysis of select members of the E26 (ETS) transcription factors family in colorectal cancer. Virchows Arch. 458, 421–430 (2011).

50. Keld, R. et al. The ERK MAP kinase-PEA3/ETV4-MMP-1 axis is operative in oesophageal adenocarcinoma. Mol. Cancer 9, 1–14 (2010).

51. Bi, X.-L. & Yang, W. Biological functions of decorin in cancer. Chin. J. Cancer 32, 266–269 (2013).

52. Gİrgİn, B., KaradaĞ-Alpaslan, M. & KocabaŞ, F. Oncogenic and tumor suppressor function of MEIS and associated factors. Turk. J. Biol. 44, 328–355 (2020).

53. Chen, J., Elfiky, A., Han, M., Chen, C. & Saif, M. W. The role of Src in colon cancer and its therapeutic implications. Clin. Colorectal Cancer 13, 5–13 (2014).

54. Du, B., Gao, W., Qin, Y., Zhong, J. & Zhang, Z. Study on the role of transcription factor SPI1 in the development of glioma. Chin Neurosurg J 8, 7 (2022).

55. Quesnelle, K. M., Boehm, A. L. & Grandis, J. R. STAT-mediated EGFR signaling in cancer. J. Cell. Biochem. 102, 311–319 (2007).

56. Diehl, V. et al. The Role of Decorin and Biglycan Signaling in Tumorigenesis. Front. Oncol. 0, (2021).

57. Jiang, X. et al. Inactivating mutations of RNF43 confer Wnt dependency in pancreatic ductal adenocarcinoma. Proc. Natl. Acad. Sci. U. S. A. 110, 12649–12654 (2013).

58. Koo, B.-K. et al. Tumour suppressor RNF43 is a stem-cell E3 ligase that induces endocytosis of Wnt receptors. Nature 488, 665–669 (2012).

59. Neumeyer, V. et al. Loss of endogenous RNF43 function enhances proliferation and tumour growth of intestinal and gastric cells. Carcinogenesis 40, 551–559 (2019).

60. Nie, X., Liu, H., Liu, L., Wang, Y.-D. & Chen, W.-D. Emerging Roles of Wnt Ligands in Human Colorectal Cancer. Front. Oncol. 10, 1341 (2020).

61. Zhao, B. et al. TEAD mediates YAP-dependent gene induction and growth control. Genes Dev. 22, 1962–1971 (2008).

62. Jiang, L., Li, J., Zhang, C., Shang, Y. & Lin, J. YAP-mediated crosstalk between the Wnt and Hippo signaling pathways (Review). Mol. Med. Rep. 22, 4101–4106 (2020).

63. Zhong, Z. A., Michalski, M. N., Stevens, P. D., Sall, E. A. & Williams, B. O. Regulation of Wnt receptor activity: Implications for therapeutic development in colon cancer. J. Biol. Chem. 296, 100782 (2021).

64. Tsukiyama, T. et al. Molecular Role of RNF43 in Canonical and Noncanonical Wnt Signaling. Mol. Cell. Biol. 35, 2007–2023 (2015).

65. Koch, M. et al. CD36-mediated activation of endothelial cell apoptosis by an N-terminal recombinant fragment of thrombospondin-2 inhibits breast cancer growth and metastasis in vivo. Breast Cancer Res. Treat. 128, 337–346 (2011).

66. Page-McCaw, A., Ewald, A. J. & Werb, Z. Matrix metalloproteinases and the regulation of tissue remodelling. Nat. Rev. Mol. Cell Biol. 8, 221–233 (2007).

67. Madunić, J. The Urokinase Plasminogen Activator System in Human Cancers: An Overview of Its Prognostic and Predictive Role. Thromb. Haemost. 118, 2020–2036 (2018).

68. Zhang, J., Sud, S., Mizutani, K., Gyetko, M. R. & Pienta, K. J. Activation of urokinase plasminogen activator and its receptor axis is essential for macrophage infiltration in a prostate cancer mouse model. Neoplasia 13, 23–30 (2011).

69. Liu, M. et al. Transcription factor c-Maf is a checkpoint that programs macrophages in lung cancer. J. Clin. Invest. 130, 2081–2096 (2020).

70. Hara, T. & Tanegashima, K. CXCL14 antagonizes the CXCL12-CXCR4 signaling axis. Biomol. Concepts 5, 167–173 (2014).

71. Reszegi, A. et al. The Protective Role of Decorin in Hepatic Metastasis of Colorectal Carcinoma. Biomolecules 10, (2020).

72. Isella, C. et al. Selective analysis of cancer-cell intrinsic transcriptional traits defines novel clinically relevant subtypes of colorectal cancer. Nat. Commun. 8, 15107 (2017).

73. Fontana, E., Eason, K., Cervantes, A., Salazar, R. & Sadanandam, A. Context matters-consensus molecular subtypes of colorectal cancer as biomarkers for clinical trials. Ann. Oncol. 30, 520–527 (2019).

74. Dunne, P. D. et al. Challenging the Cancer Molecular Stratification Dogma: Intratumoral Heterogeneity Undermines Consensus Molecular Subtypes and Potential Diagnostic Value in Colorectal Cancer. Clin. Cancer Res. 22, 4095–4104 (2016).

75. Georgoudaki, A.-M. et al. Reprogramming Tumor-Associated Macrophages by Antibody Targeting Inhibits Cancer Progression and Metastasis. Cell Rep. 15, 2000–2011 (2016).

76. Thasler, W. E. et al. Charitable State-Controlled Foundation Human Tissue and Cell Research: Ethic and Legal Aspects in the Supply of Surgically Removed Human Tissue For Research in the Academic and Commercial Sector in Germany. Cell Tissue Bank. 4, 49–56 (2003).

77. Bankhead, P. et al. QuPath: Open source software for digital pathology image analysis. Sci. Rep. 7, 1–7 (2017).

78. Hao, Y. et al. Integrated analysis of multimodal single-cell data. Cell 184, 3573–3587.e29 (2021).

79. Wolf, F. A., Angerer, P. & Theis, F. J. SCANPY: large-scale single-cell gene expression data analysis. Genome Biol. 19, 15 (2018).

80. Amezquita, R. A. et al. Orchestrating single-cell analysis with Bioconductor. Nat. Methods 17, 137–145 (2020).

81. Hafemeister, C. & Satija, R. Normalization and variance stabilization of single-cell RNA-seq data using regularized negative binomial regression. Genome Biol. 20, 296 (2019).

82. Korsunsky, I. et al. Fast, sensitive and accurate integration of single-cell data with Harmony. Nat. Methods 16, 1289–1296 (2019).

83. Schubert, M. et al. Perturbation-response genes reveal signaling footprints in cancer gene expression. Nat. Commun. 9, 20 (2018).

84. Holland, C. H. et al. Robustness and applicability of transcription factor and pathway analysis tools on single-cell RNA-seq data. Genome Biol. 21, 36 (2020).

85. Alvarez, M. J. et al. Functional characterization of somatic mutations in cancer using network-based inference of protein activity. Nat. Genet. 48, 838–847 (2016).

86. Garcia-Alonso, L., Holland, C. H., Ibrahim, M. M., Turei, D. & Saez-Rodriguez, J. Benchmark and integration of resources for the estimation of human transcription factor activities. Genome Res. 29, 1363–1375 (2019).

87. Gonzalez, I., Déjean, S., Martin, P. & Baccini, A. CCA: AnRPackage to extend canonical correlation analysis. J. Stat. Softw. 23, (2008).

88. McCarthy, D. J., Campbell, K. R., Lun, A. T. L. & Wills, Q. F. Scater: pre-processing, quality control, normalization and visualization of single-cell RNA-seq data in R. Bioinformatics 33, 1179–1186 (2017).

89. Robinson, M. D., McCarthy, D. J. & Smyth, G. K. edgeR: a Bioconductor package for differential expression analysis of digital gene expression data. Bioinformatics 26, 139–140 (2010).

90. Zhao, E. et al. Spatial transcriptomics at subspot resolution with BayesSpace. Nat. Biotechnol. (2021) doi:10.1038/s41587-021-00935-2.

91. Lun, A. T. L., McCarthy, D. J. & Marioni, J. C. A step-by-step workflow for low-level analysis of single-cell RNA-seq data with Bioconductor. F1000Res. 5, 2122 (2016).

92. Liberzon, A. et al. The Molecular Signatures Database (MSigDB) hallmark gene set collection. Cell Syst 1, 417–425 (2015).

93. Wu, T. et al. clusterProfiler 4.0: A universal enrichment tool for interpreting omics data. The Innovation vol. 2 100141 Preprint at https://doi.org/10.1016/j.xinn.2021.100141 (2021).

94. Türei, D. et al. Integrated intra- and intercellular signaling knowledge for multicellular omics analysis. Mol. Syst. Biol. 17, e9923 (2021).

95. Dimitrov, D. et al. Comparison of methods and resources for cell-cell communication inference from single-cell RNA-Seq data. Nat. Commun. 13, 1–13 (2022).

96. Kolde, R., Laur, S., Adler, P. & Vilo, J. Robust rank aggregation for gene list integration and meta-analysis. Bioinformatics 28, 573–580 (2012).

97. Shannon, P. et al. Cytoscape: a software environment for integrated models of biomolecular interaction networks. Genome Res. 13, 2498–2504 (2003).

98. Yuan, H. et al. CancerSEA: a cancer single-cell state atlas. Nucleic Acids Res. 47, D900–D908 (2019).

