## Supplementary Materials for "Charting the Heterogeneity of Colorectal Cancer Consensus Molecular Subtypes using Spatial Transcriptomics"

Supplementary figures and tables for the "*Charting the Heterogeneity of Colorectal Cancer Consensus Molecular Subtypes using Spatial Transcriptomics*" publication.

#### 1. Supplementary Figures

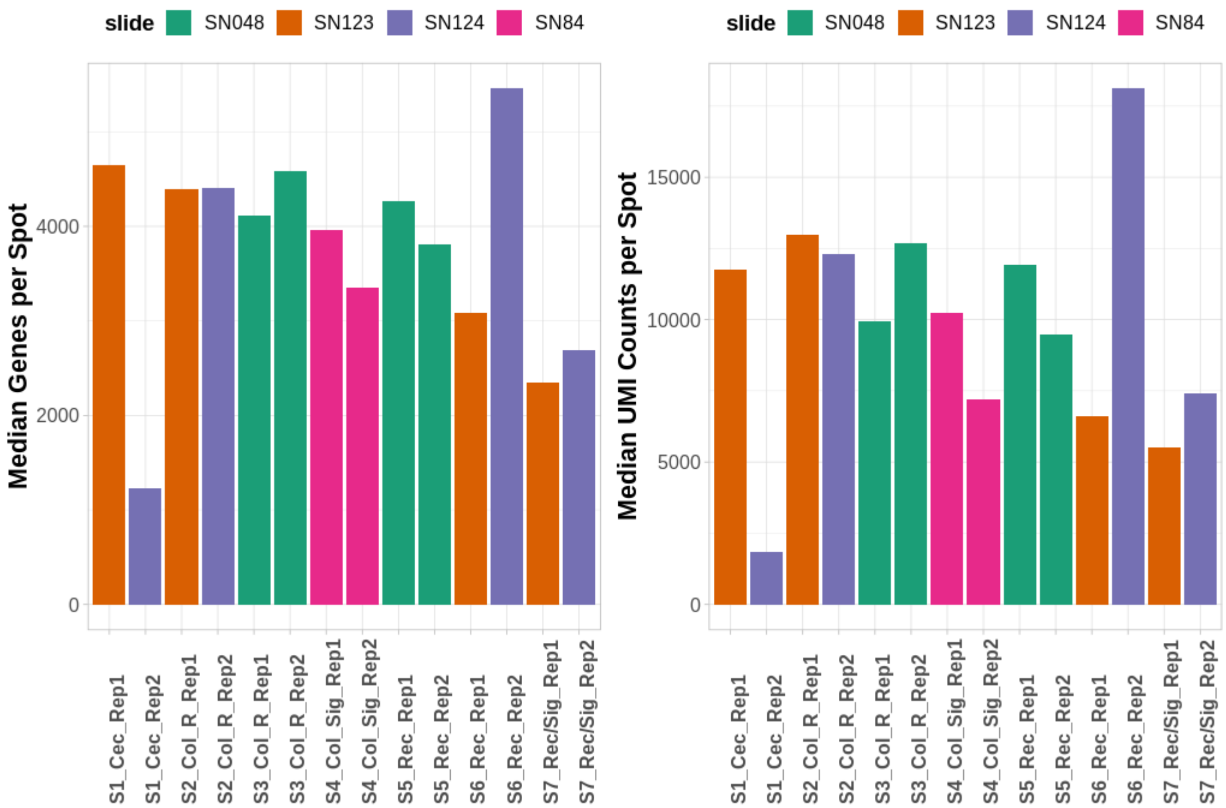


***Supplementary Figure S1:*** *Median Number of Genes and UMI counts per spot in our set of CRC samples. The bar color indicates the samples that were processed on the same gene expression slide. Note the reduced number of genes and UMI counts in the S1_Cec_Rep2 sample, which was considered of substandard quality.*

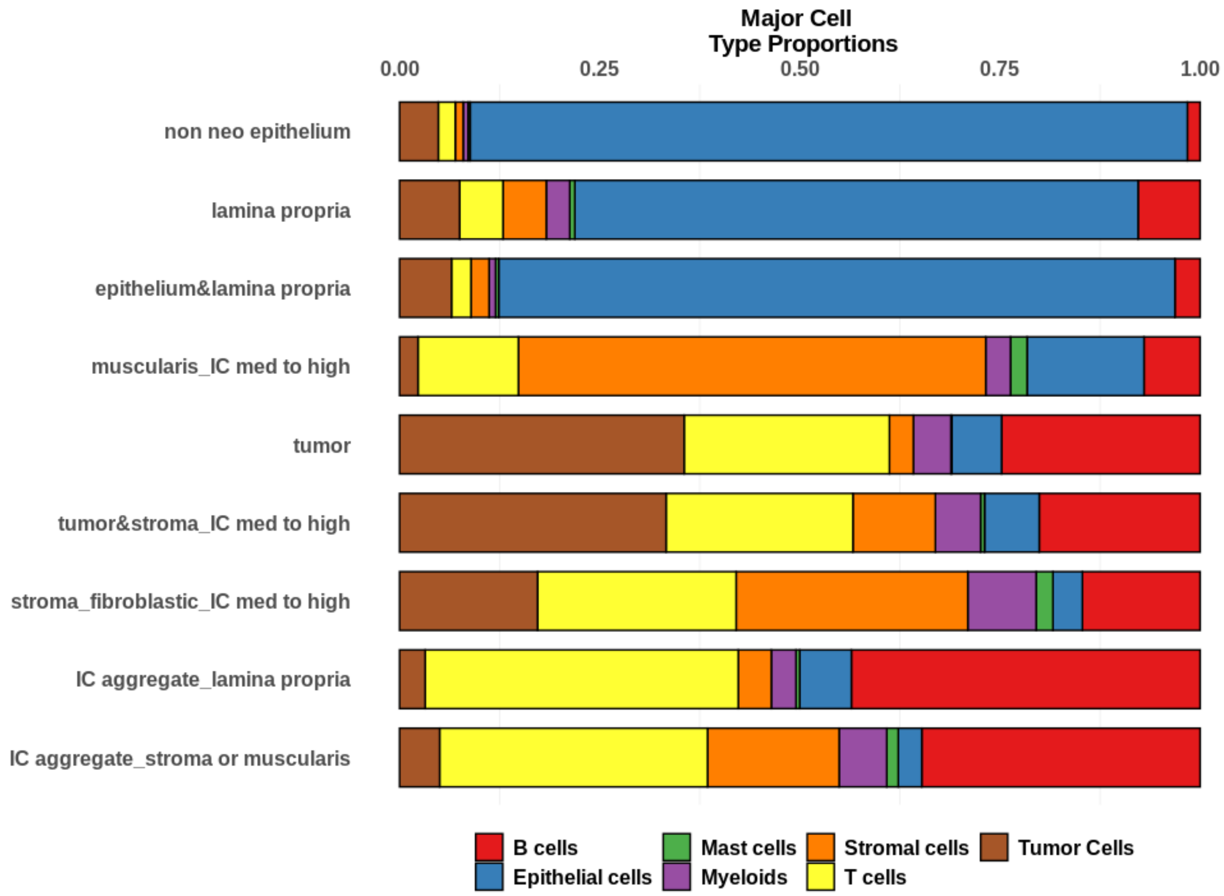


***Supplementary Figure S2:*** *Proportions of major cell classes as estimated by the results of the deconvolution approach in the different anatomical regions annotated by the pathologists.*

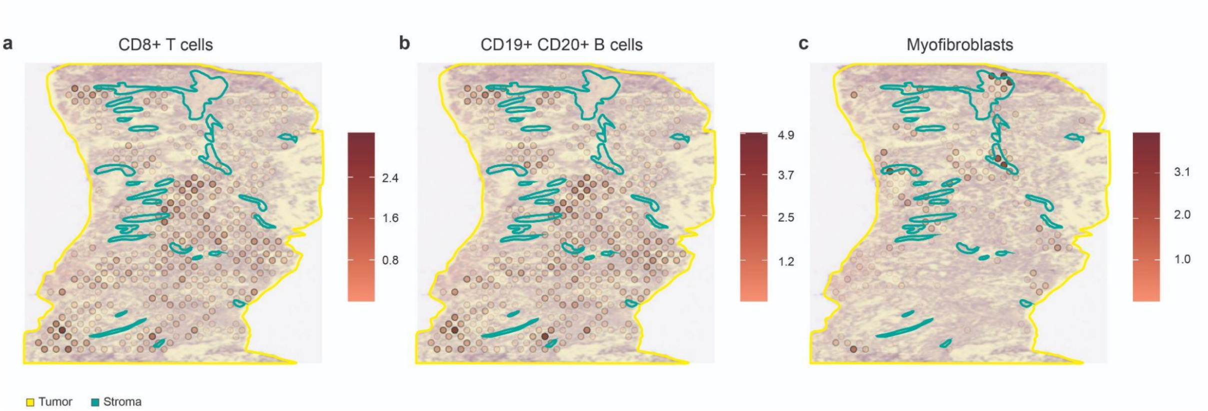


***Supplementary Figure S3:*** *Spatial mapping of the predicted number of a) CD8*$^{+}$ *T cells, b) CD19*$^{+}$ *CD20*$^{+}$ *B cells and c) myofibroblasts according to the deconvolution results overlaying with the pathologists's tissue annotations for the sample S1_Cec_Rep1.*

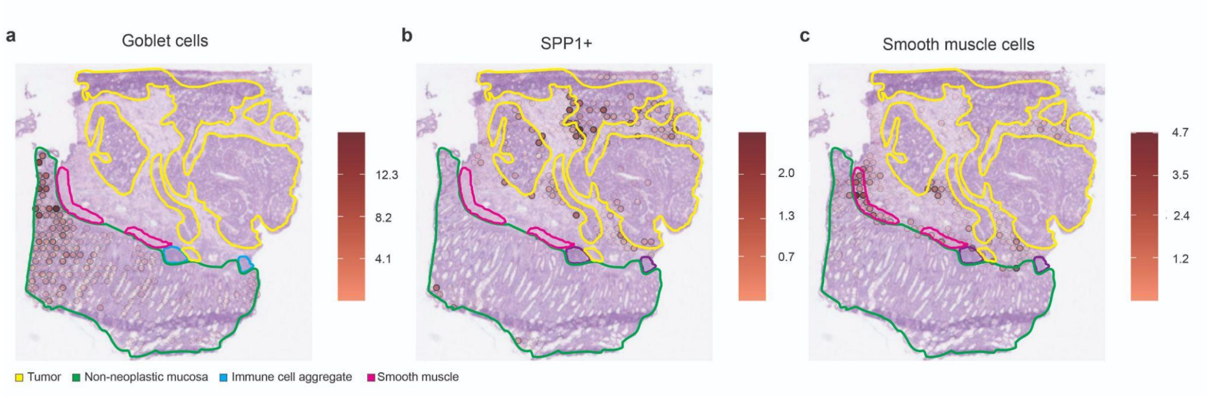


***Supplementary Figure S4:*** *Spatial mapping of the predicted number of a) goblet cells, b) SPP1*$^{+}$ *macrophages and c) smooth muscle cells according to the deconvolution results overlaying with the pathologists' tissue annotations for the sample S2_ Col_ R_ Rep1.*

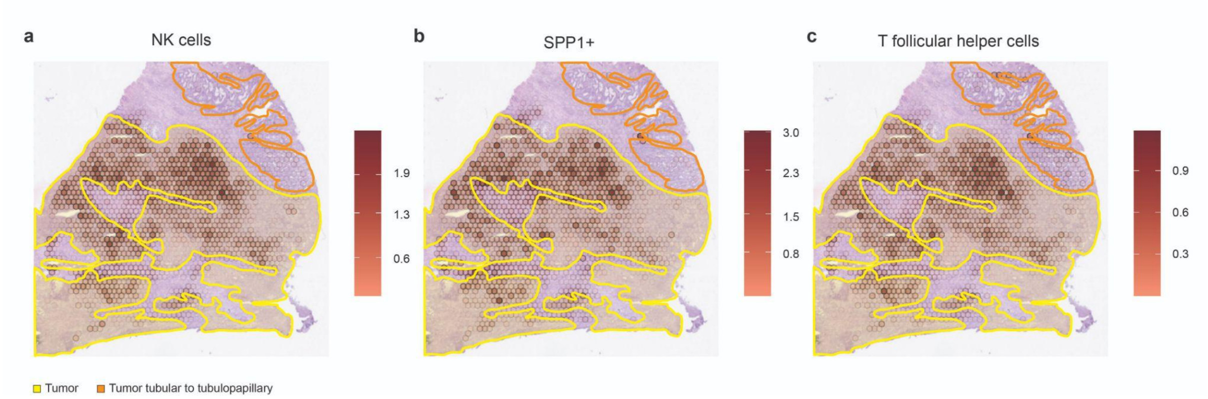


***Supplementary Figure S5:*** *Spatial mapping of the predicted number of a) NK cells, b) SPP1$^+$ macrophages and c) T follicular helper cells according to the deconvolution results overlaying with the pathologists' tissue annotations for the sample S3_ Col_ Rep1.*

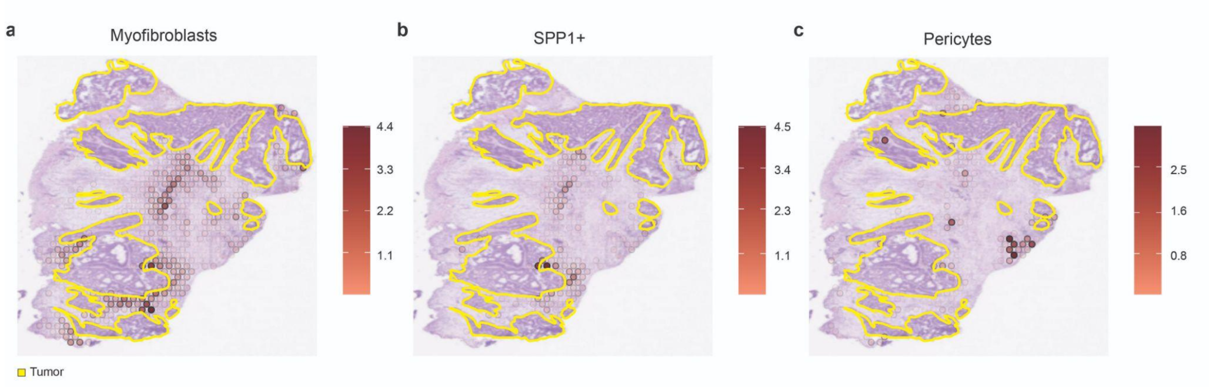


***Supplementary Figure S6:*** *Spatial mapping of the predicted number of a) myofibroblasts, b) SPP1$^{+}$ macrophages and c) pericytes according to the deconvolution results overlaying with the pathologists' tissue annotations for the sample S4_ Col_ Sig_ Rep2.*

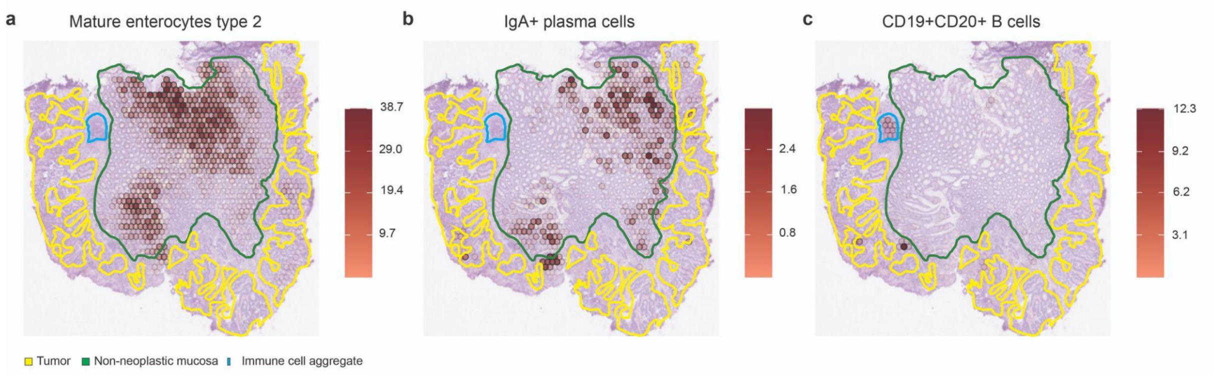


***Supplementary Figure S7:*** *Spatial mapping of the predicted number of a) mature enterocytes type 2, b) IgA$^{+}$ plasma cells and c) CD19$^{+}$ CD20$^{+}$ B cells according to the deconvolution results overlaying with the pathologists' tissue annotations for the sample S5_Rec_Rep1.*

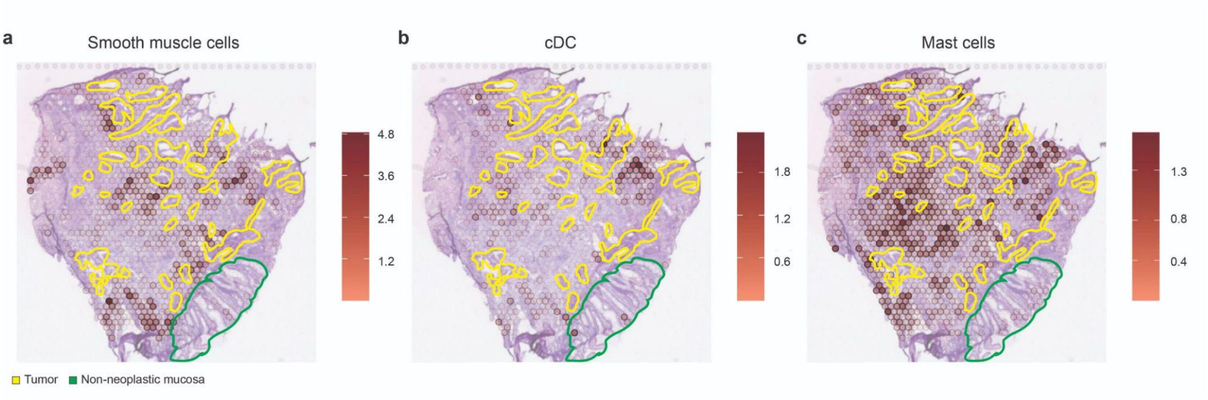


***Supplementary Figure S8:*** *Spatial mapping of the predicted number of a) smooth muscle cells, b) conventional dendritic cells (cDC) and c) mast cells according to the deconvolution results overlaying with the pathologists' tissue annotations for the sample S6_Rec_Rep2.*

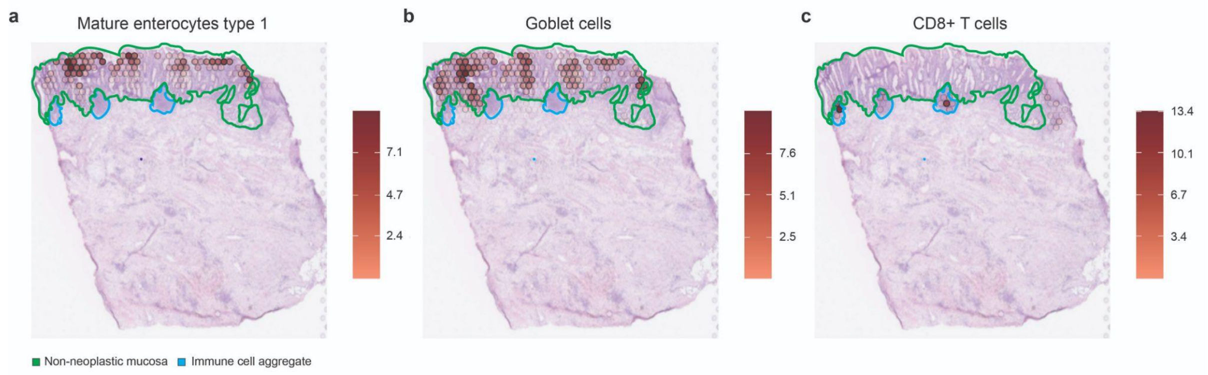


***Supplementary Figure S9:*** *Spatial mapping of the predicted number of a) mature enterocytes type 1, b) goblet cells and c) CD8+ T cells according to the deconvolution results overlaying with the pathologists' tissue annotations for the sample S7_Rec/Sig_Rep1.*

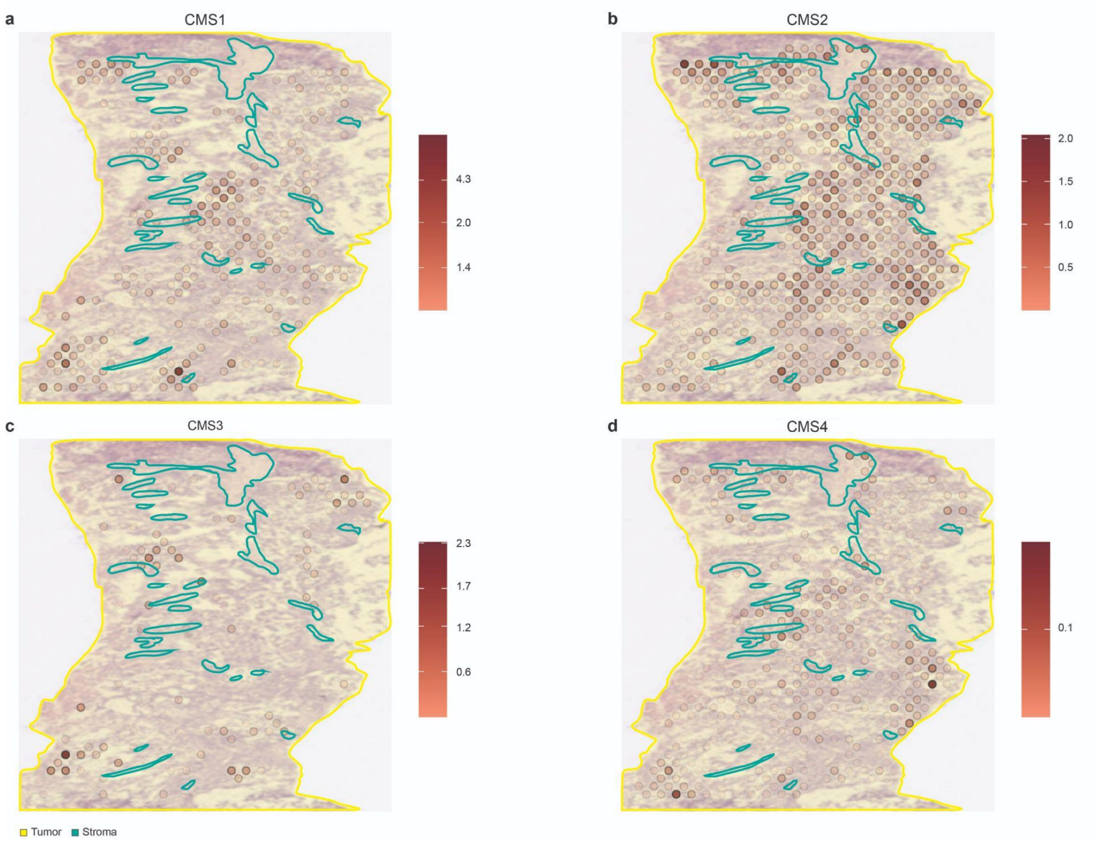


***Supplementary Figure S10:*** *Spatial mapping of the abundance of the different CMS signatures according to the deconvolution results overlaying with the pathologists' tissue annotations for the sample S1_ Cec_ Rep1.*

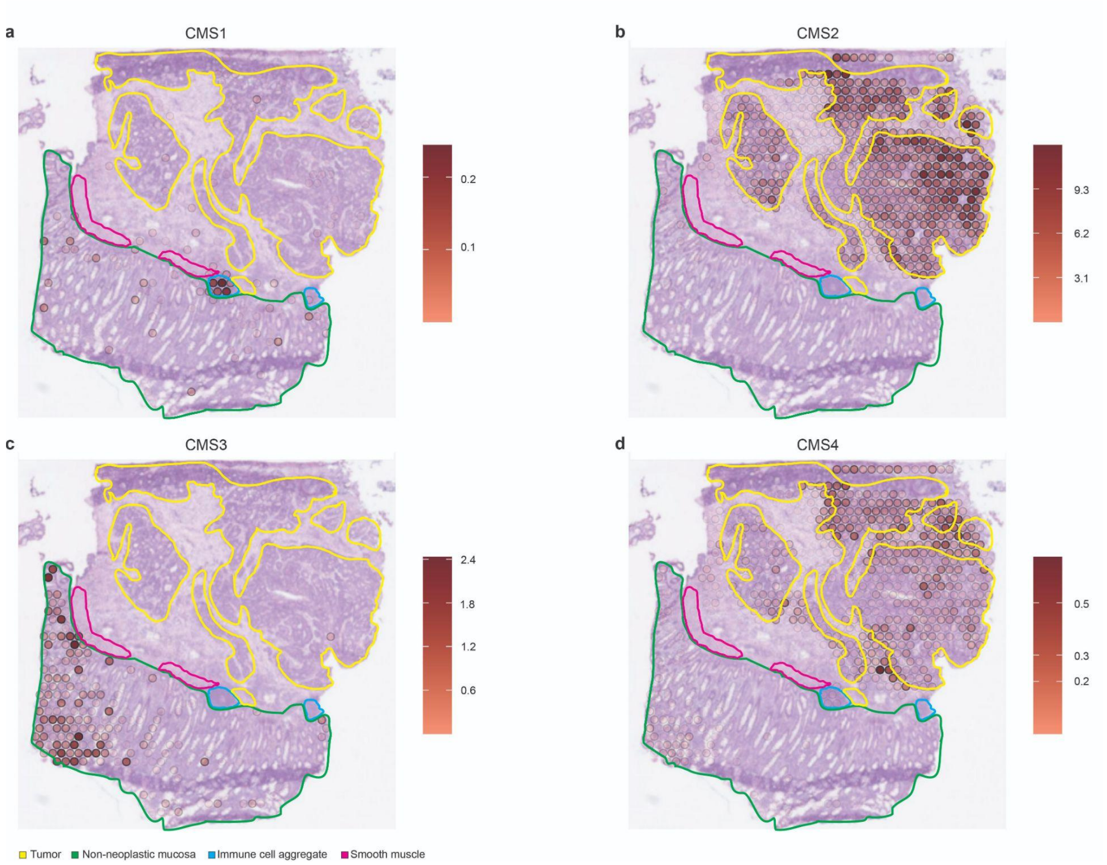


***Supplementary Figure S11:*** *Spatial mapping of the abundance of the different CMS signatures according to the deconvolution results overlaying with the pathologists' tissue annotations for the sample S2_Col_R_Rep1.*

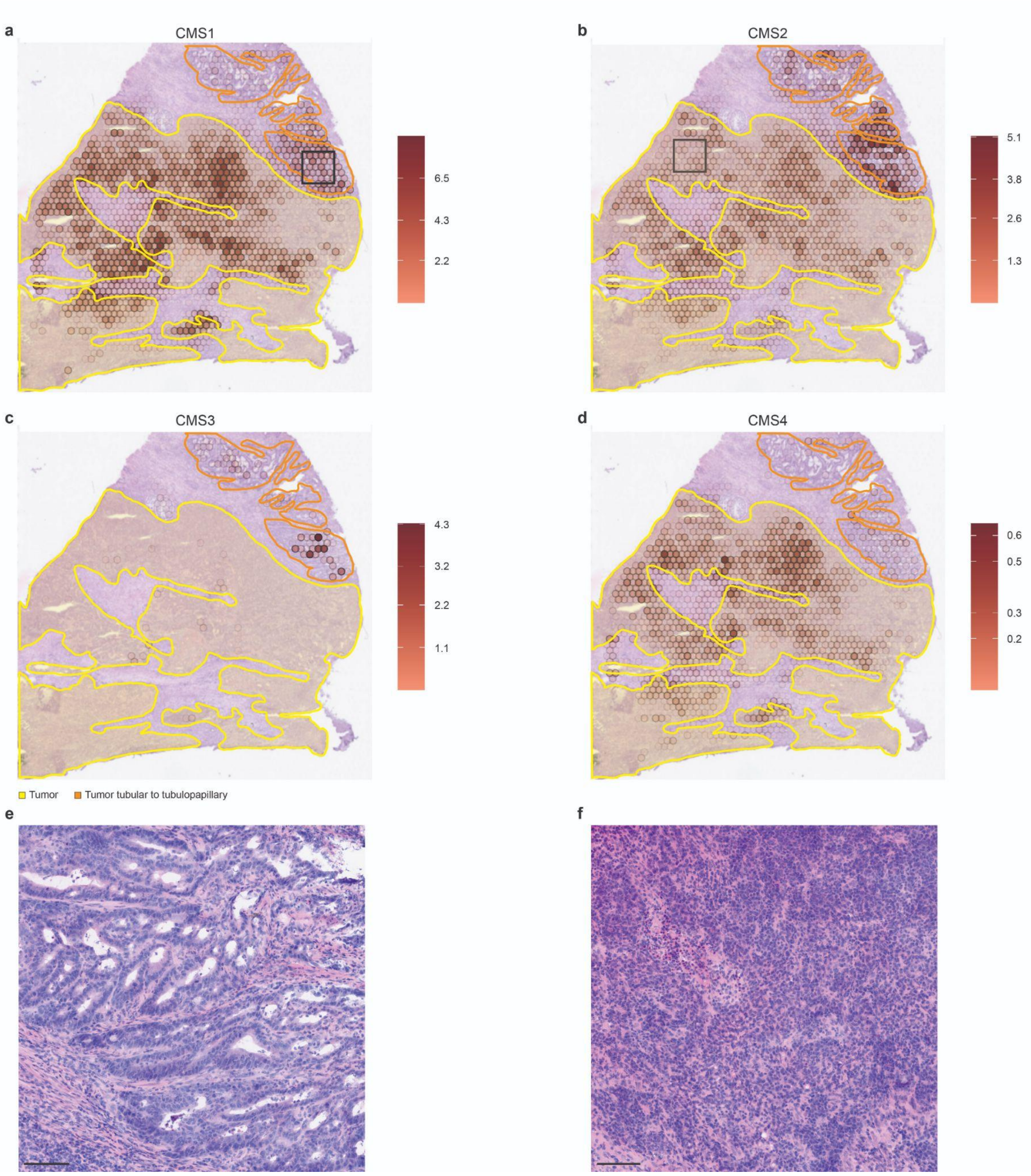


***Supplementary Figure S12:*** *Spatial mapping of the abundance of the different CMS signatures according to the deconvolution results overlaying with the pathologists' tissue annotations for the sample S3_ Col_ Rep1. Deconvolution results in the sample S3_ ColR_ S2 showing a predominant CMS2 signature in anatomical regions with a tubular to tubulopapillary growth pattern at the morphological level (e). The former are delineated from regions with a predominant CMS1 signature which are linked to morphological features displaying a predominant diffuse growth pattern (f). (e, f) Hematoxylin and eosin staining. Scale bar 100 µm.*

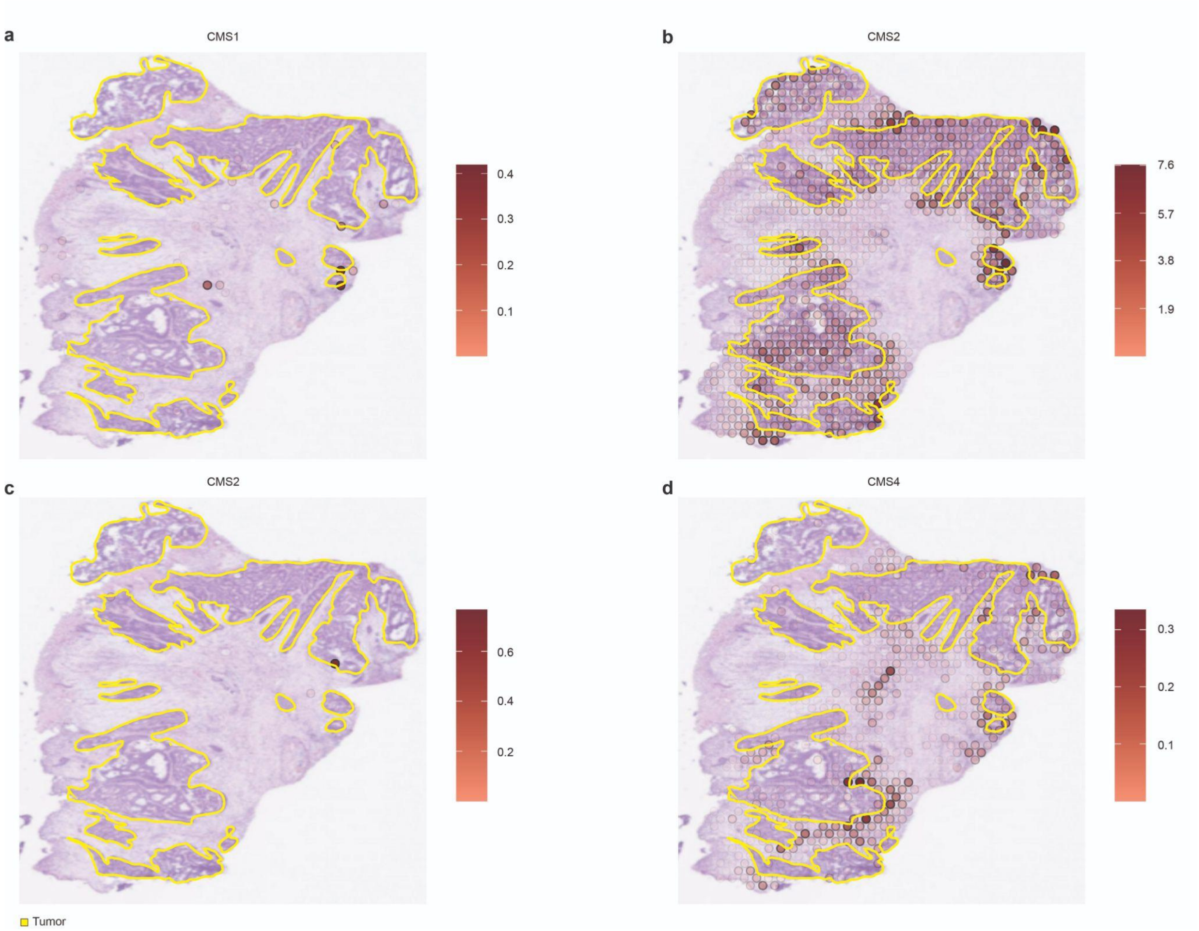


***Supplementary Figure S13:*** *Spatial mapping of the abundance of the different CMS signatures according to the deconvolution results overlaying with the pathologists' tissue annotations for the sample S4_Col_Sig_Rep2.*

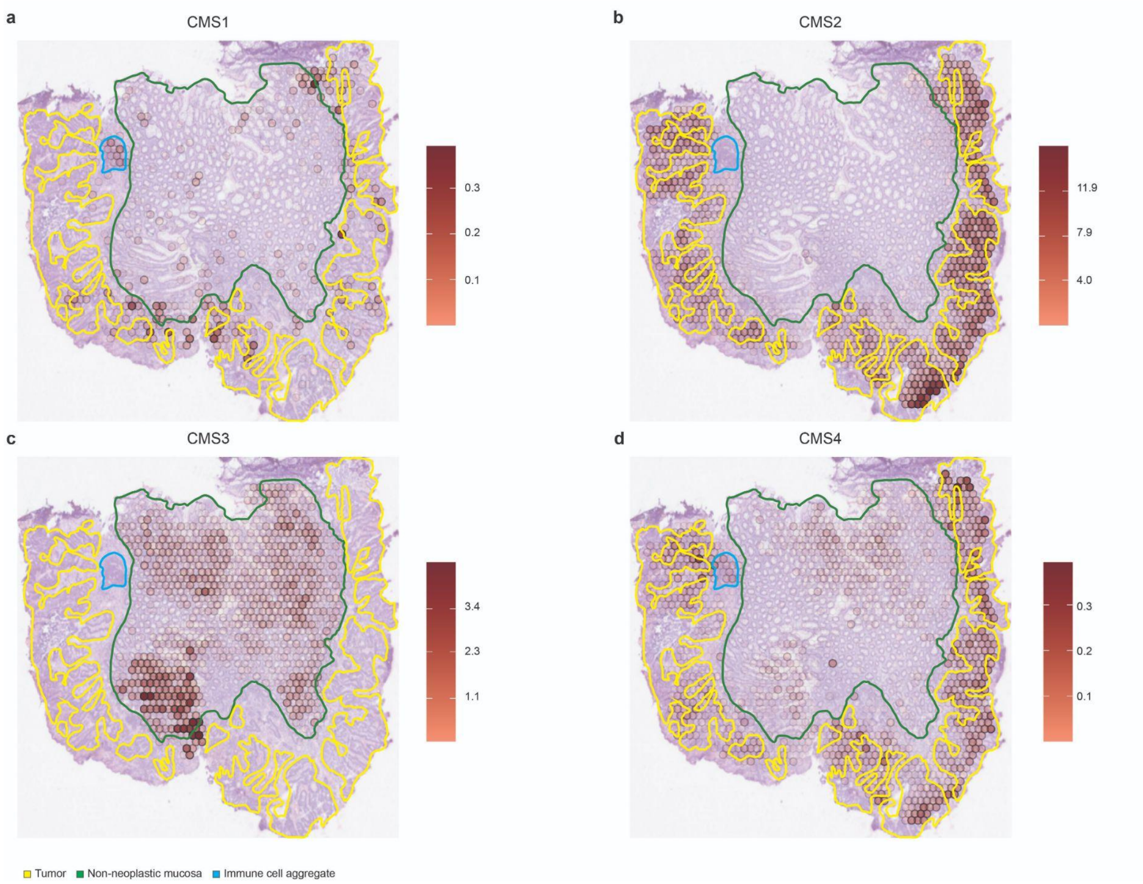


***Supplementary Figure S14:*** *Spatial mapping of the abundance of the different CMS signatures according to the deconvolution results overlaying with the pathologists' tissue annotations for the sample S5_Rec_Rep1.*

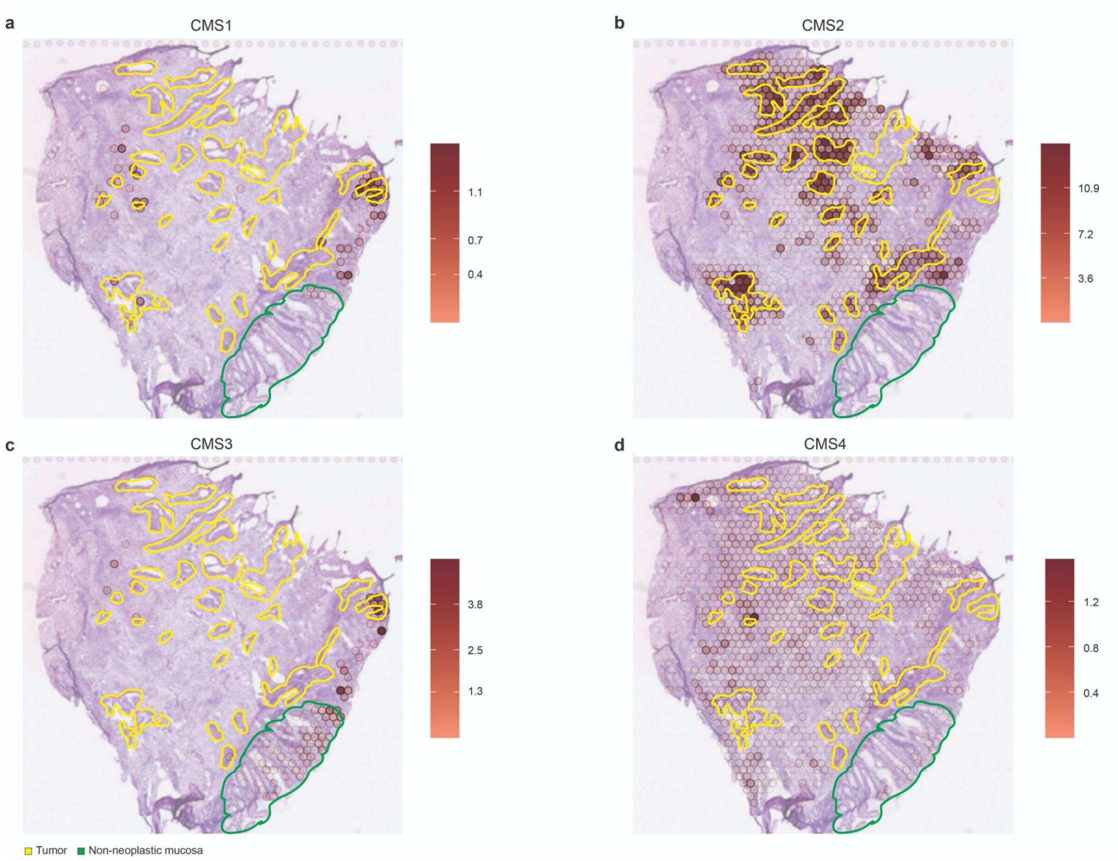


***Supplementary Figure S15:*** *Spatial mapping of the abundance of the different CMS signatures according to the deconvolution results overlaying with the pathologists' tissue annotations for the sample S6_Rec_Rep2.*

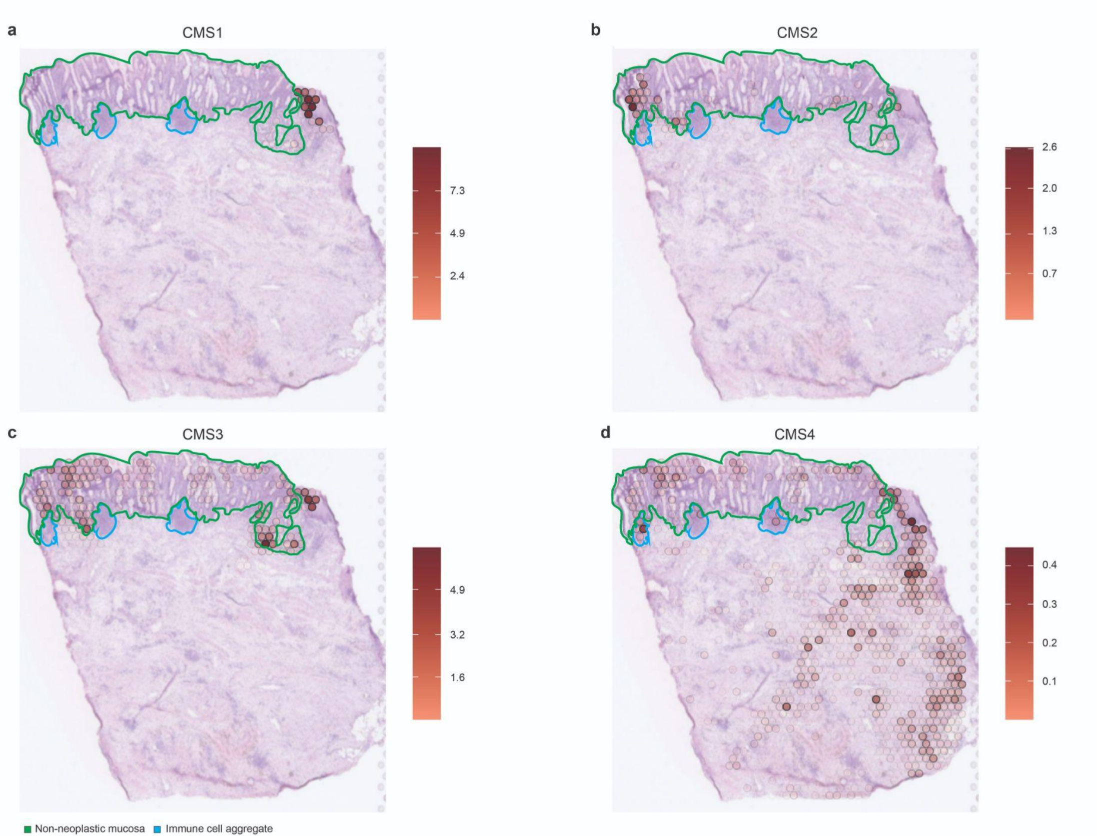


***Supplementary Figure S16:*** *Spatial mapping of the abundance of the different CMS signatures according to the deconvolution results overlaying with the pathologists' tissue annotations for the sample S7_Rec/Sig_Rep1.*

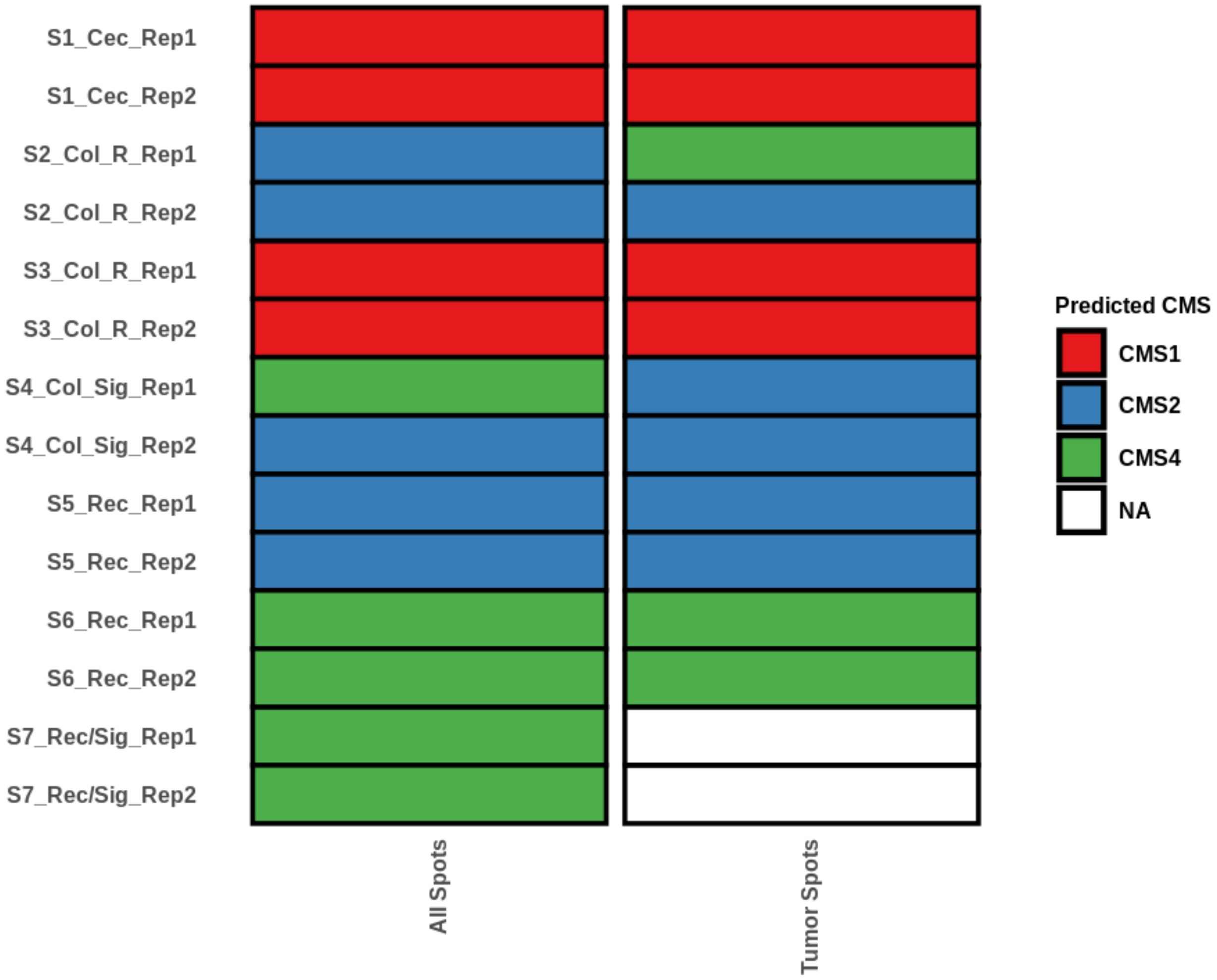


***Supplementary Figure S17:*** *Predicted CMS classification resulting from applying CMScaller on pseudo-bulk RNA-seq generated by either pooling together all the spots or only the tumor annotated spots for each of our CRC samples.*

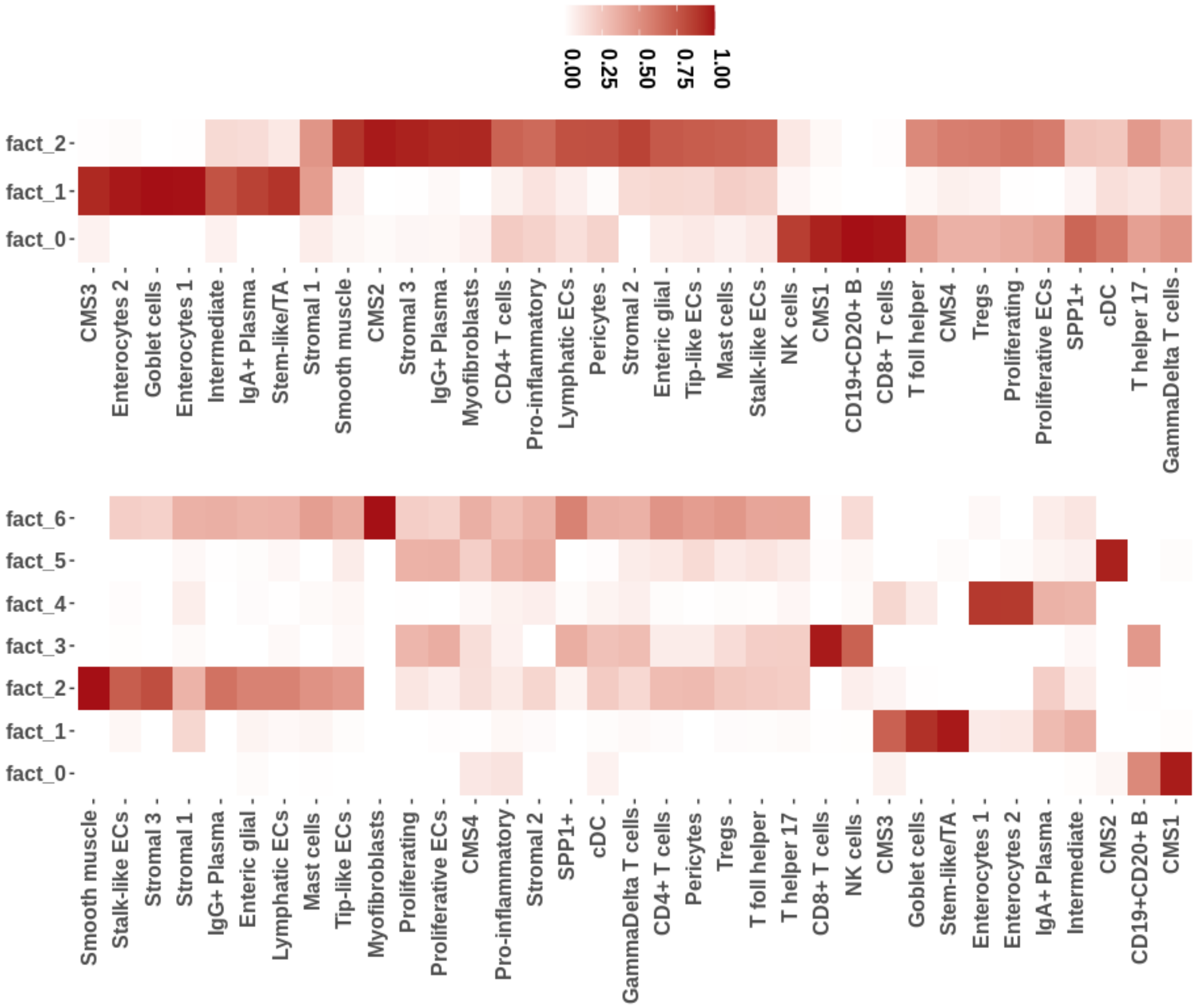


***Supplementary Figure S18:*** *Cell type co-localization results when considering 3 factors (above) or 7 factors (below) across all our samples and replicates. A number larger of factors indicates stronger co-localization signals along more confined anatomical regions.*

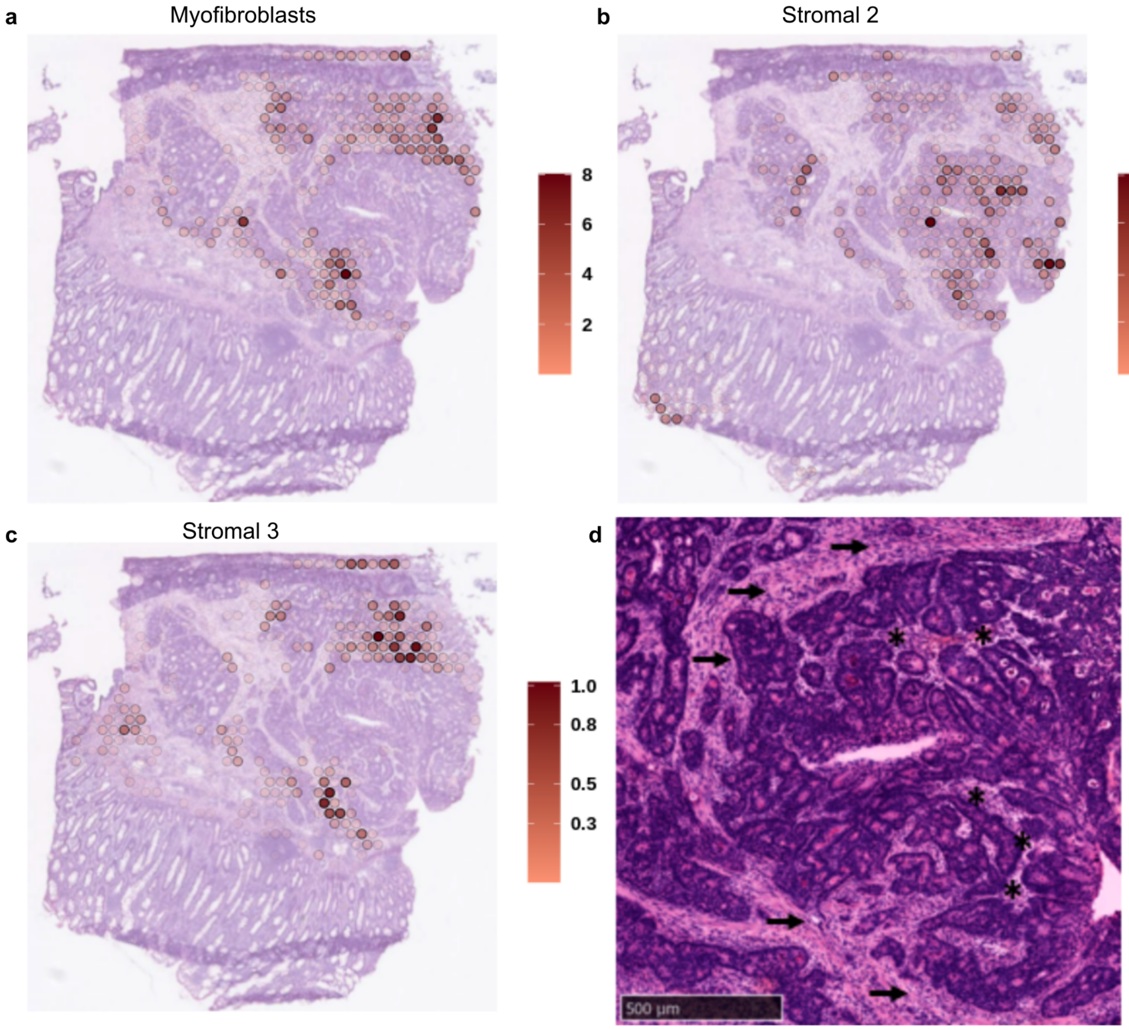


***Supplementary Figure S19: Deconvolution results for stromal cells and its assessment by the pathologists in the S2_Col_R_Rep1 sample.*** *Larger stroma bundles separating tumor lobes displaying a) a predominant myofibroblast and c) a minor stroma 3 (c) signature. b) Stroma 2 was predominantly identified in close association to tumor lobules. (d) The higher magnification shows the spatially distinct localization of the intratumoral stroma (asterisk) and the larger interlobular stroma bundles (arrows). Scale bar 500 µm.*

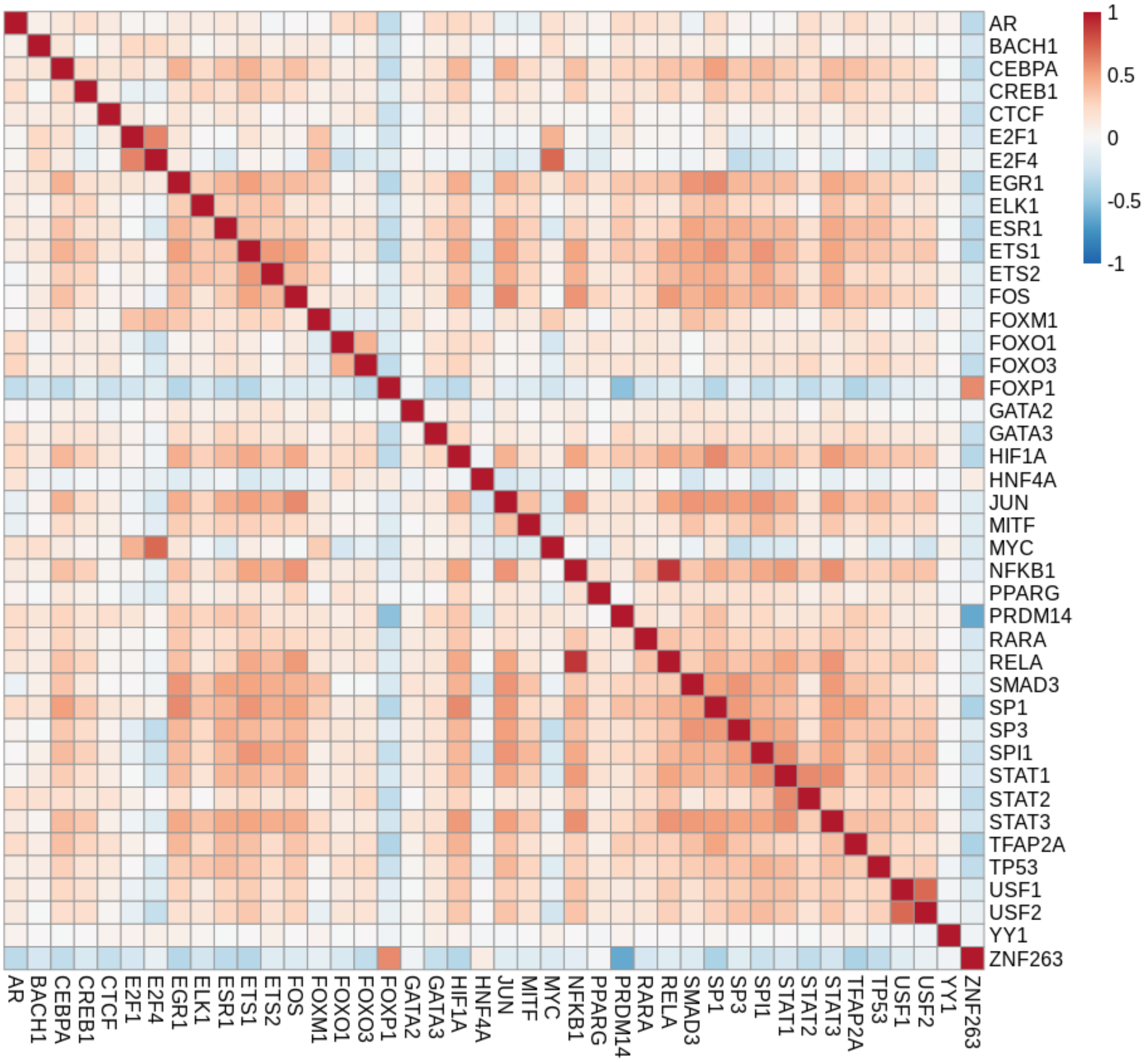


***Supplementary Figure S20:*** *Per spot Pearson's correlation of the TF activity across all the samples under study.*

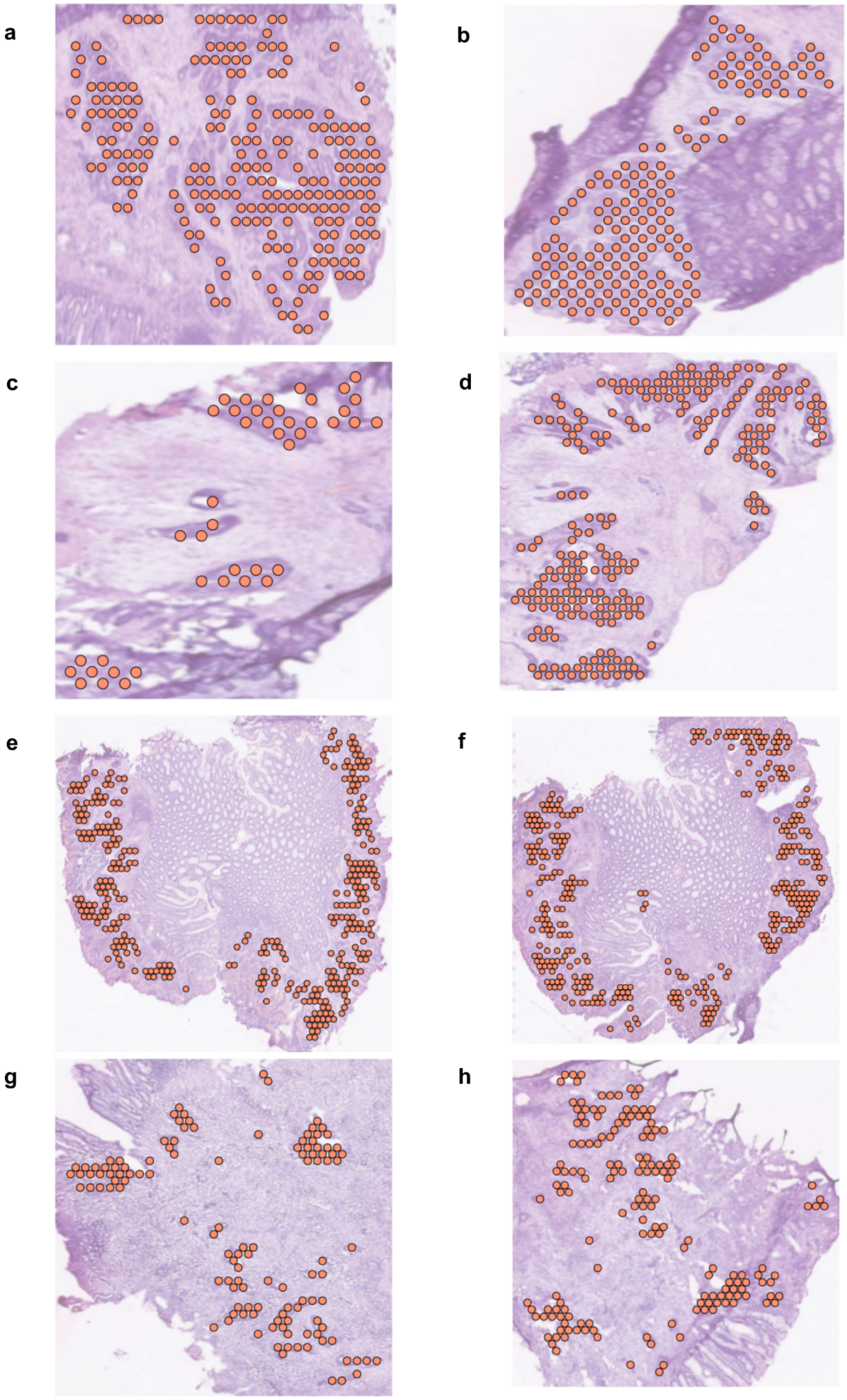

***Supplementary Figure S21:*** *Spots annotated as tumor by the pathologists for the CMS2 tumor samples, a) S2_Col_R_Rep1, b) S2_Col_R_Rep2, c) S4_Col_Sig_Rep1, d) S4_Col_Sig_Rep2, e) S5_Rec_Rep1, f) S5_Rec_Rep2, g) S6_Rec_Rep1 and h) S6_Rec_Rep2.*

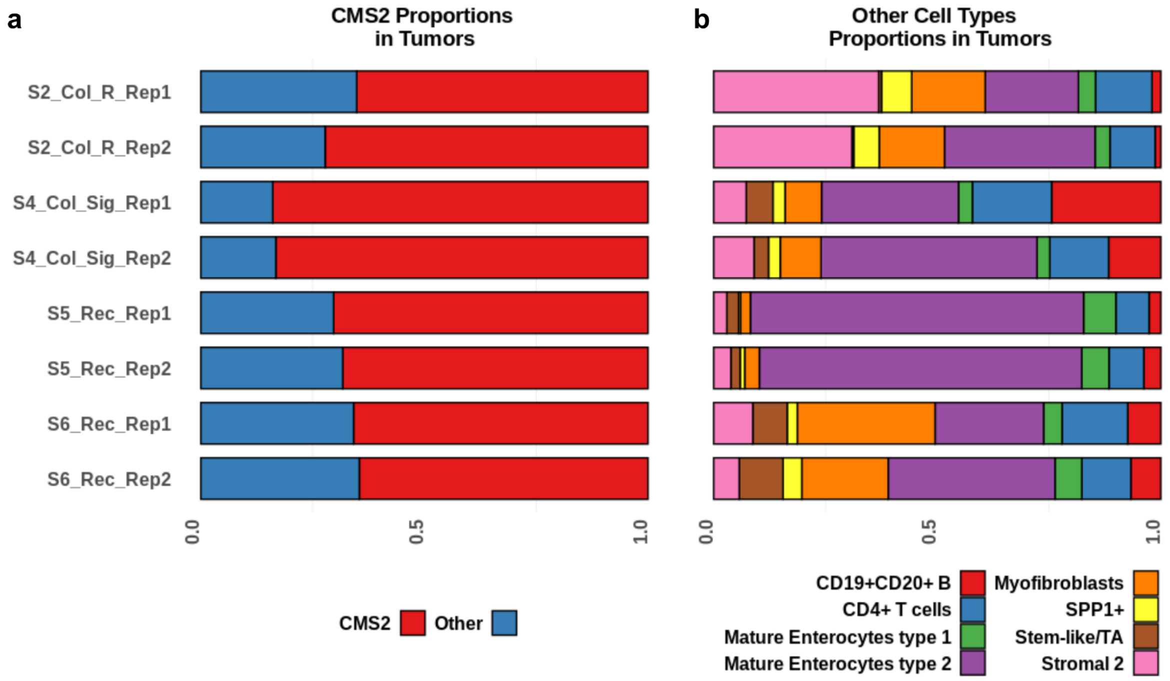


***Supplementary Figure S22:*** *Cell type proportions in the tumor-annotated spots per sample as estimated by the results of the deconvolution: a) CMS2 proportions as compared to all the remaining cell types considered in the study, and b) proportions of the most relevant cell types within CMS2 tumors excluding CMS2 cells.*

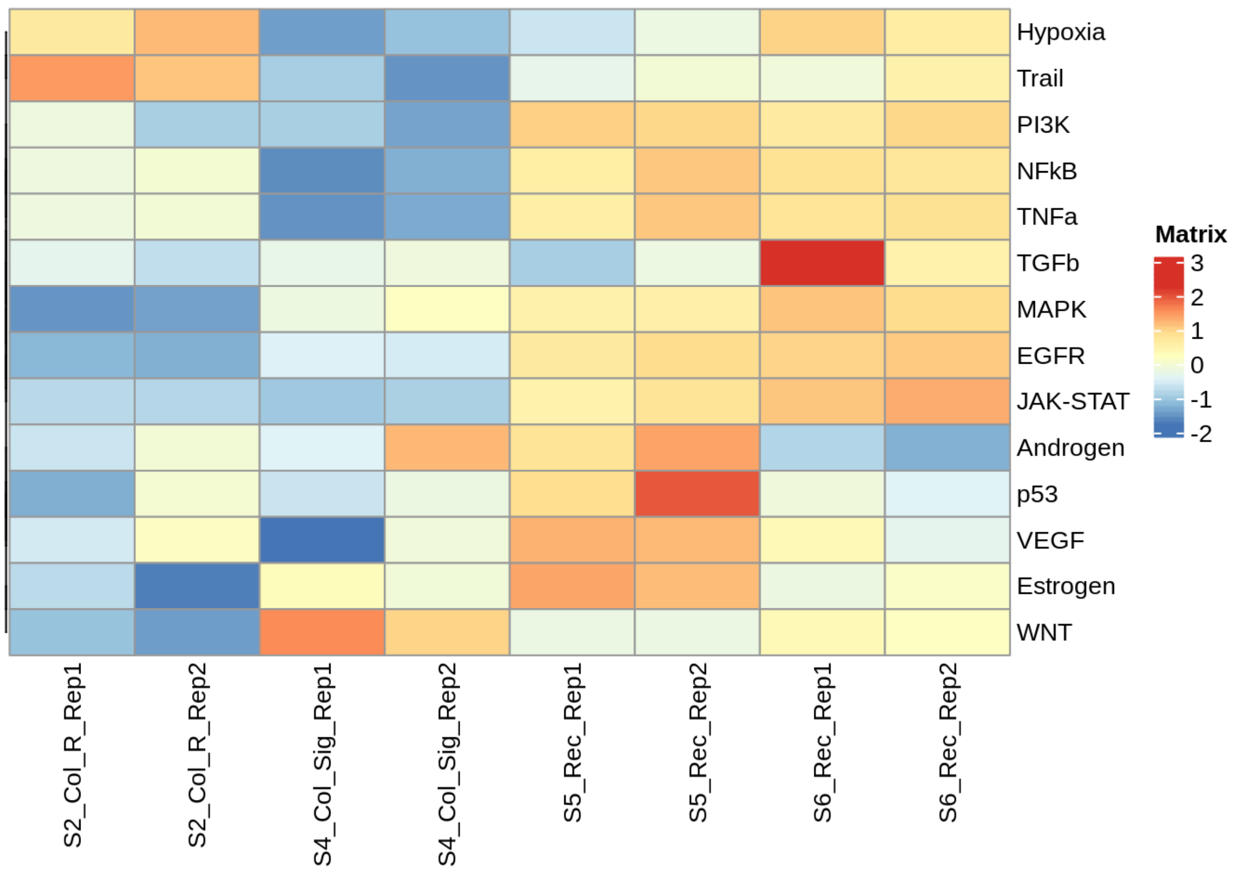


***Supplementary Figure S23:*** *Differential pathway activity computed on pseudo-bulk RNA-seq generated from the tumor-annotated spots for the different CMS2 samples.*

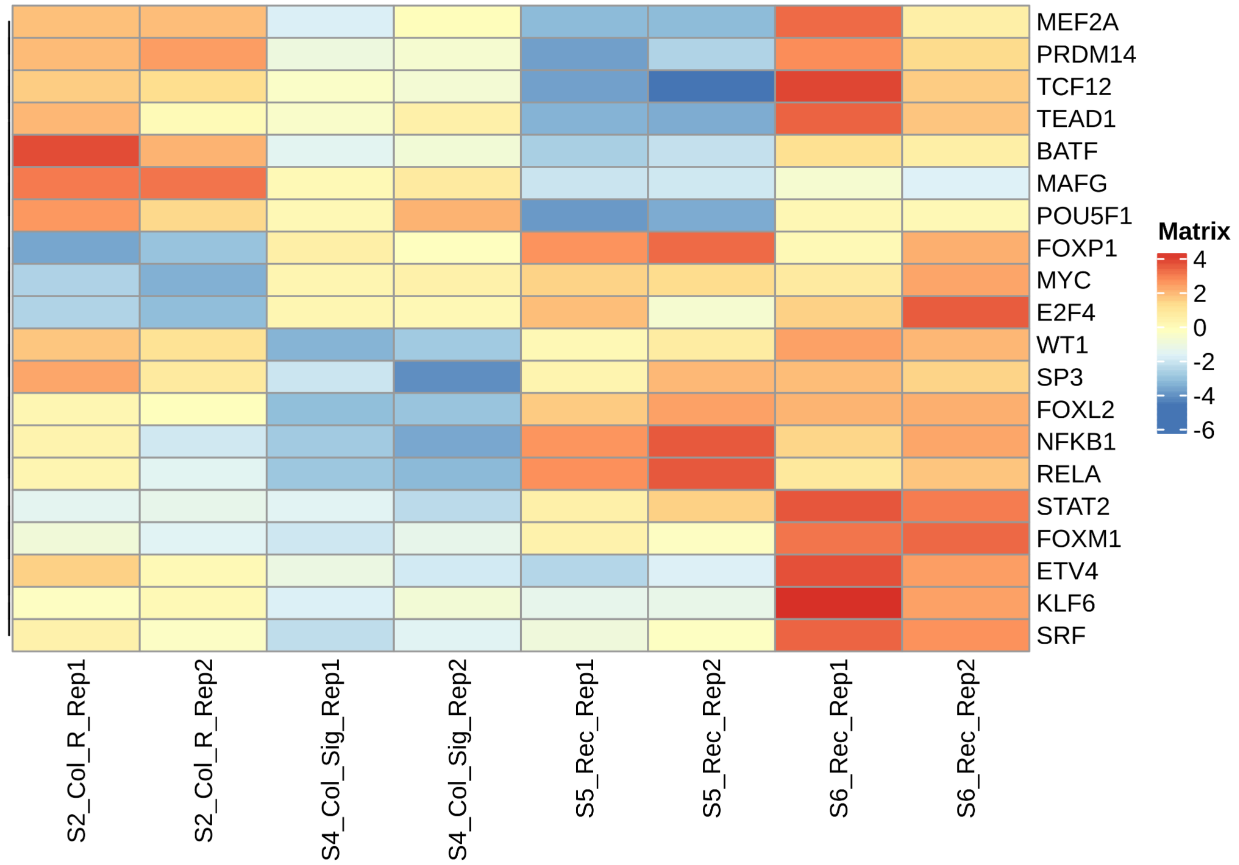


***Supplementary Figure S24:*** *Differential transcriptional activity of selected TFs computed on pseudo-bulk RNA-seq generated from the tumor-annotated spots for the different CMS2 samples.*

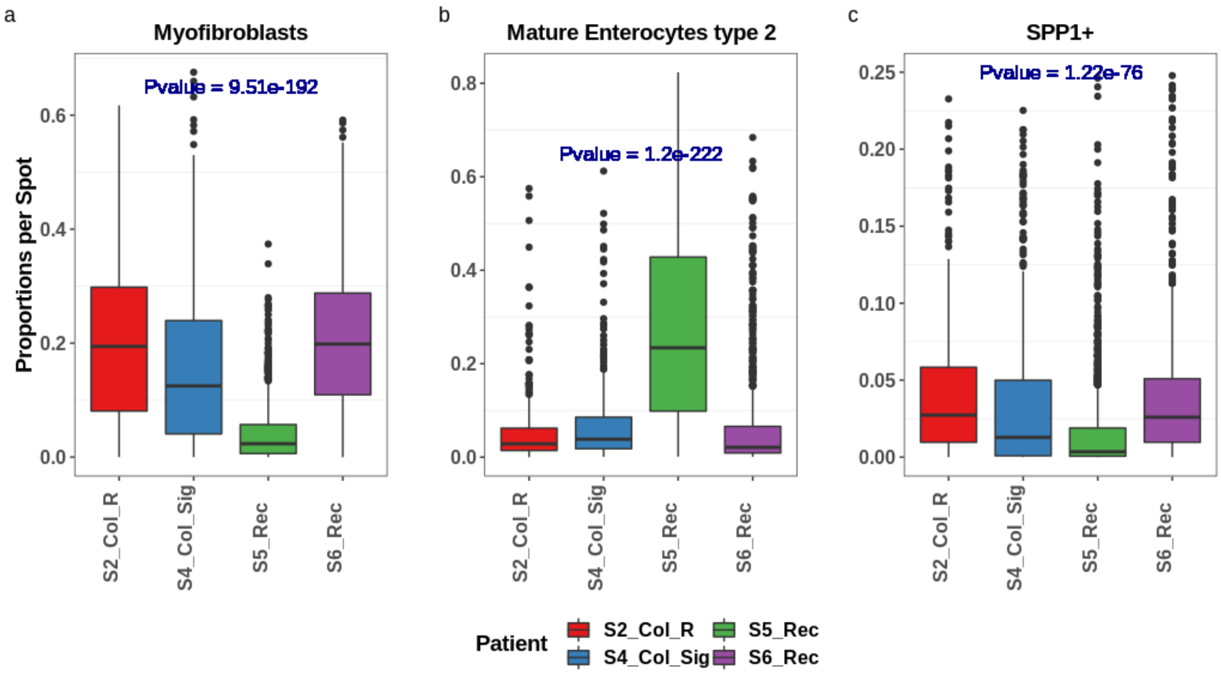


***Supplementary Figure S25:*** *Myofibroblast, Mature enterocytes type 2 and SPP1+ proportions per spot in the tumor neighborhood of the different patients (both replicates considered) as estimated by the results of the deconvolution approach. A Kruskal-Wallis statistical test was performed to assess whether the cell type proportions for the different samples originated from the same distribution (p-value).*

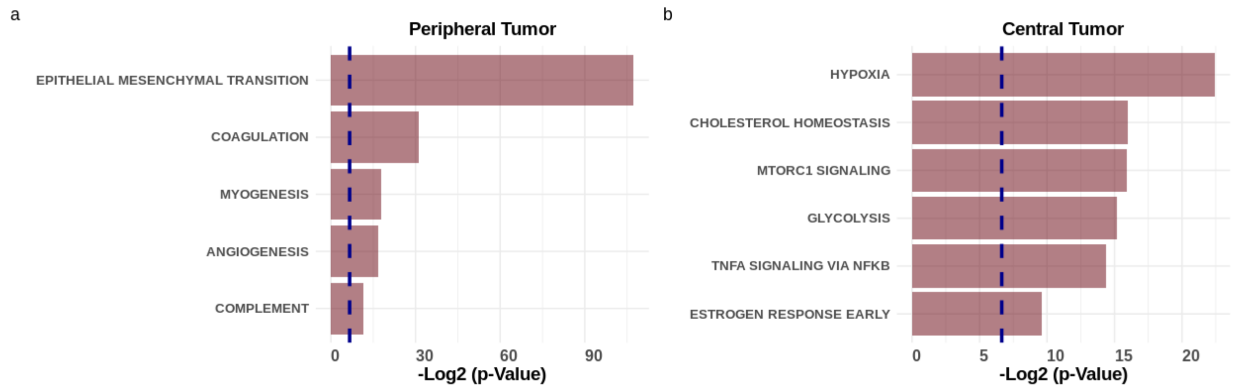


***Supplementary Figure S26:*** *Significant enrichment results (p-value<0.01) on the differentially expressed genes between the different anatomical regions of the CMS2 tumor in sample S2_Col_R_Rep1.*

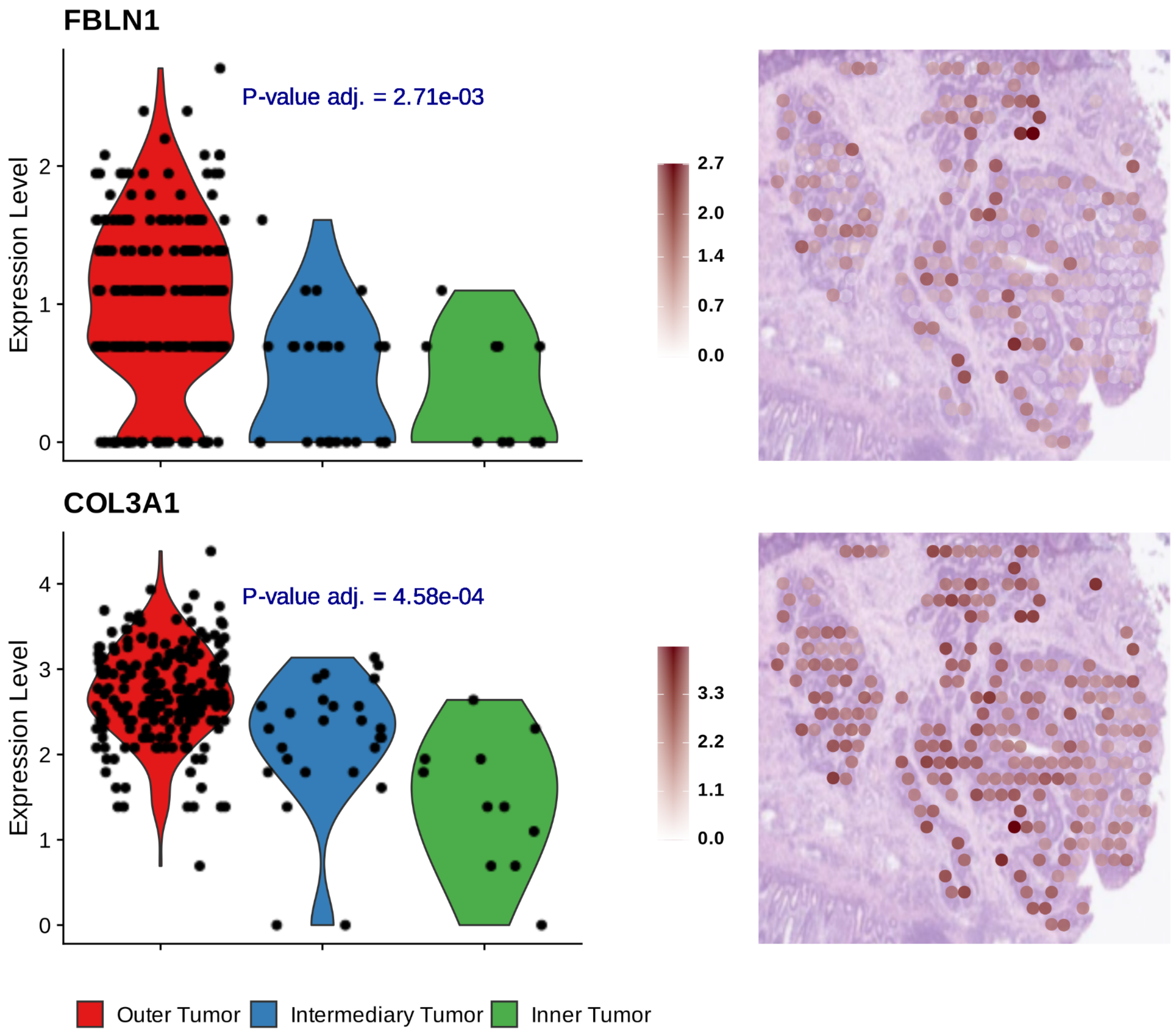


***Supplementary Figure S27:*** *Expression gradients of FBLN1 and COL3A1 in the different anatomical regions of tumor-annotated spots in the S2_Col_R_Rep1 sample. A Wilcoxon rank sum test was conducted to assess the significance of the gene expression variation (p-value adjusted).*

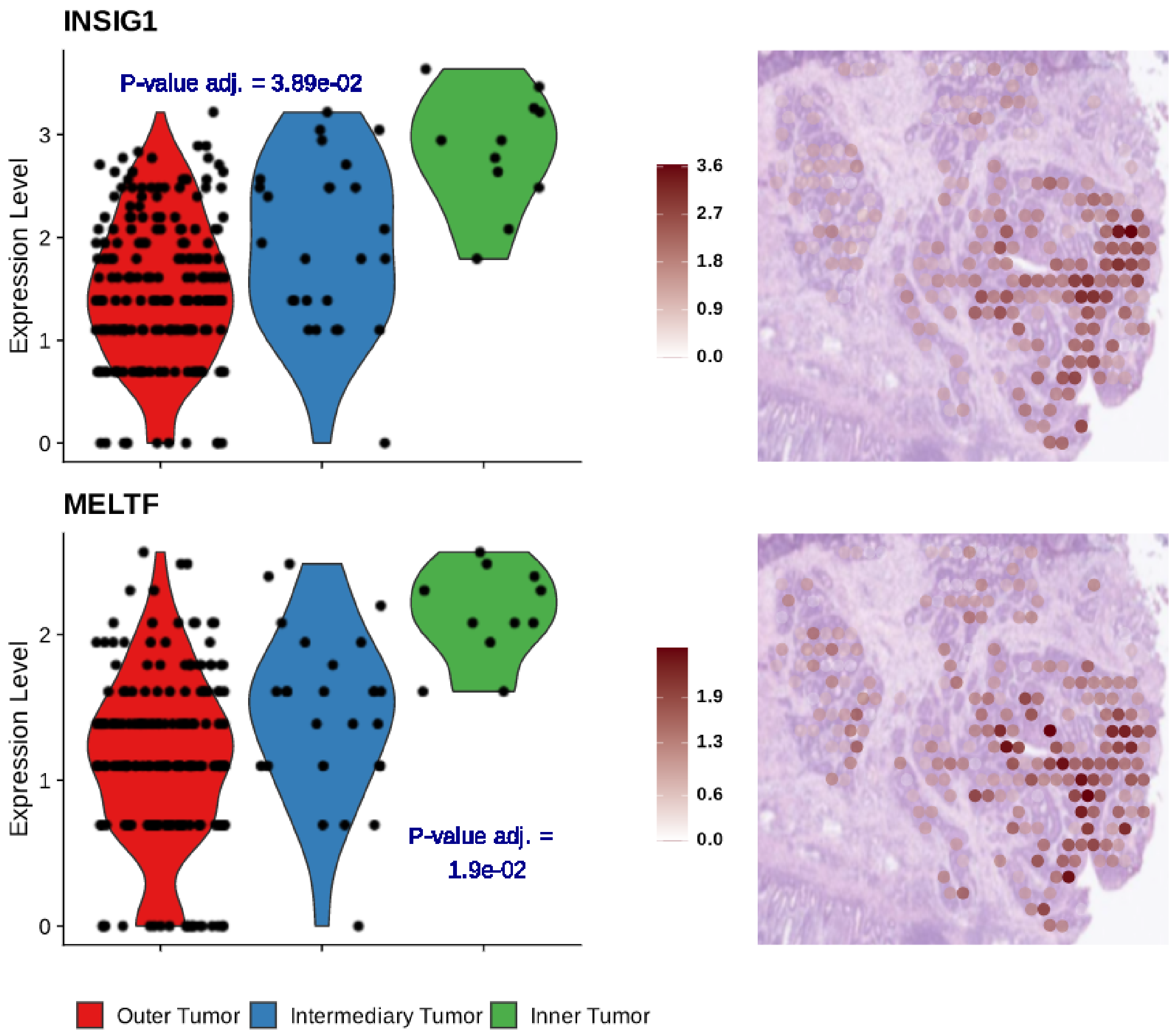


***Supplementary Figure S28:*** *Expression gradients of INSIG1 and MELTF in the different anatomical regions of tumor-annotated spots in the S2_Col_R_Rep1 sample. A Wilcoxon rank sum test was conducted to assess the significance of the gene expression variation (p-value adjusted).*

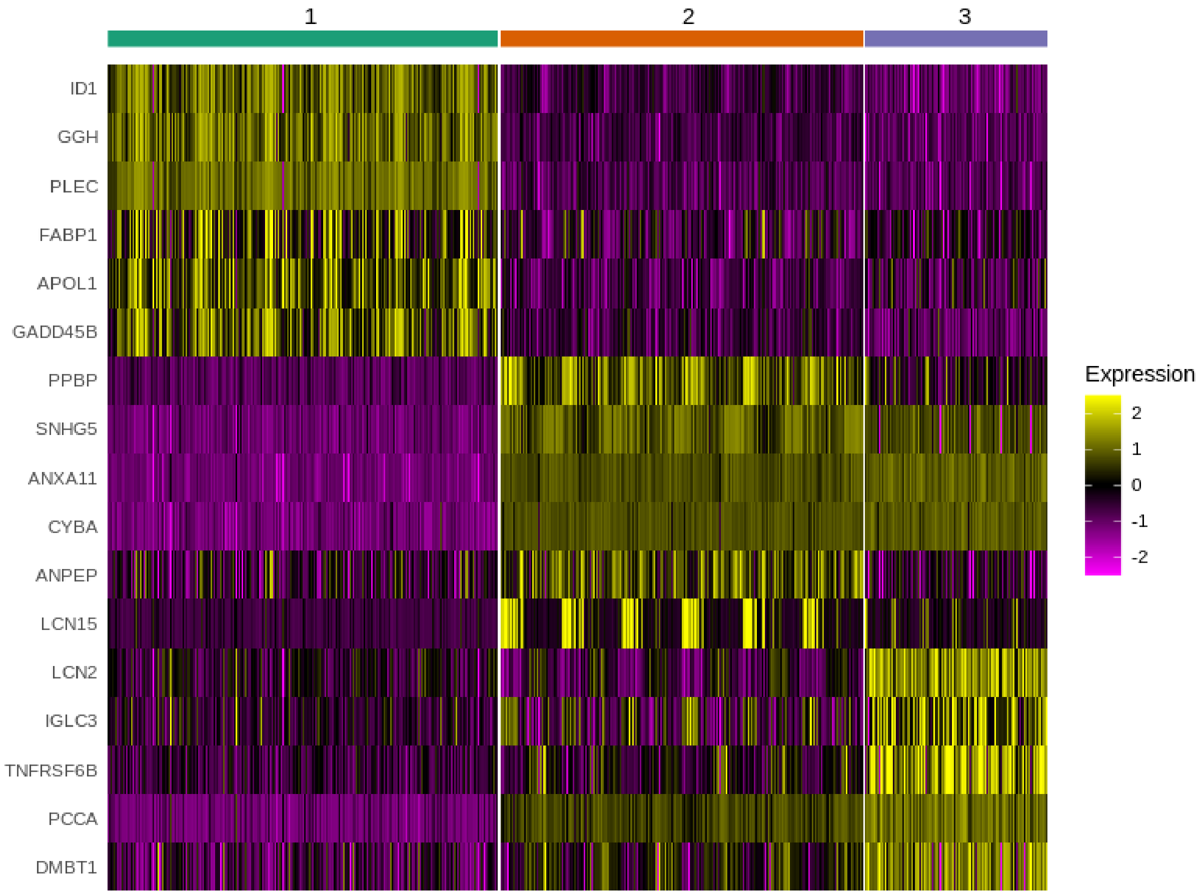


***Supplementary Figure S29:*** *Heatmap displaying the differentially expressed genes with a larger fold-change per group between the different subclusters within the CMS2 tumor-annotated spots in the S5_Rec_Rep1 sample. For visualization purposes, a maximum of 6 markers are displayed per group.*

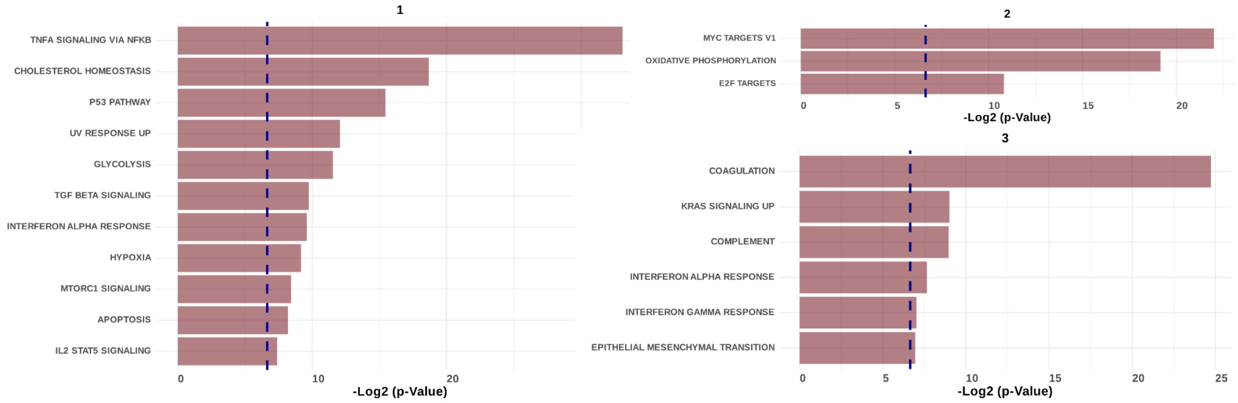


***Supplementary Figure S30:*** *Significant enrichment results (p-value<0.01) on the differentially expressed genes between the different sub-clustered regions of the CMS2 tumor in the S5_Rec_Rep1 sample.*

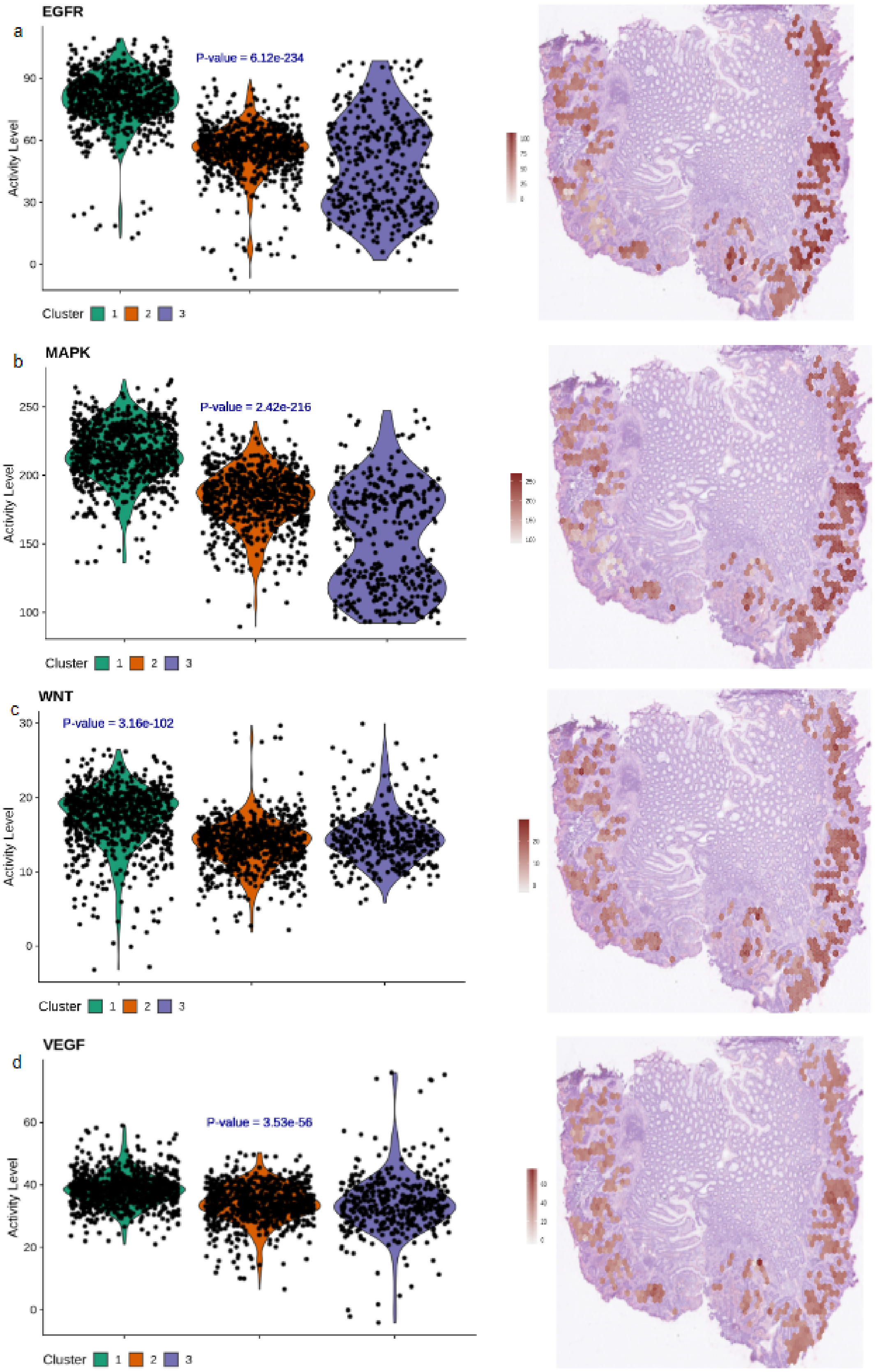
EGFR
a
P-value = 6.12e-234
Activity Level
90
60
30
0
Cluster 1 2 3
MAPK
b
P-value = 2.42e-216
Activity Level
250
200
150
100
Cluster 1 2 3
WNT
c
P-value = 3.16e-102
Activity Level
30
20
10
0
Cluster 1 2 3
VEGF
d
P-value = 3.53e-56
Activity Level
60
40
20
0
Cluster 1 2 3

***Supplementary Figure S31: Pathway activities at the enhanced subspot resolution****. Per cluster violin plots and spatial distribution for the following pathways: a) EGFR, b) MAPK, c) WNT and d) VEGF in the S5_Rec_Rep1 sample. A Kruskal-Wallis statistical test was performed in each case to assess whether the pathway activities in the different subclusters originated from the same distribution (p-value).*

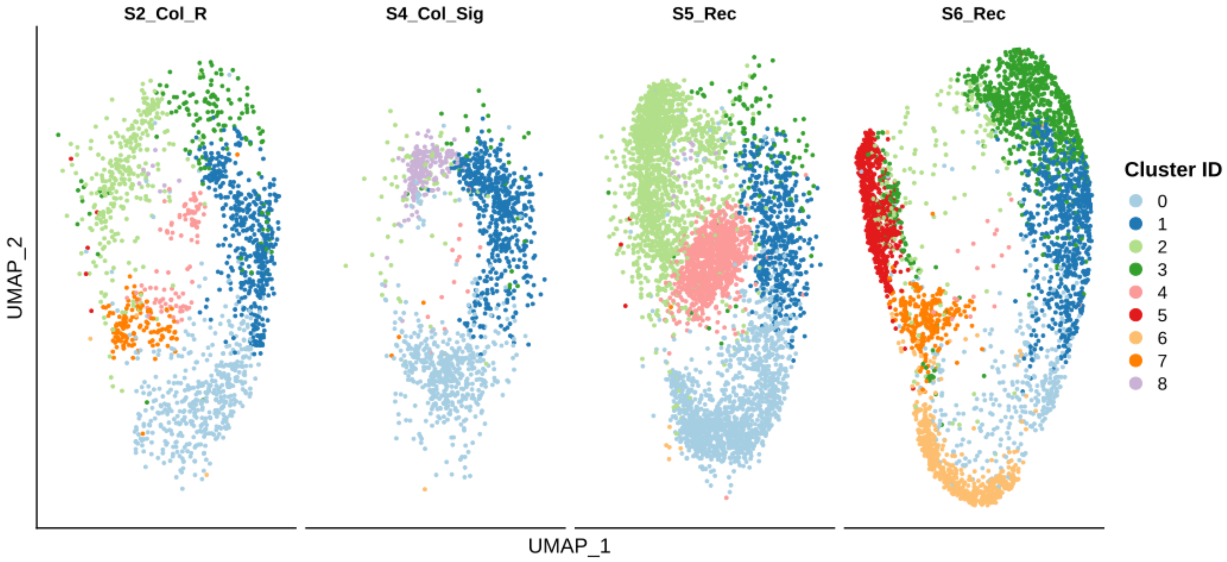


***Supplementary Figure S32:*** *UMAP embedding of the TF activity profiles for our set of CMS2 samples split by patient. The spots were colored by cluster identifiers.*

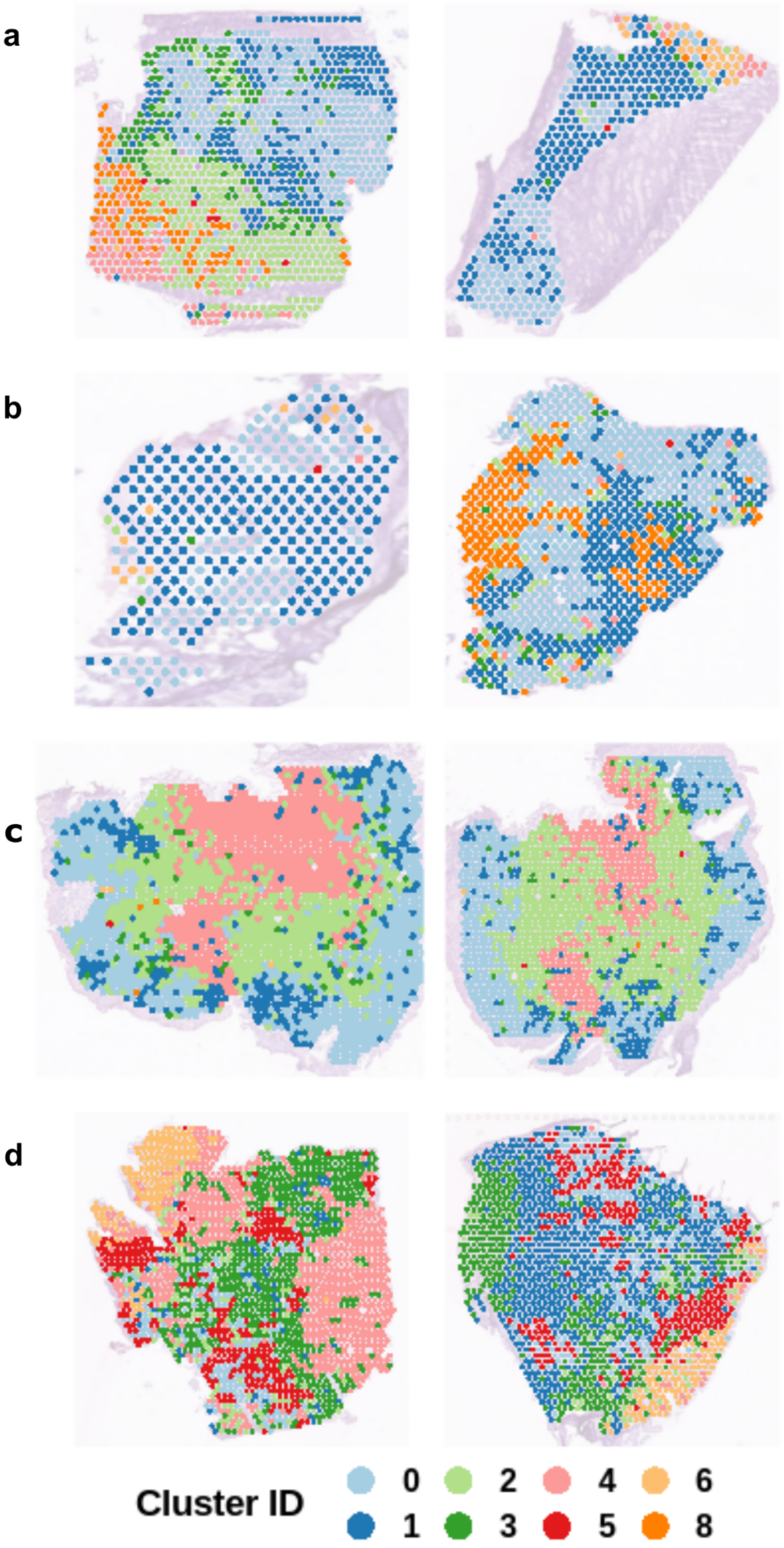

a
b
c
d
Cluster ID
0
1
2
3
4
5
6
8

***Supplementary Figure S33:*** *Spatial arrangement of the clusters based on the TF activity profiles per spot in our four CMS2 tumor patients: a)S2_Col_R, b) S4_Col_Sig, c)S5_Rec and d) S6_Rec. Both replicates are displayed per patient. The spots were colored by cluster identifiers.*

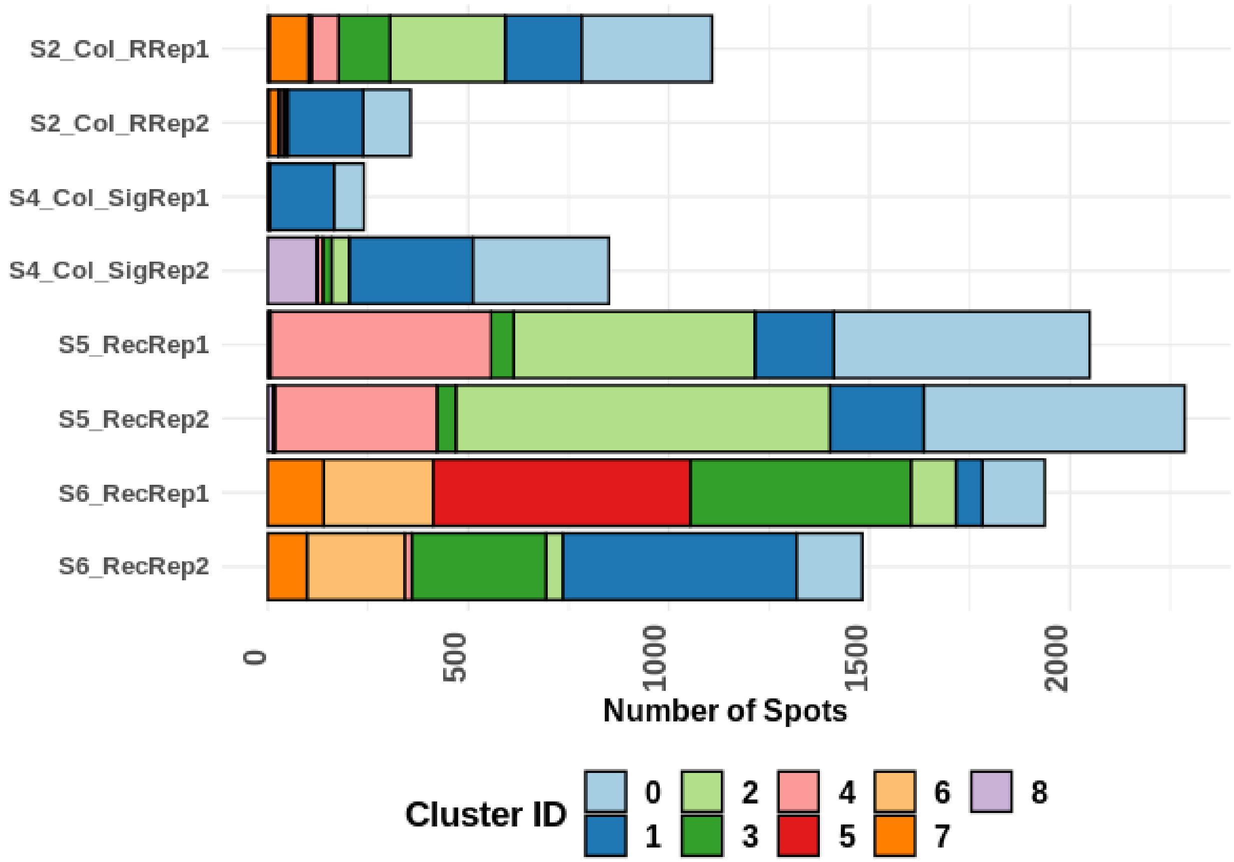


***Supplementary Figure S34:*** *Number of spots per sample and per cluster identifier according to the results of the clustering based on TF activity profiles.*

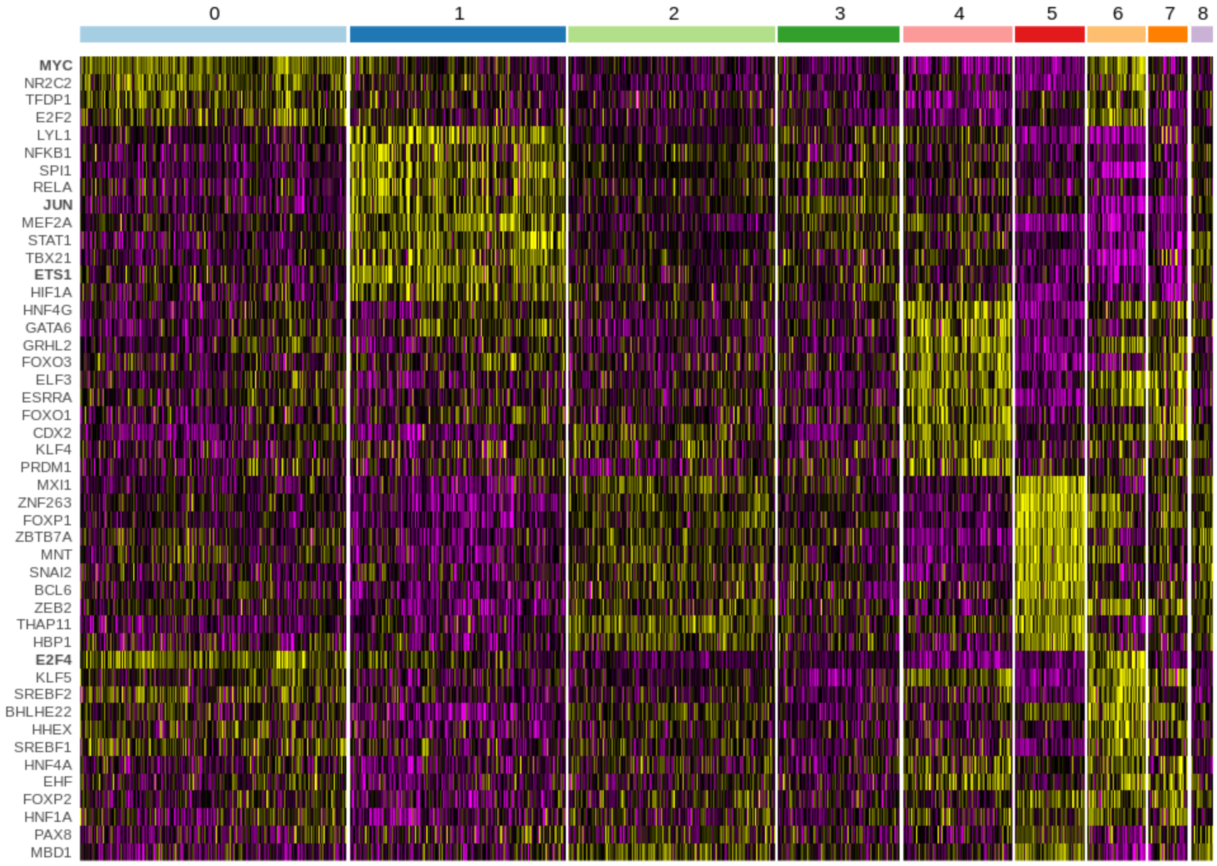


***Supplementary Figure S35:*** *Most differentially active TF between the different clusters based on the TF activity profiles per spot in our four CMS2 tumor patients (S2_Col_R, S4_Col_Sig, S5_Rec and S6_Rec). We display TF with an average* $Log_2$ *fold-change > 1.25 and a maximum of 10 TF per cluster. Bold characters were used to highlight the name of TFs referred to in the main text.*

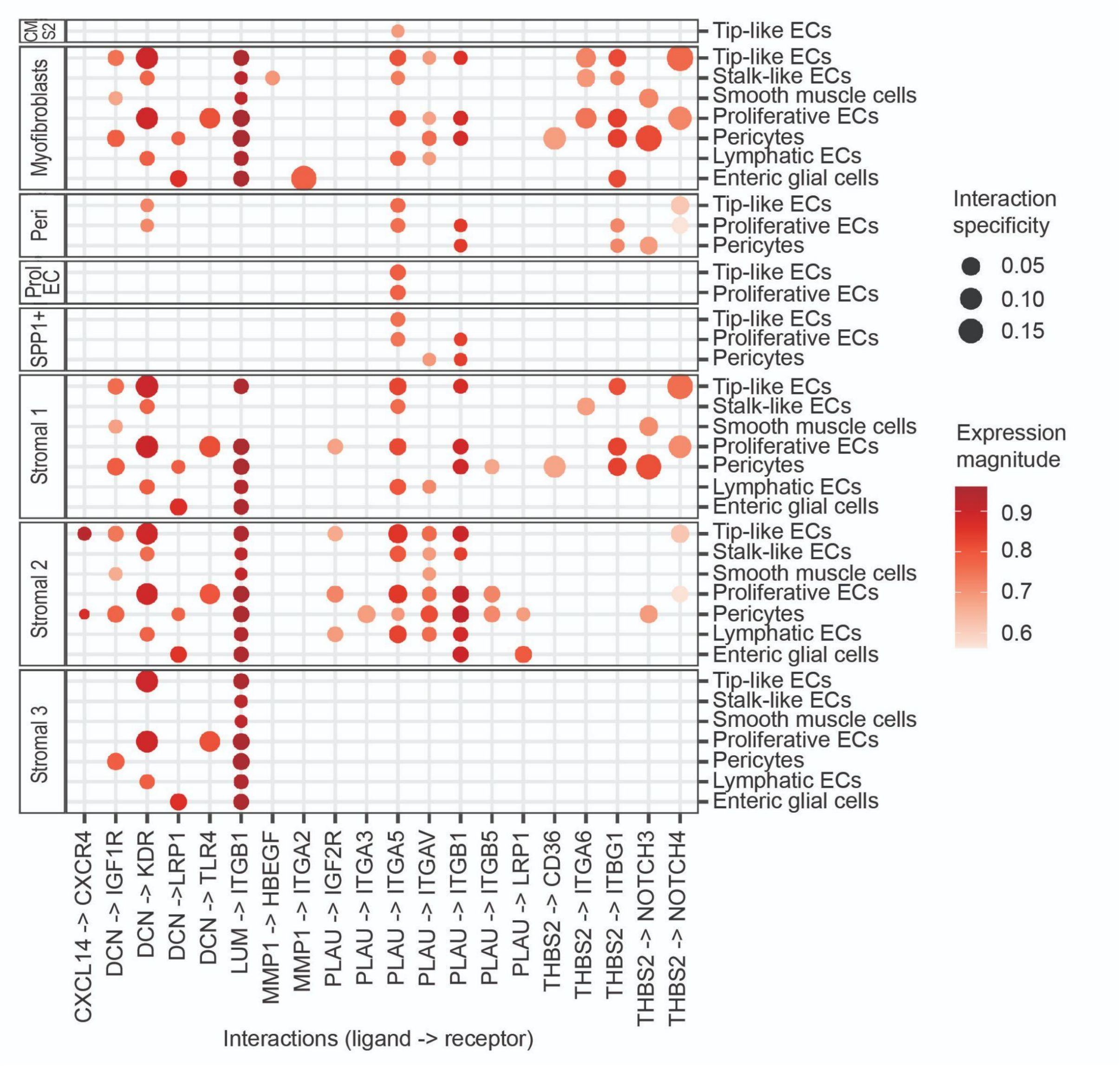


***Supplementary Figure S36:*** *Ligand-receptor interactions between the different cell types derived from scRNA-seq dataset from Lee et al. that overlap with the interactions predicted in our ST data. The left panel shows the source of the interaction (ligands) and the right the target (receptors), which in this case are the minor stromal cell populations.*

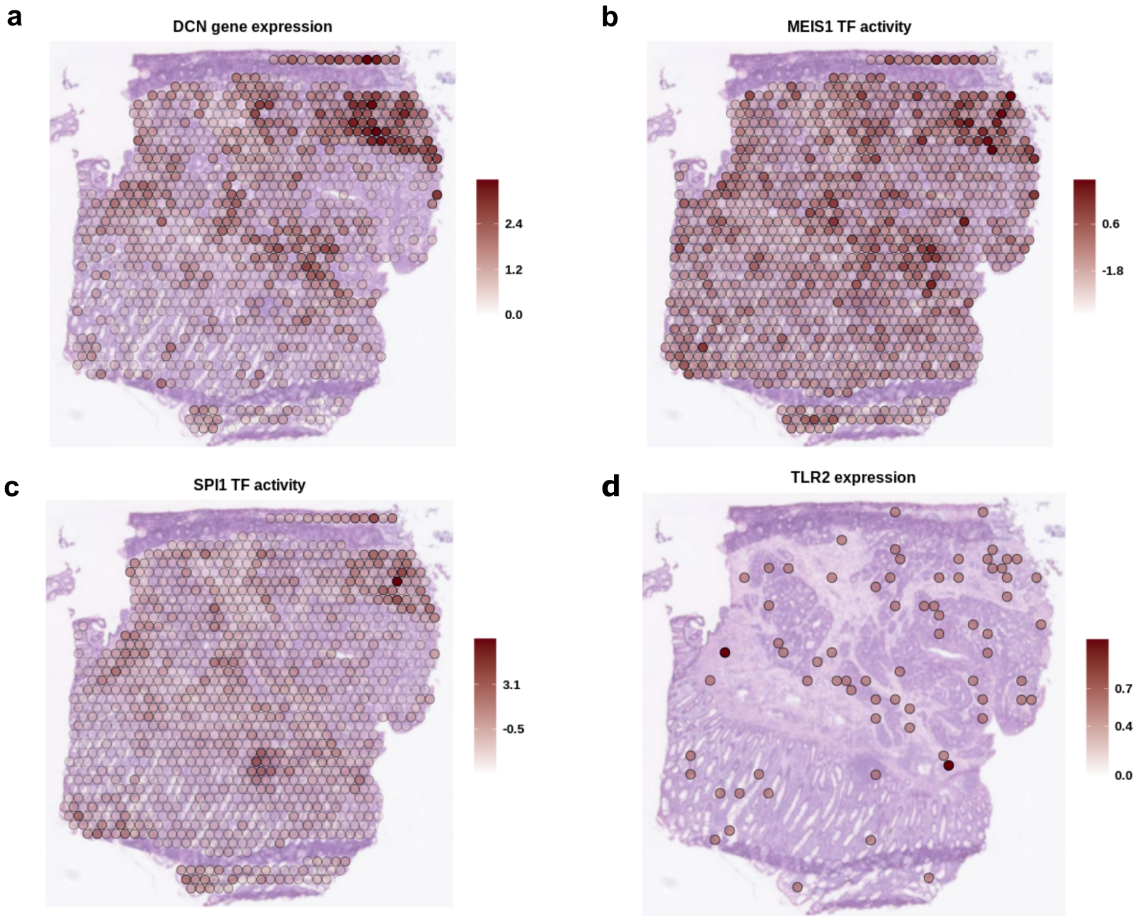


***Supplementary Figure S37:*** *Spatial mapping of: a) DCN gene expression, b) MEIS1 TF activity, c) SPI1 TF activity and d) TLR2 gene expression in the S2_Col_R_Rep1 sample.*

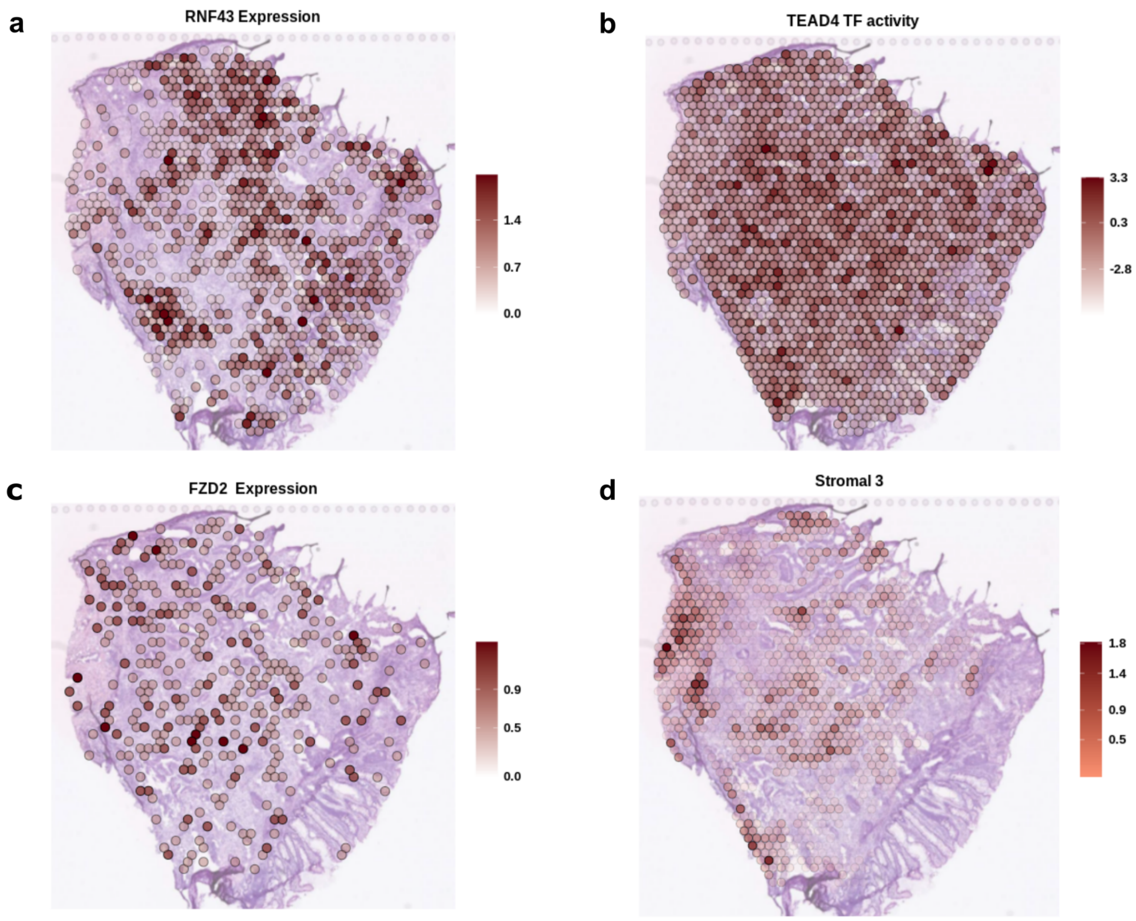


***Supplementary Figure S38:*** *Spatial mapping of: a) RNF43 gene expression, b) TEAD4 TF activity, c) FZD2 gene expression and d) estimated stromal 3 cell abundance in the S6_Rec_Rep2 sample.*

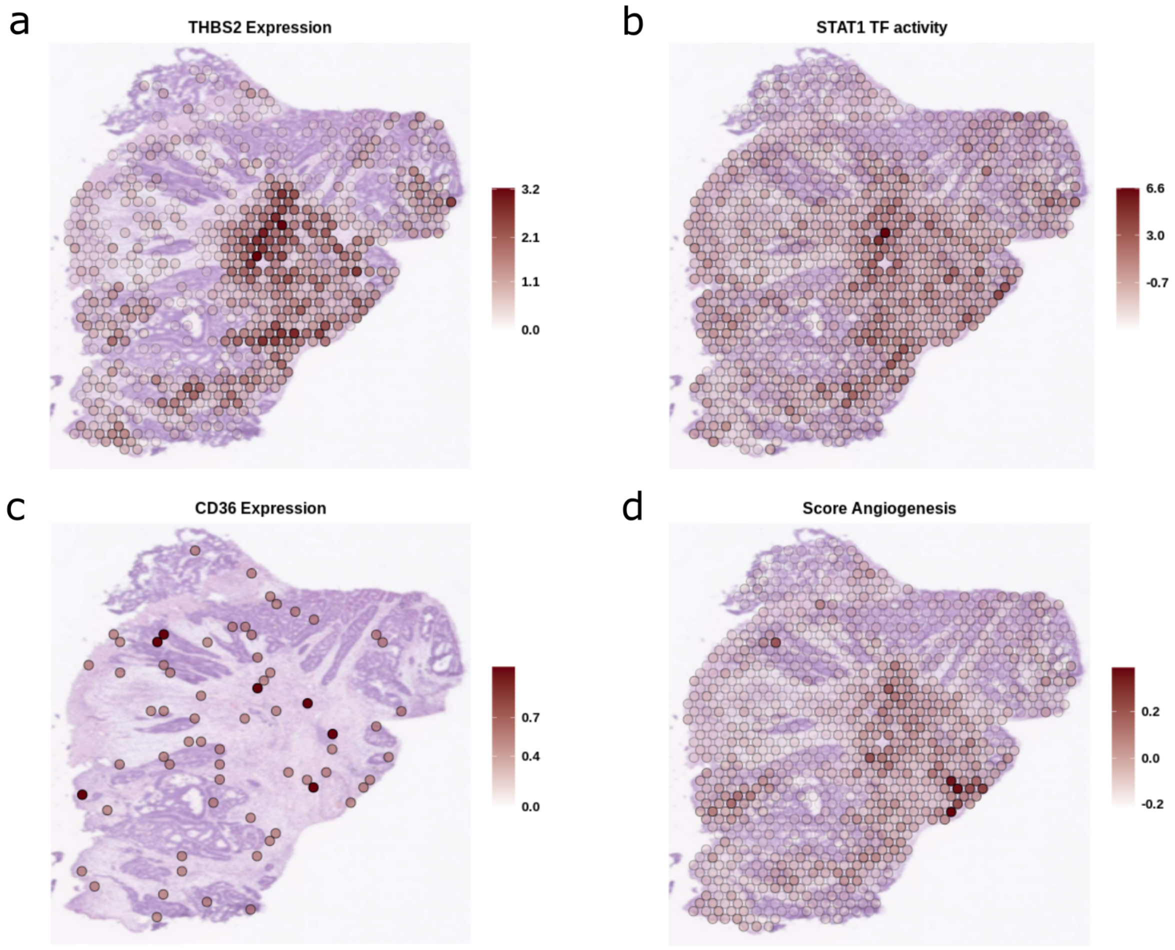


***Supplementary Figure S39:*** *Spatial mapping of: a) THBS2 gene expression, b) STAT1 TF activity, c) CD36 gene expression and d) the angiogenesis score in the S4_ Col_ Sig_ Rep2 sample.*

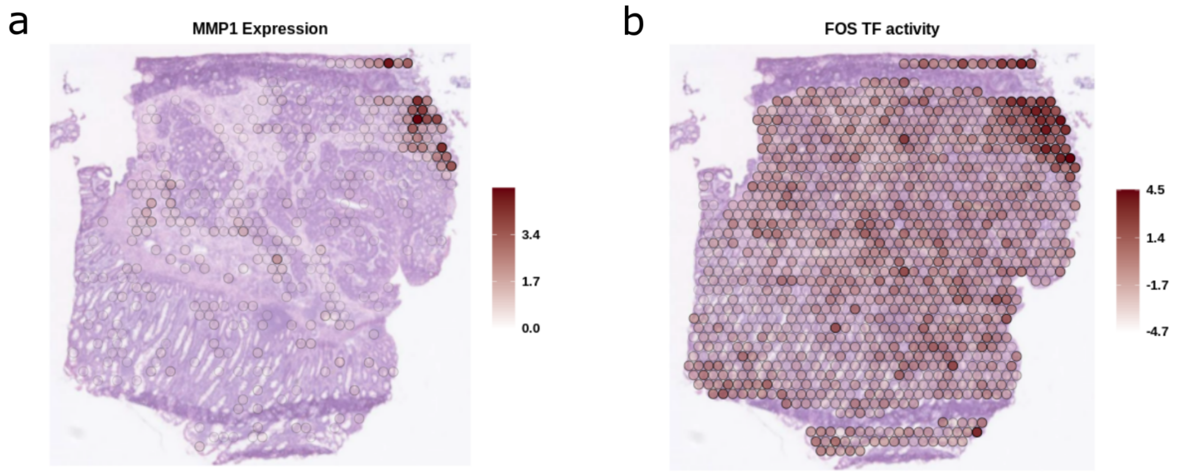


***Supplementary Figure S40:*** *Spatial mapping of: a) MMP1 gene expression and b) FOS TF activity in the S2_ Col_ R_ Rep1 sample.*

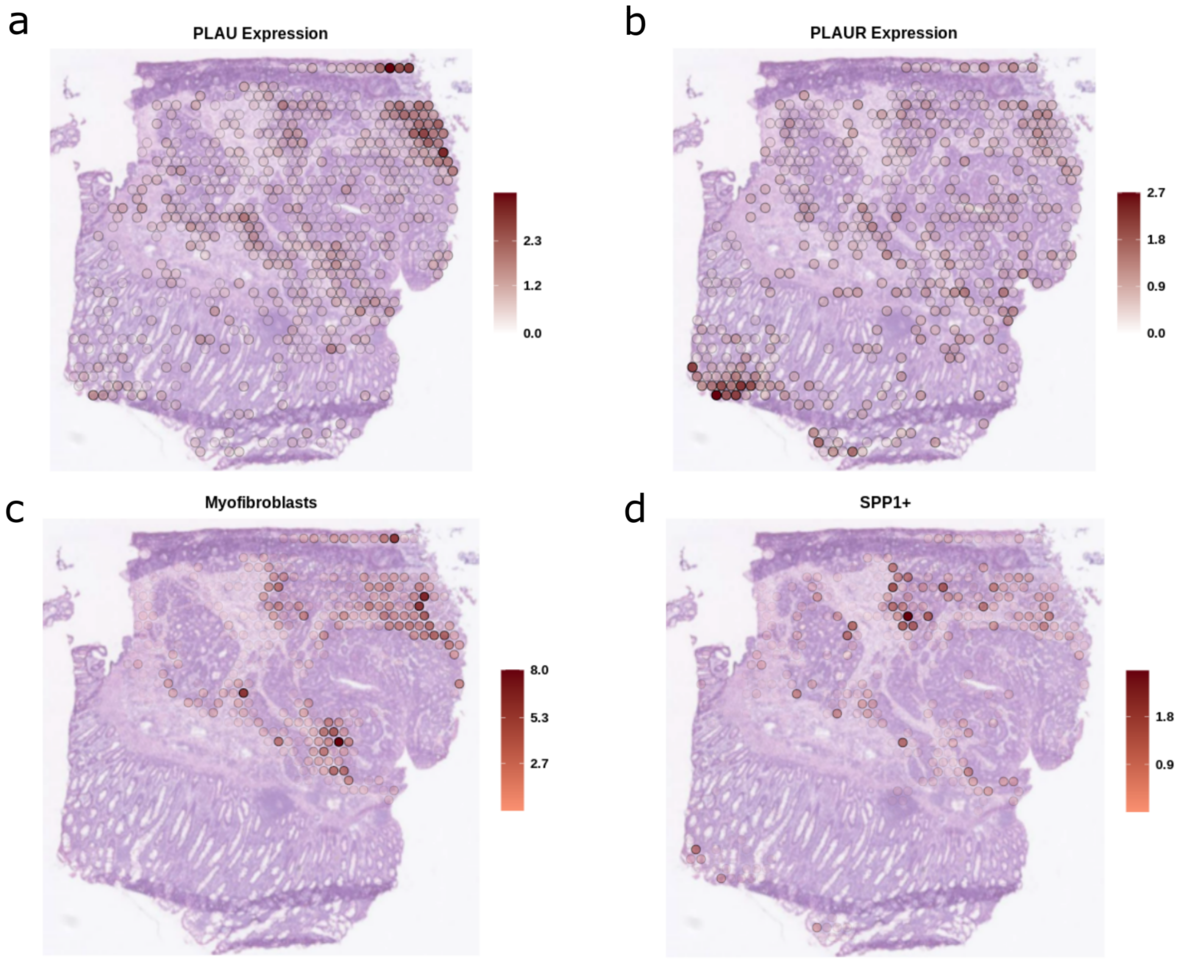


***Supplementary Figure S41:*** *Spatial mapping of a) PLAU gene expression, b) PLAUR gene expression, c) estimated myofibroblast cell abundance and d) estimated SPP1+ cell abundance in the S2_Col_R_Rep1 sample.*

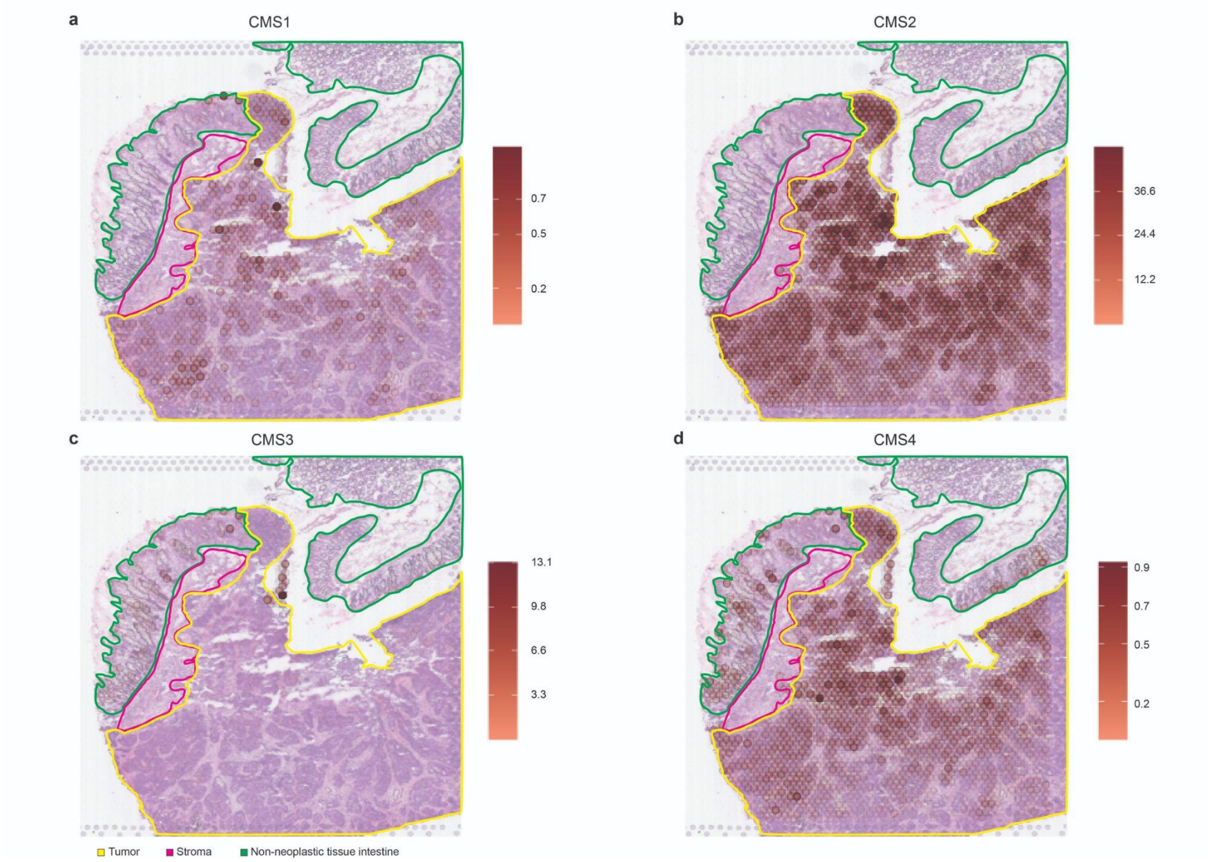


***Supplementary Figure S42:*** *Overlay of the spatial mapping of the abundance of the different CMS signatures according to the deconvolution results with the pathologists' tissue annotations for the ST-colon1_ Unt sample.*

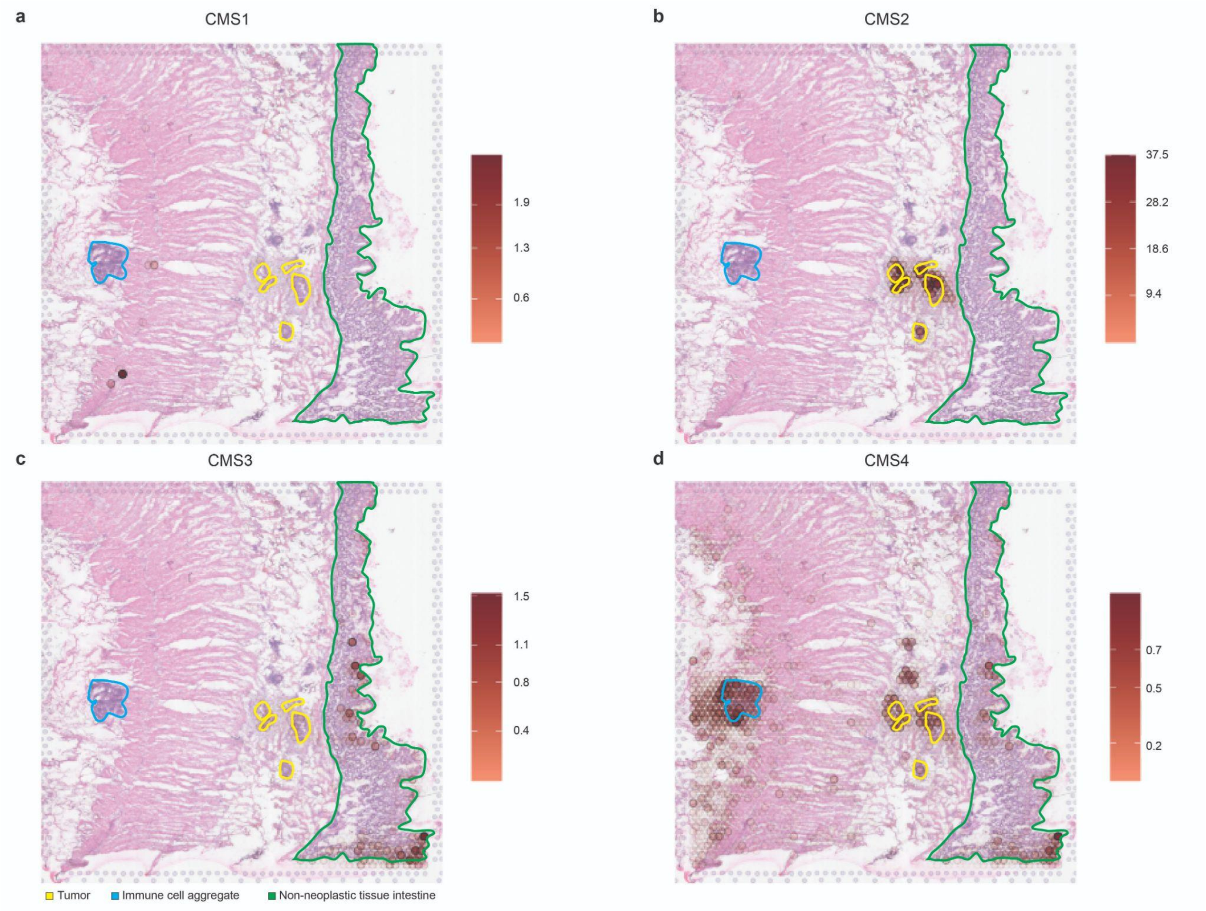


***Supplementary Figure S43:*** *Overlay of the spatial mapping of the abundance of the different CMS signatures according to the deconvolution results with the pathologists' tissue annotations for the ST-colon2_ Unt sample.*

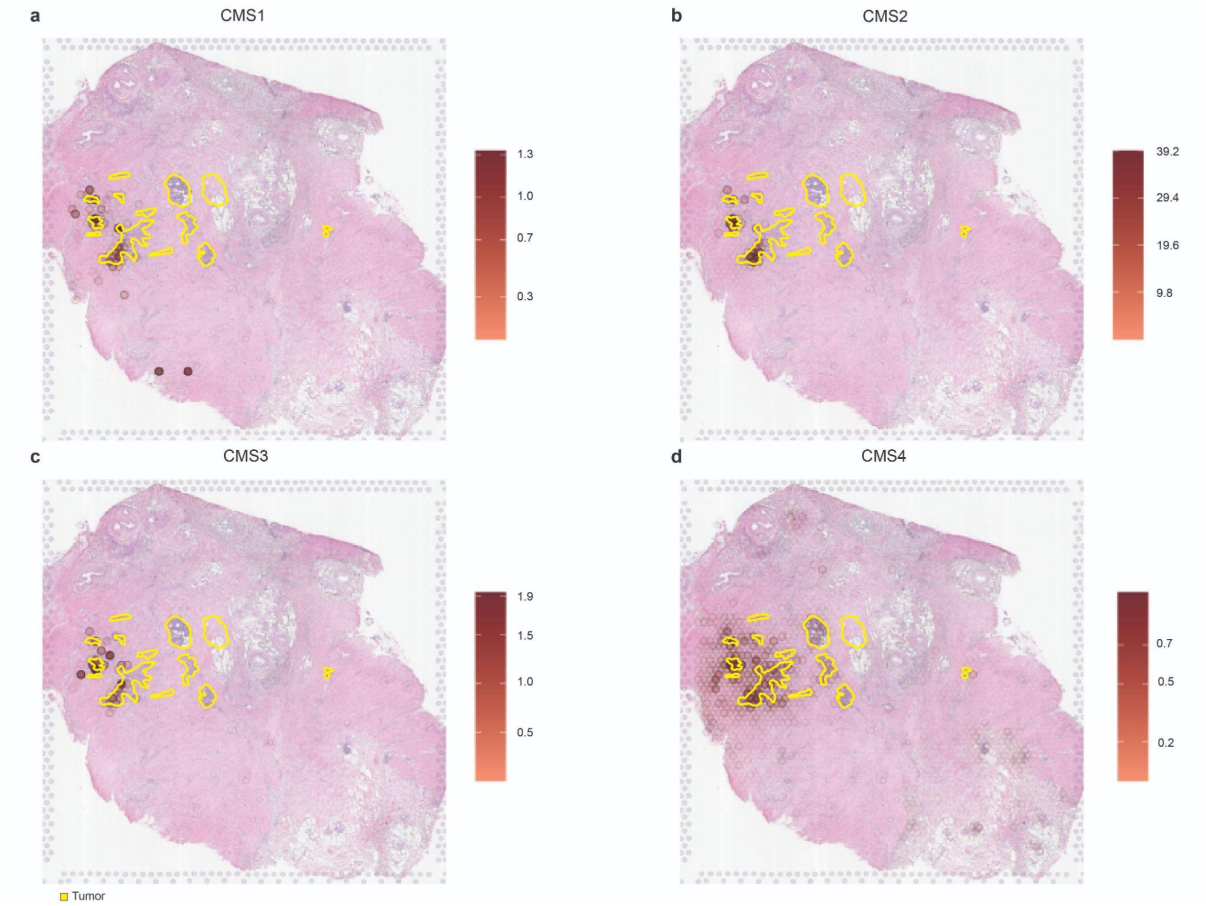


***Supplementary Figure S44:*** *Overlay of the spatial mapping of the abundance of the different CMS signatures according to the deconvolution results with the pathologists' tissue annotations for the ST-colon3_ Tre sample.*

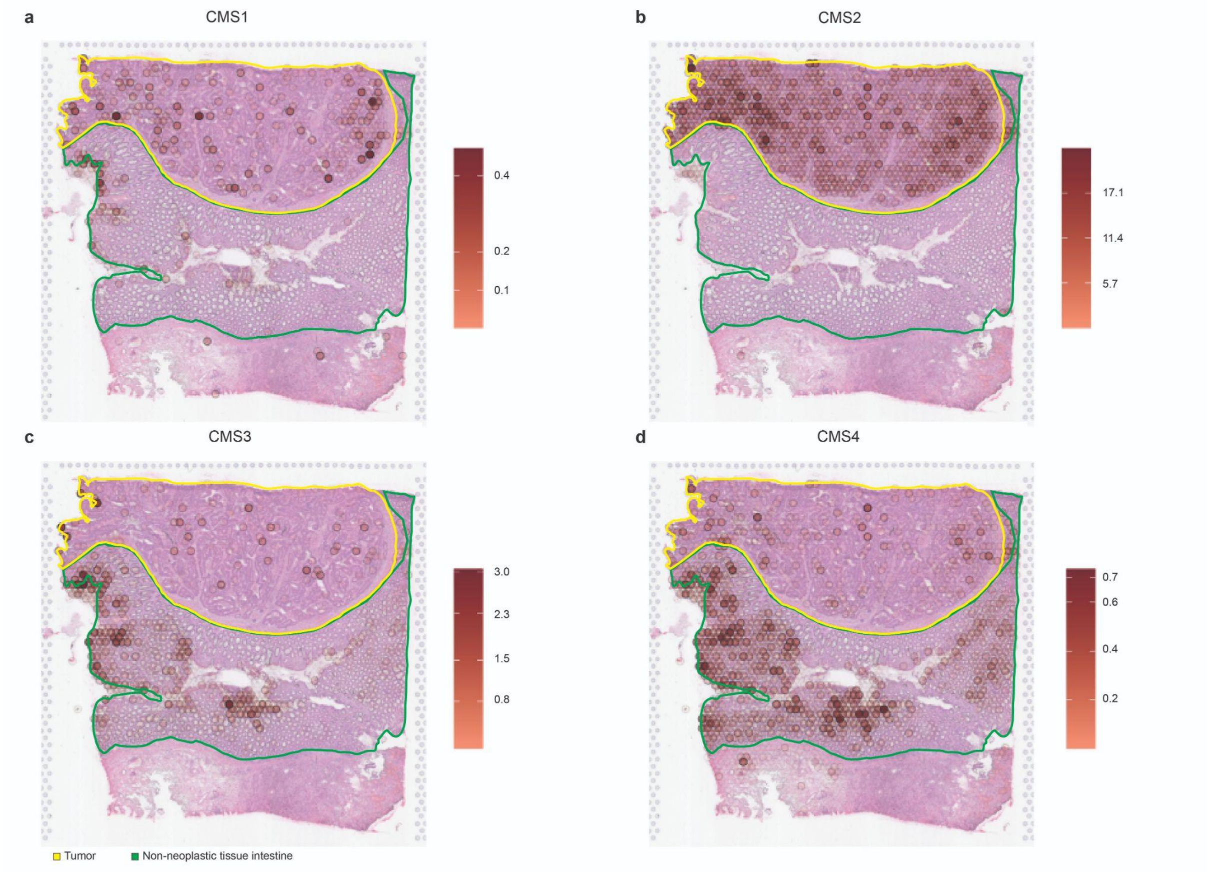


***Supplementary Figure S45:*** *Overlay of the spatial mapping of the abundance of the different CMS signatures according to the deconvolution results with the pathologists' tissue annotations for the ST-colon4_ Tre sample.*

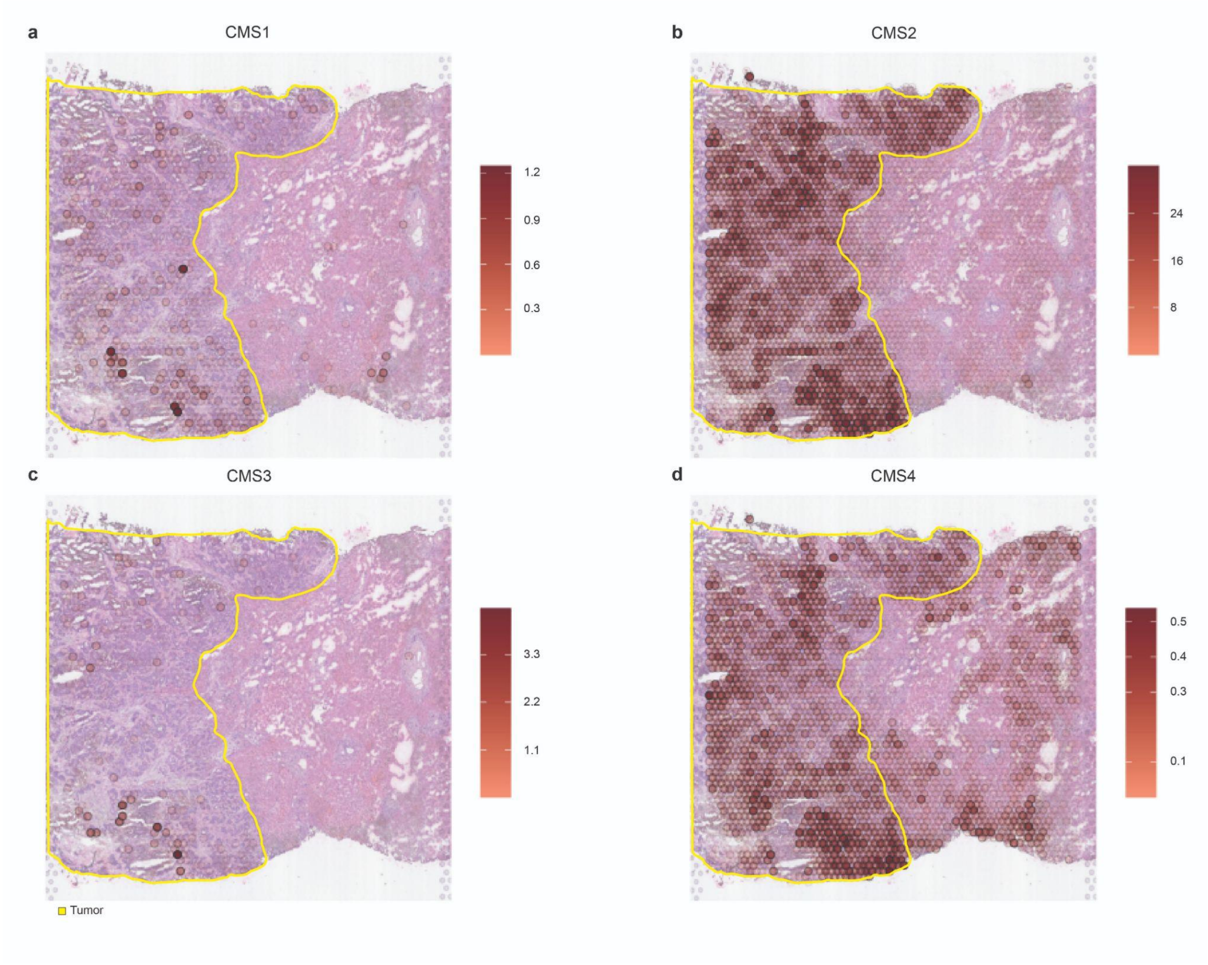


***Supplementary Figure S46:*** *Overlay of the spatial mapping of the abundance of the different CMS signatures according to the deconvolution results with the pathologists' tissue annotations for the ST-liver1_ Unt sample.*

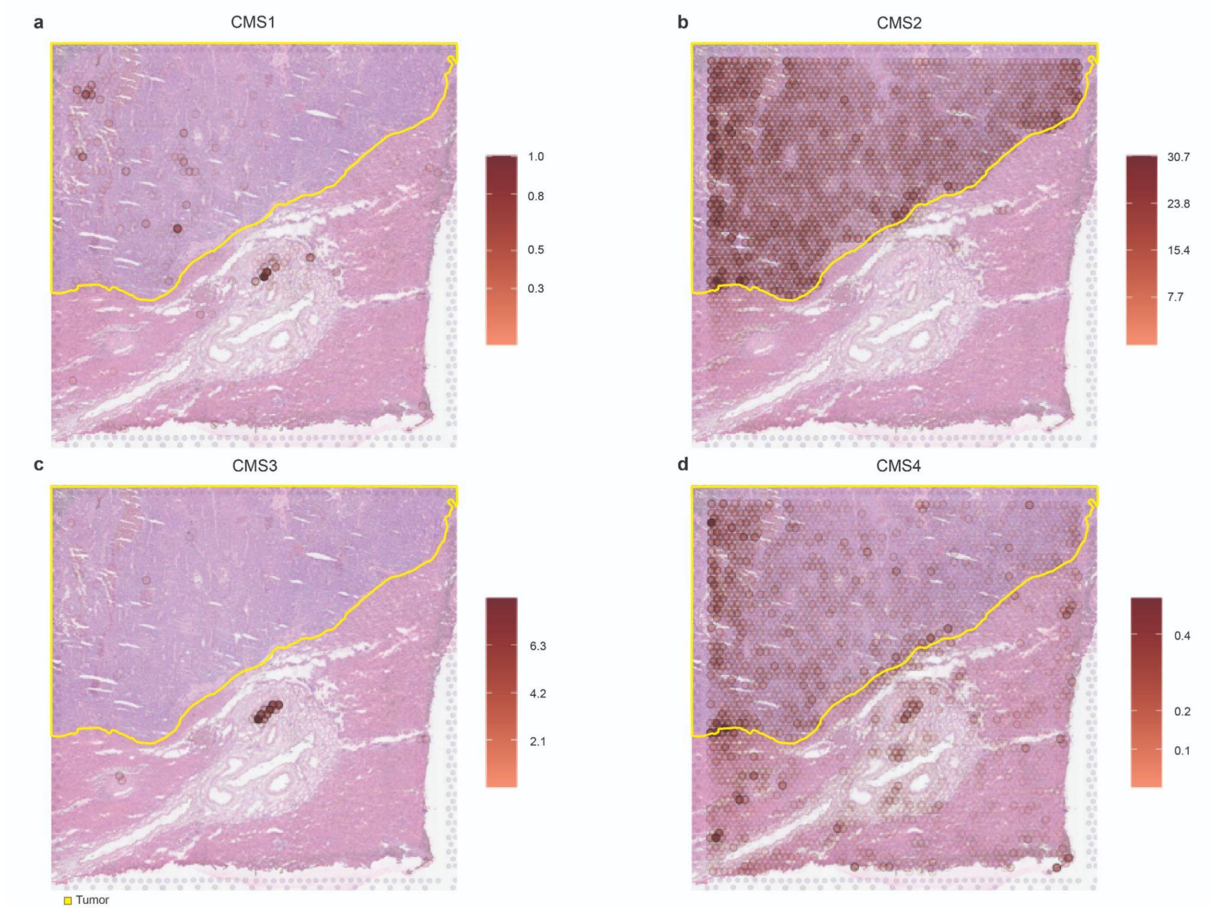


***Supplementary Figure S47:*** *Overlay of the spatial mapping of the abundance of the different CMS signatures according to the deconvolution results with the pathologists' tissue annotations for the ST-liver2_Unt sample.*

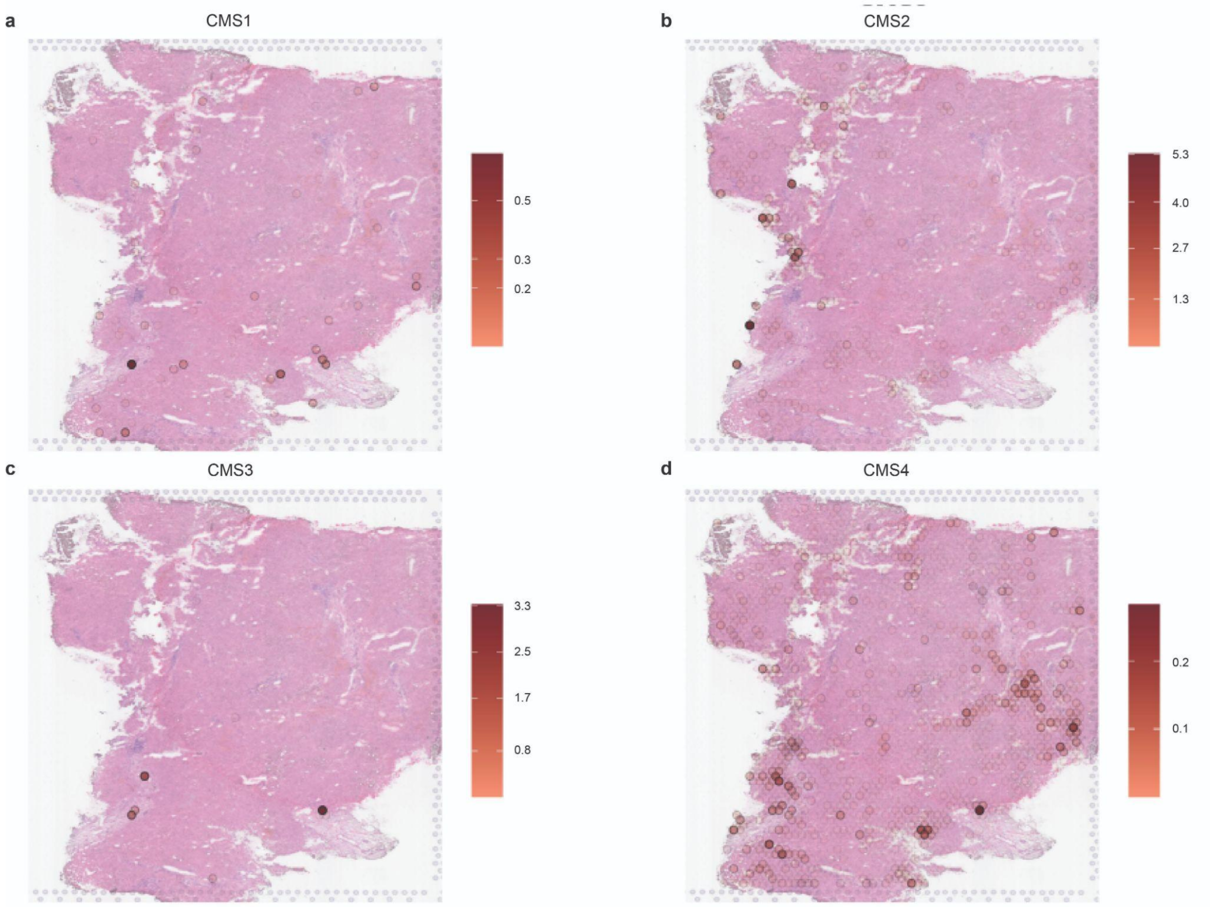


***Supplementary Figure S48:*** *Overlay of the spatial mapping of the abundance of the different CMS signatures according to the deconvolution results with the pathologists' tissue annotations for the ST-liver3_Tre sample.*

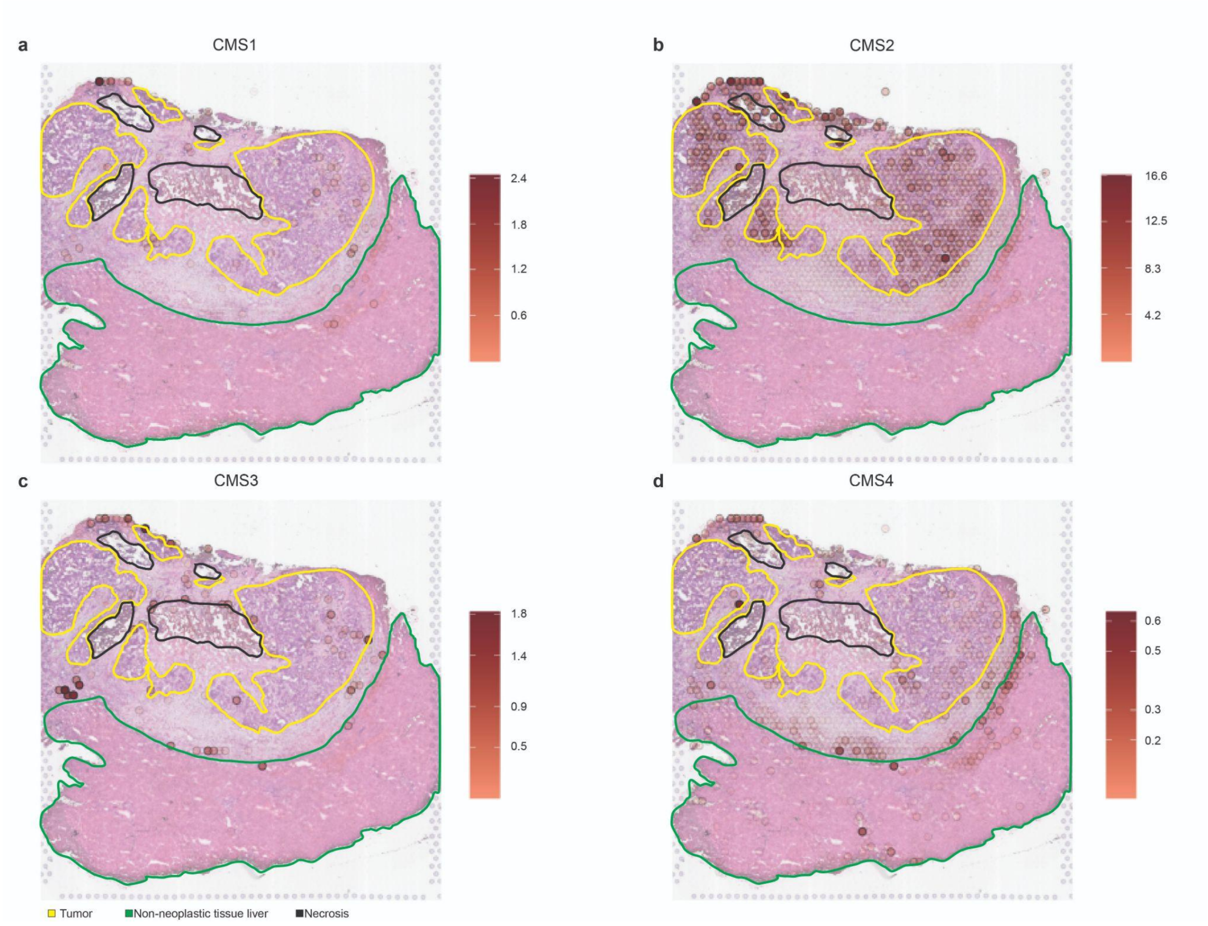


***Supplementary Figure S49:*** *Overlay of the spatial mapping of the abundance of the different CMS signatures according to the deconvolution results with the pathologists' tissue annotations for the ST-liver4_ Tre sample.*

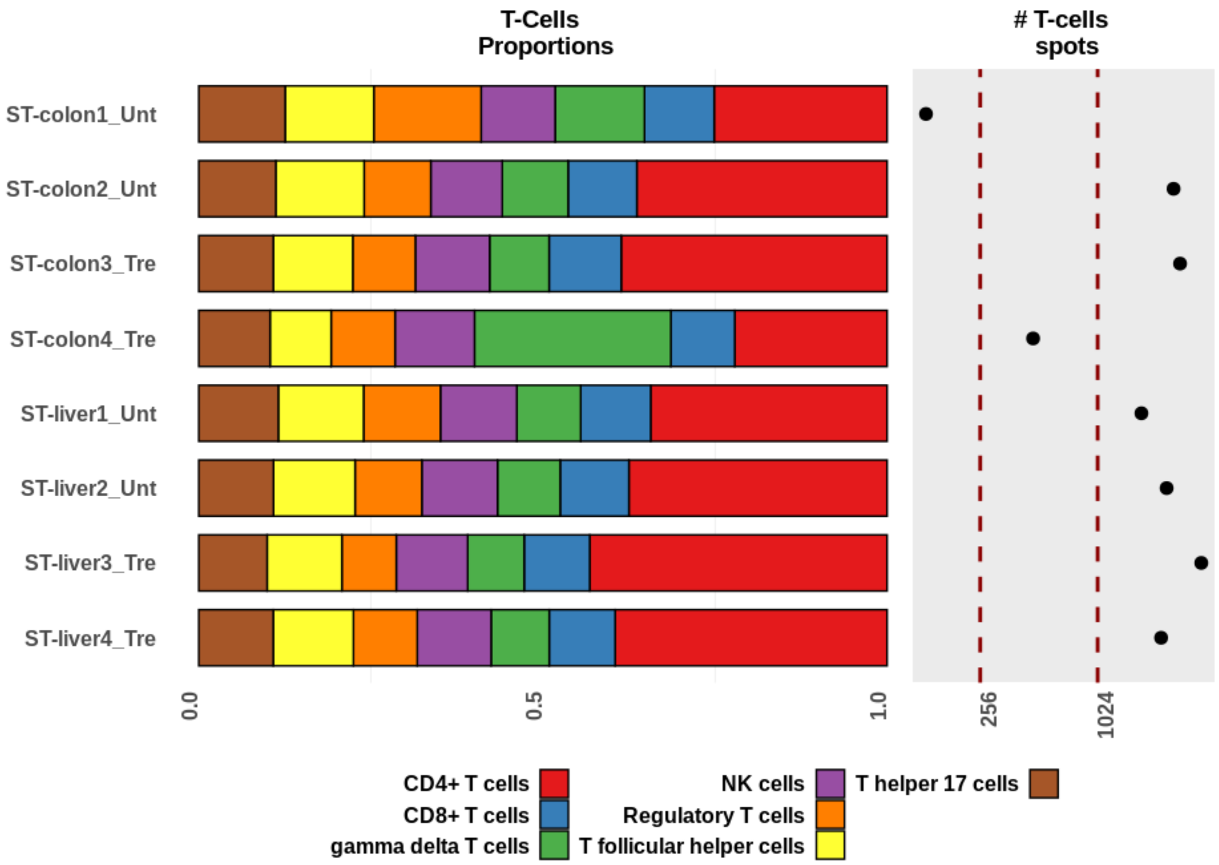


***Supplementary Figure S50:*** *Per sample proportions of the T-cells subtypes as estimated by the results of the deconvolution approach. The number of spots containing an abundance of at least 20% of T-cells is also displayed.*

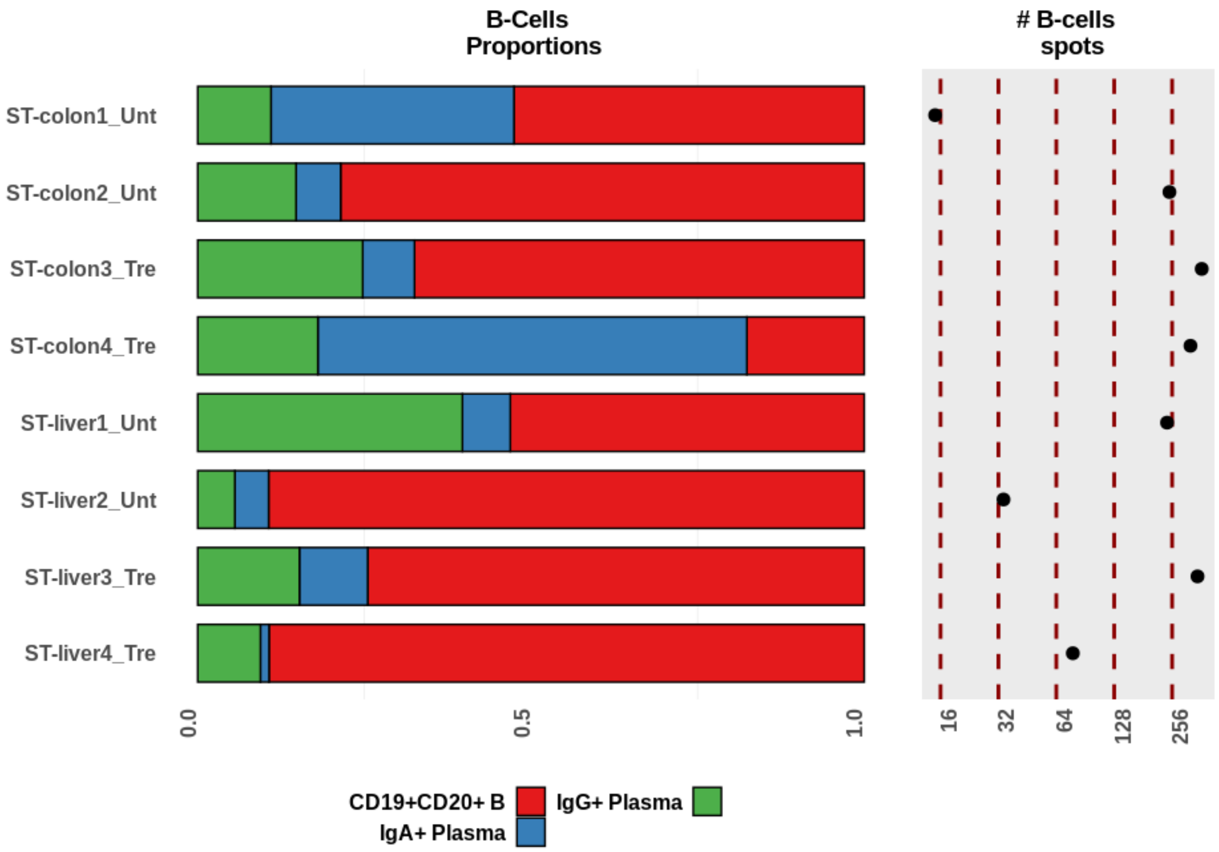


***Supplementary Figure S51:*** *Per sample proportions of the B-cells subtypes as estimated by the results of the deconvolution approach. The number of spots containing an abundance of at least 20% of B-cells is also displayed.*

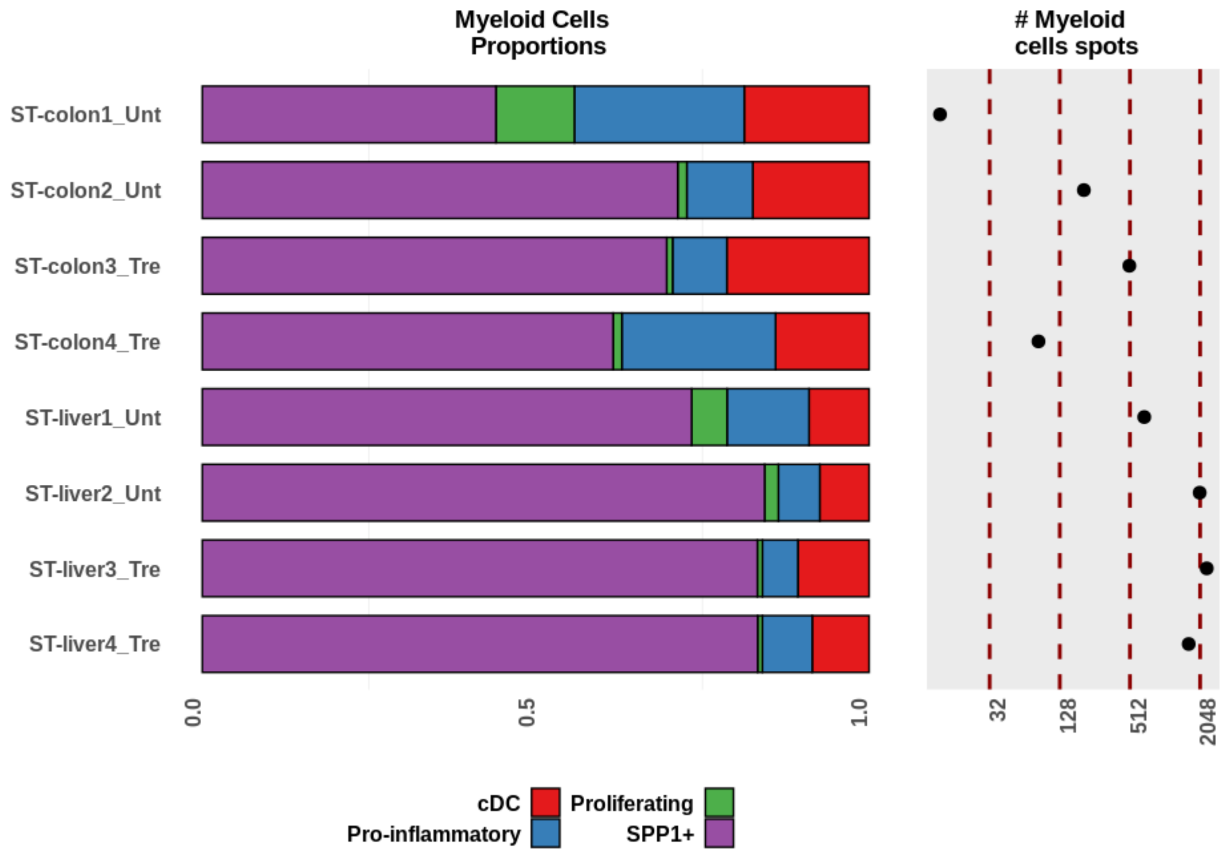


***Supplementary Figure S52:*** *Per sample proportions of the myeloid cell subtypes as estimated by the results of the deconvolution approach. The number of spots containing an abundance of at least 20% of myeloid cells is also displayed.*

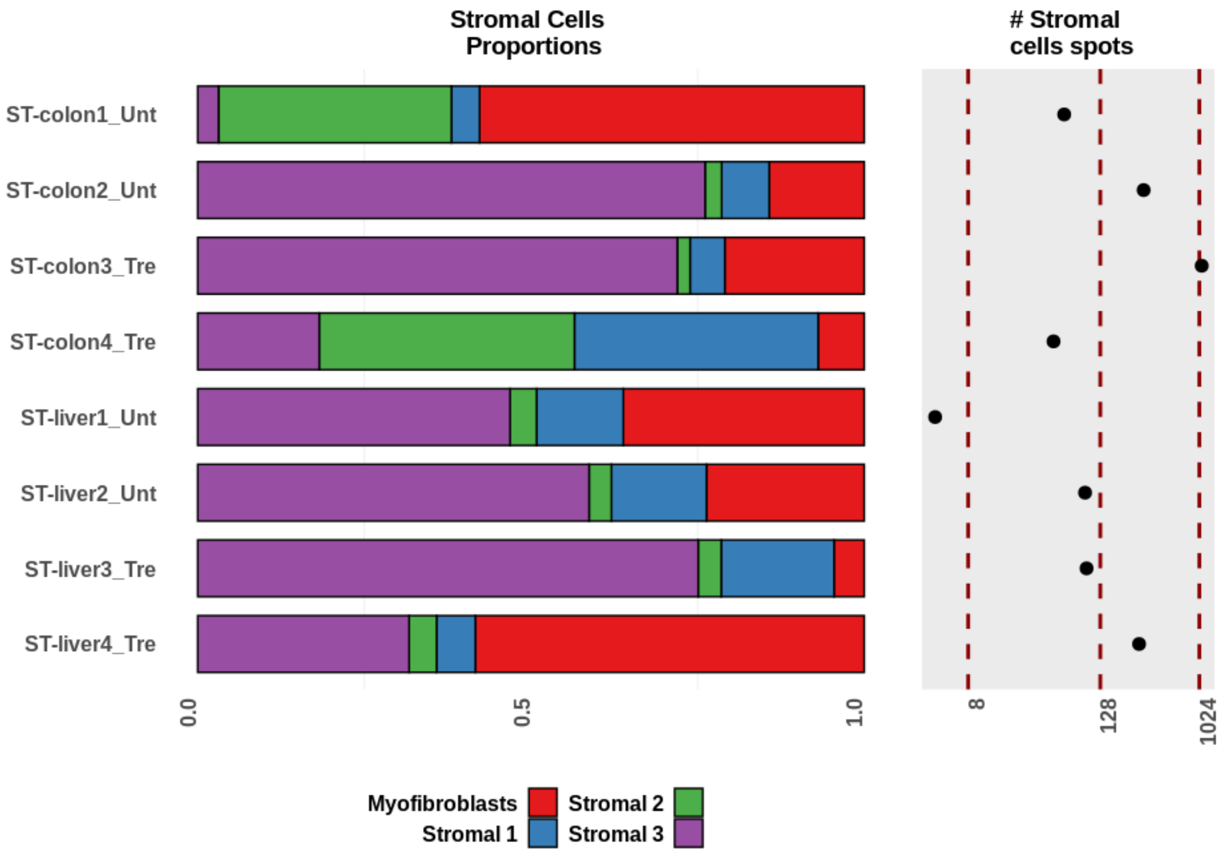


***Supplementary Figure S53:*** *Per sample proportions of the main stromal cell subtypes as estimated by the results of the deconvolution approach. The number of spots containing an abundance of at least 20% of stromal cells is also displayed.*

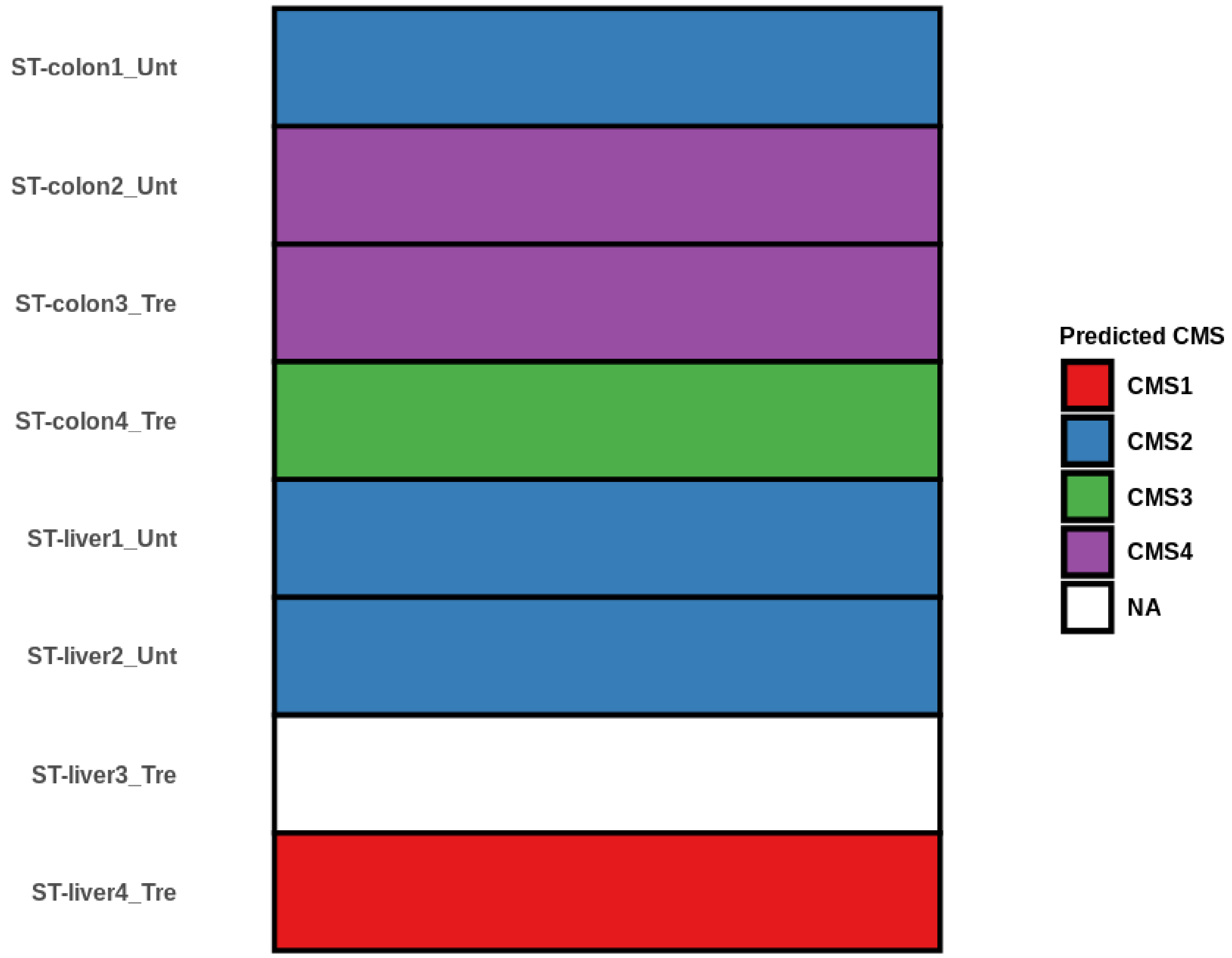


***Supplementary Figure S54:*** *Predicted CMS classification resulting from applying CMScaller on pseudo-bulk RNA-seq generated by pooling together all the spots from the external ST CRC dataset.*

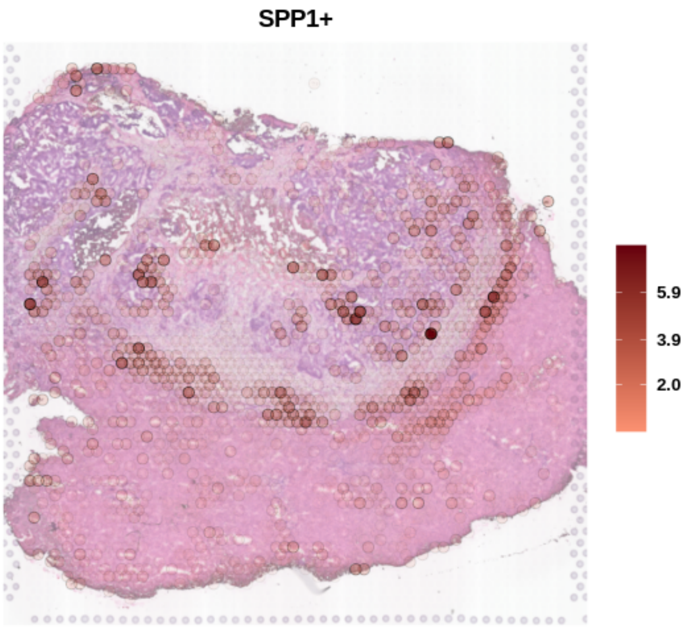


***Supplementary Figure S55:*** *Spatial mapping of the abundance of SPP1+ macrophages as predicted by the deconvolution for the ST-liver4_ Tre sample.*

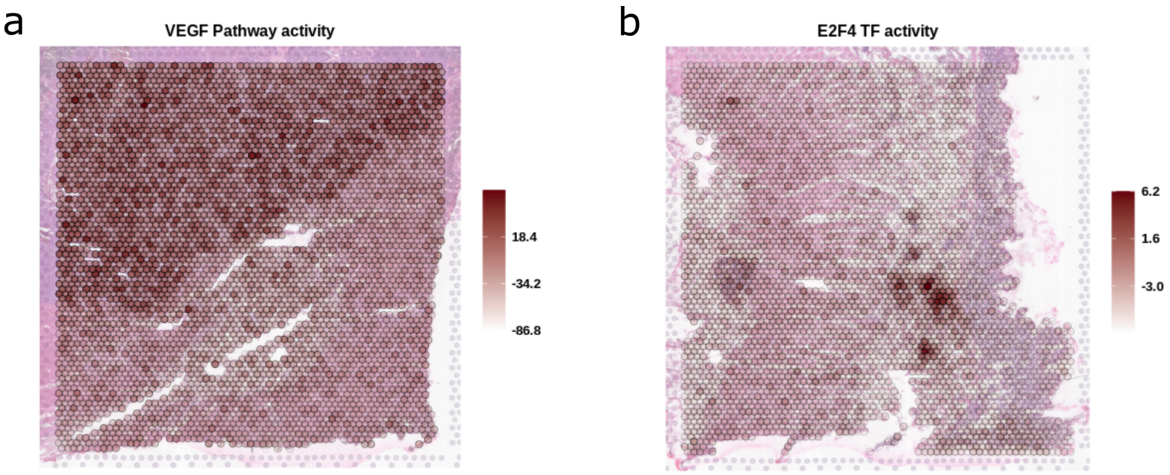


***Supplementary Figure S56:*** *Spatial mapping of: a) the VEGF pathway activity in the ST-liver2_ Unt sample, and b) the E2F4 TF activity in the ST-colon2_ Unt sample.*

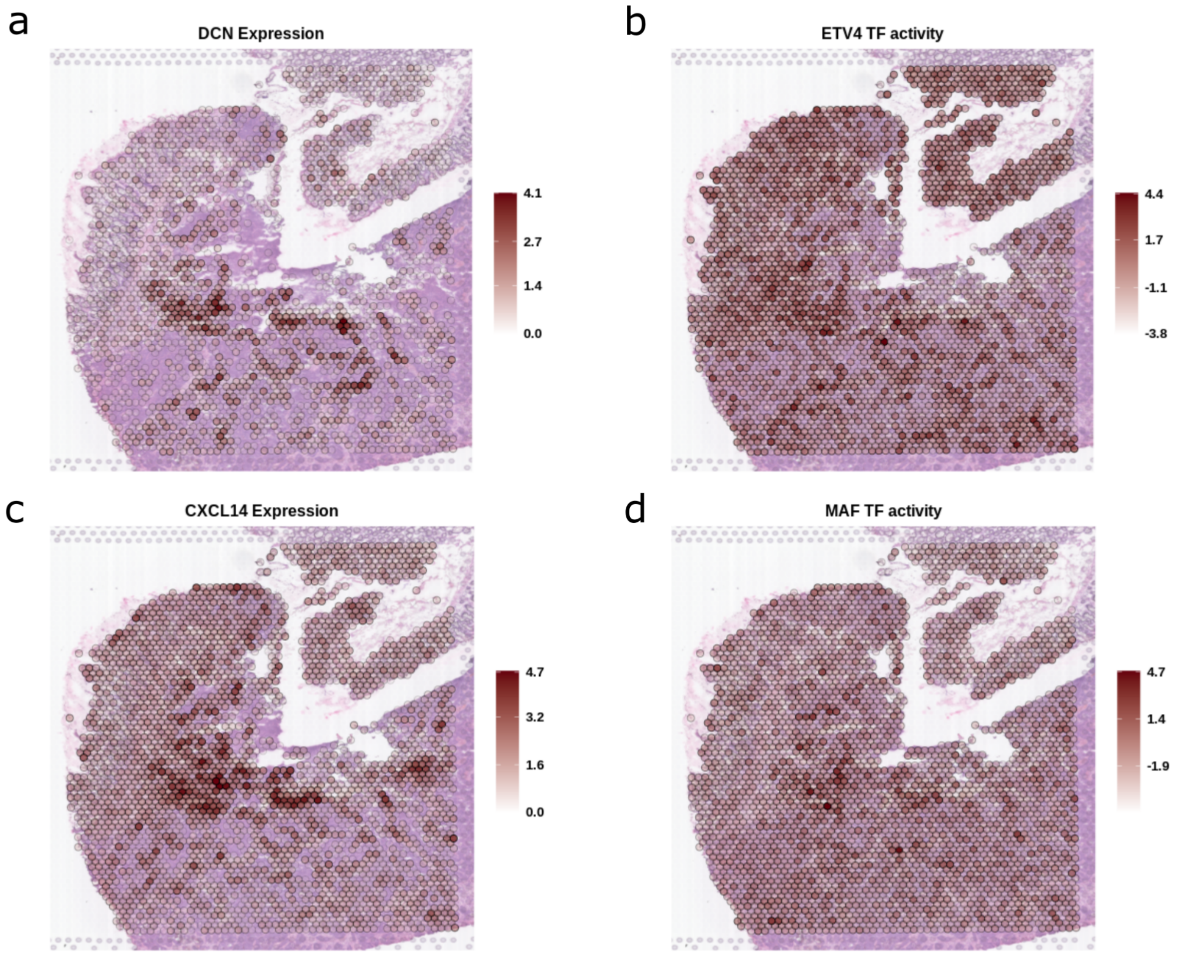


***Supplementary Figure S57:*** *Spatial mapping of: a) DCN gene expression, b) ETV4 TF activity, c) CXCL14 gene expression and d) MAS TF activity in the ST-colon1_ Unt sample.*

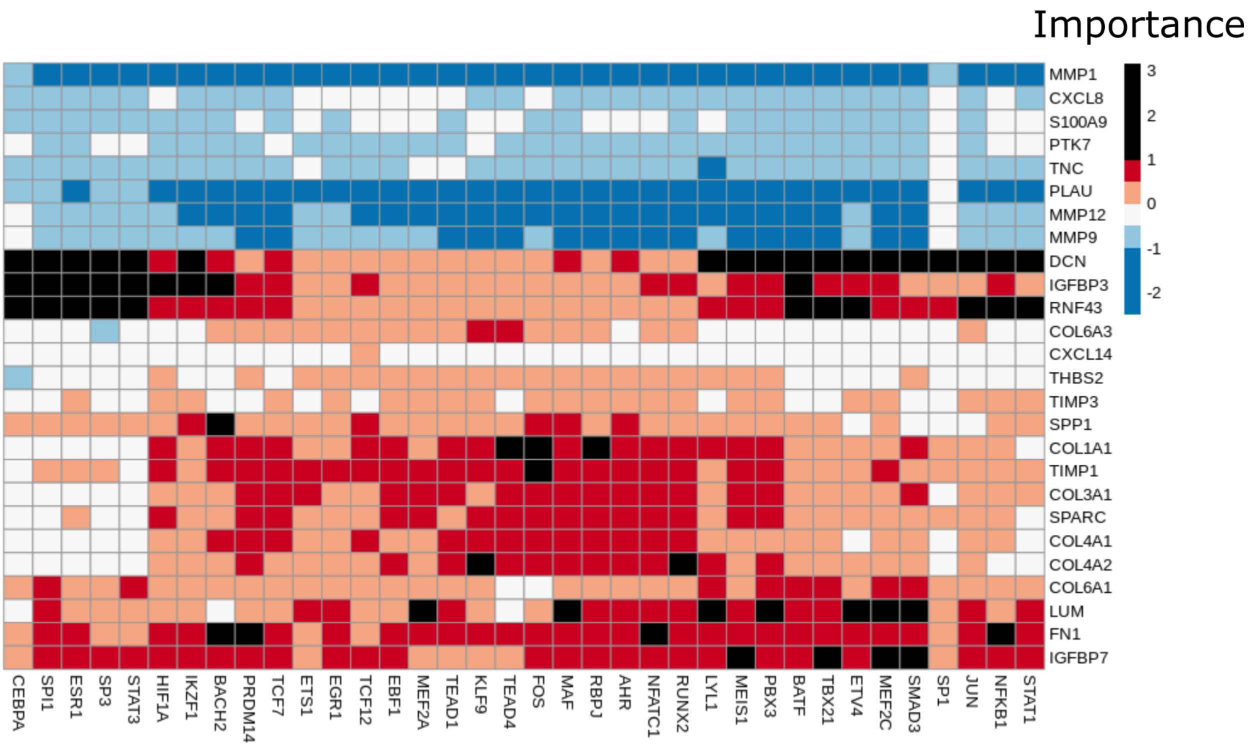


***Supplementary Figure S58:*** *Misty results showing the potential importance of ligands (rows) expression on TF (columns) activity when considering the samples with hepatic metastasis of CRC tumors. The ligand-TFs relationships with an importance score over 1 are represented as black slots and were considered as relevant.*

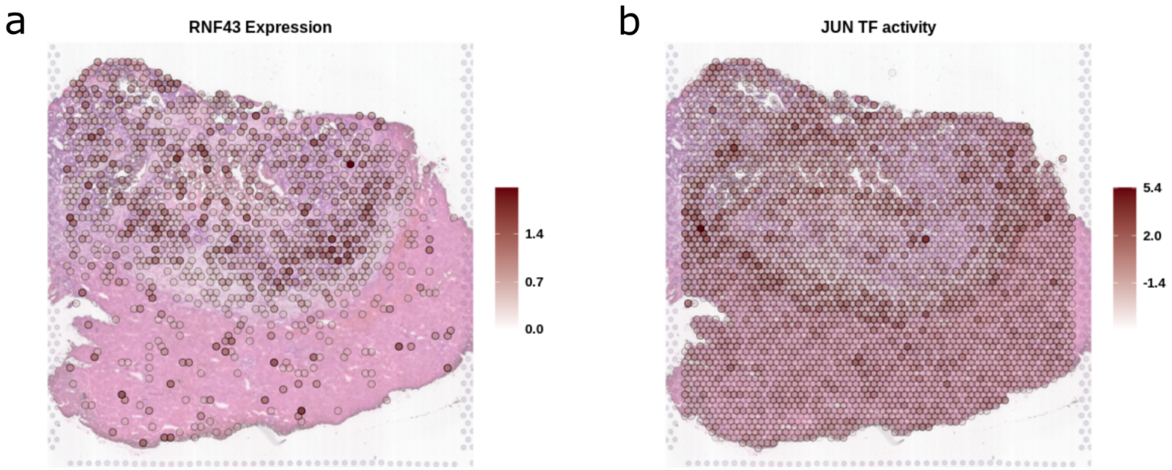


***Supplementary Figure S59:*** *Spatial mapping of: a) RNF43 gene expression and b) JUN TF activity in the ST-liver4_ Tre sample.*

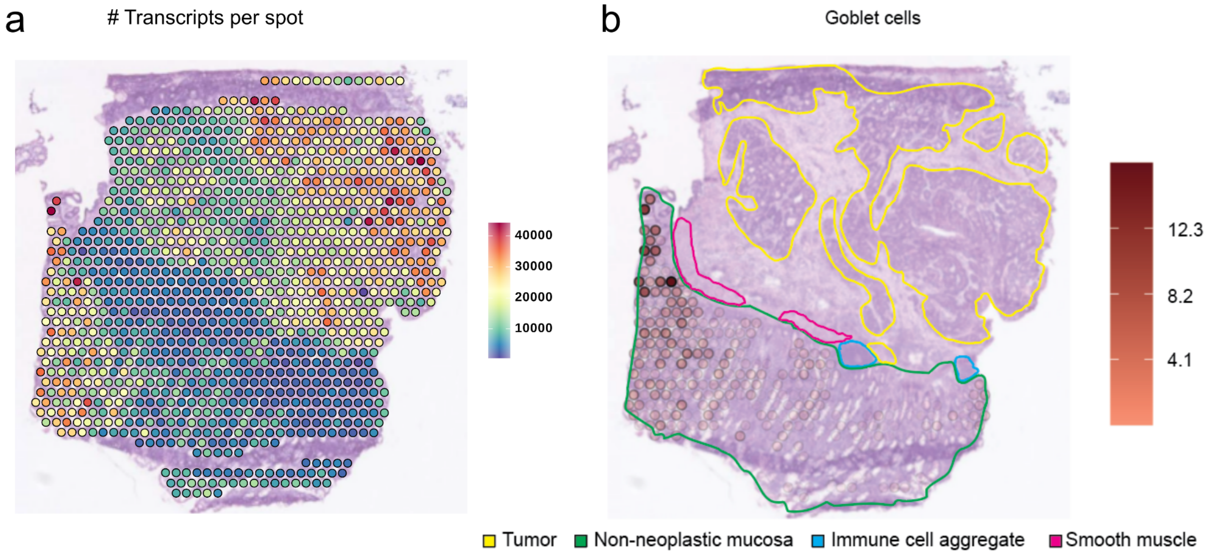


***Supplementary Figure S60:*** *Spatial mapping of a) the number of transcript per spot, and b) the predicted abundance of goblet cells for the S2_Col_R_Rep1 sample. It is to note that the regions of the non-neoplastic mucosa showing a lower number of transcripts per spot were accordingly predicted to have a reduced amount of goblet cells. Pathologists' tissue annotations are also displayed for clarification.*

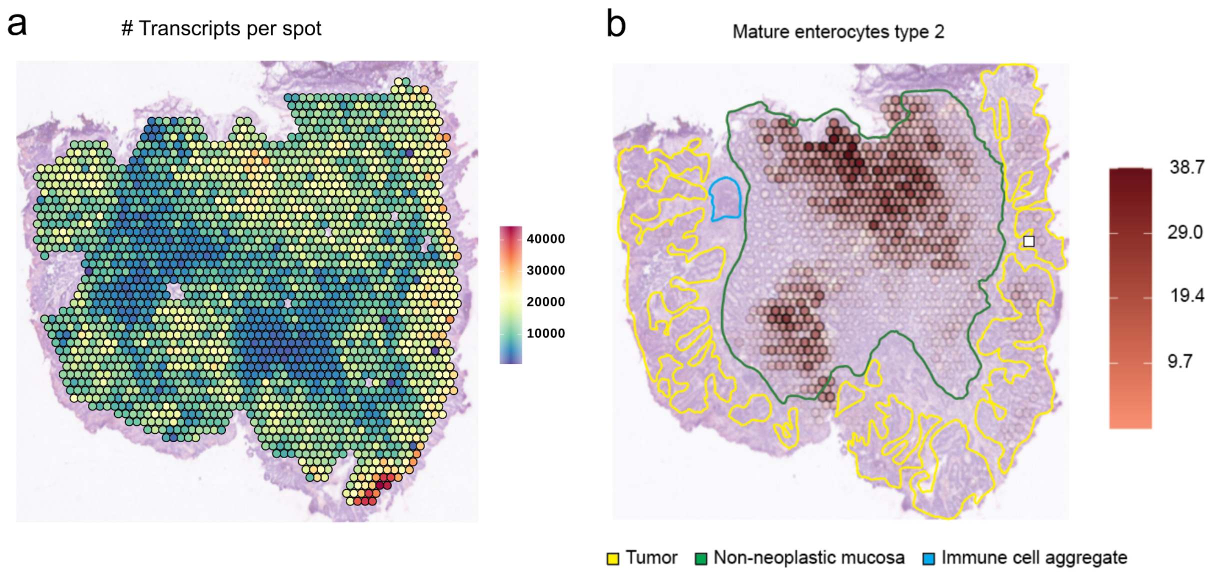


***Supplementary Figure S61:*** *Spatial mapping of: a) the number of transcript per spot, and b) the predicted abundance of mature enterocytes type 2 cells for the S5_Rec_Rep1 sample. It is to note that the regions of the non-neoplastic mucosa showing a lower number of transcripts per spot were accordingly predicted to have a reduced amount of mature enterocytes type 2. Pathologists' tissue annotations are also displayed for clarification.*

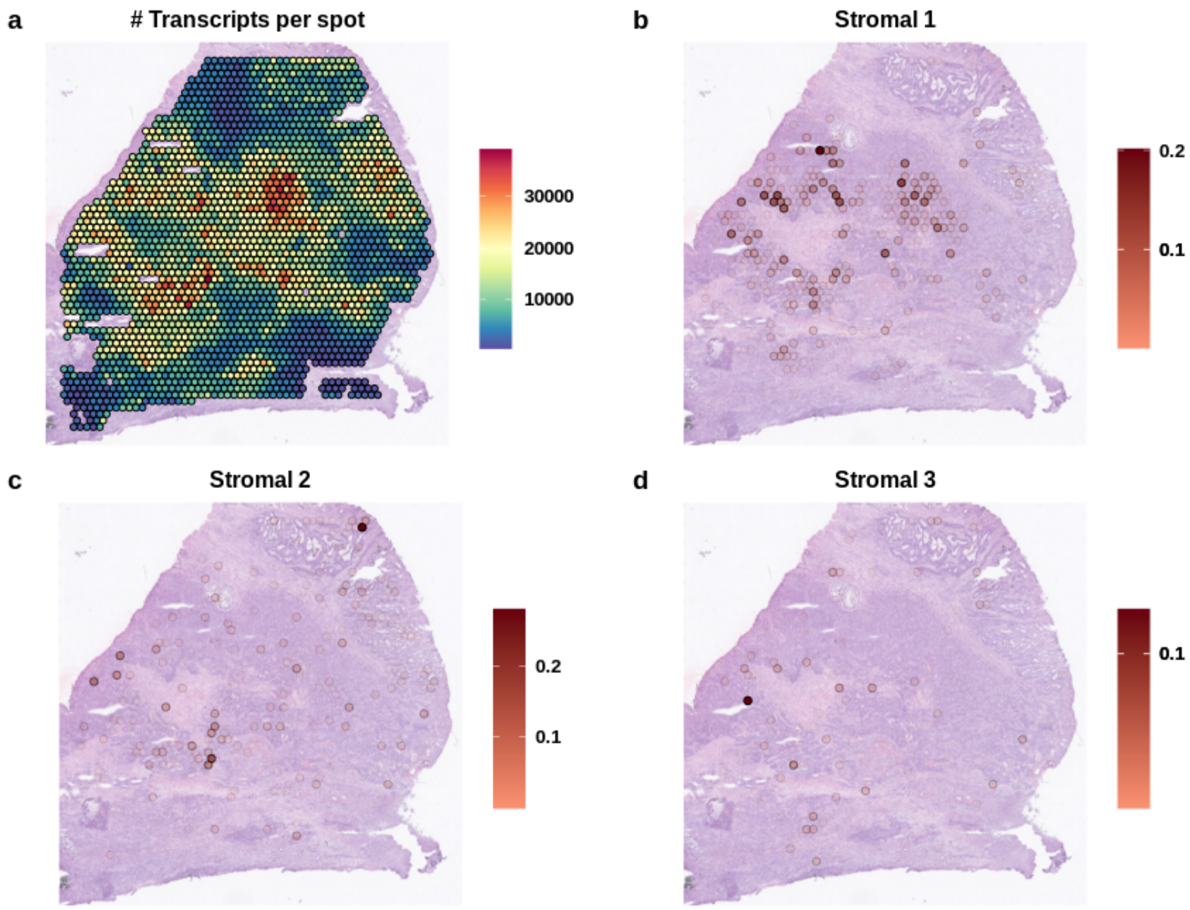


***Supplementary Figure S62:*** *Spatial mapping of the number of transcript per spot (a) and the predicted abundance of the different stromal main populations, excluding myofibroblasts, (b-d) for the S3_Col_Rep1 sample. The regions with lower number of transcripts match with stromal morphological features, however non stromal abundance was mapped into those areas.*

### 2.Supplementary Tables

| *Cell type* | *Cell subtype* | *Comments* | *Number of Cells* | *Genes used for deconvolution* |
|---|---|---|---|---|
| *B cells* | *CD19+CD20+ B* | *B lymphocytes* | *3733* | *38* |
| *B cells* | *IgA+ Plasma* | *B lymphocytes* | *7305* | *7* |
| *B cells* | *IgG+ Plasma* | *B lymphocytes* | *6404* | *15* |
| *Epithelial cells* | *CMS1* | *Tumor cells* | *1201* | *412* |
| *Epithelial cells* | *CMS2* | *Tumor cells* | *10771* | *620* |
| *Epithelial cells* | *CMS3* | *Tumor cells* | *5486* | *228* |
| *Epithelial cells* | *CMS4* | *Tumor cells* | *11* | *22* |
| *Epithelial cells* | *Goblet cells* | *Normal epithelial cells* | *305* | *176* |
| *Epithelial cells* | *Intermediate* | *Normal epithelial cells* | *139* | *131* |
| *Epithelial cells* | *Mature Enterocytes type 1* | *Normal epithelial cells* | *3546* | *182* |
| *Epithelial cells* | *Mature Enterocytes type 2* | *Normal epithelial cells* | *1755* | *172* |
| *Epithelial cells* | *Stem-like/TA* | *Normal epithelial cells* | *181* | *14* |
| *Mast cells* | *Mast cells* | *Normal epithelial cells* | *108* | *186* |
| *Myeloids* | *cDC* | *Conventional dendritic cells* | *187* | *43* |
| *Myeloids* | *Pro-inflammatory* | *Pro-inflammatory Macrophages* | *213* | *77* |
| *Myeloids* | *Proliferating* | *Proliferating Macrophages* | *131* | *73* |
| *Myeloids* | *SPP1+* | *SPP1+ Macrophages* | *1155* | *305* |
| *Stromal cells* | *Enteric glial cells* | | *2399* | *55* |

| | | | | |
|---|---|---|---|---|
| *Stromal cells* | *Lymphatic ECs* | *Lymphatic Endothelial Cells* | *171* | *239* |
| *Stromal cells* | *Myofibroblasts* | | *44* | *521* |
| *Stromal cells* | *Pericytes* | | *3097* | *32* |
| *Stromal cells* | *Proliferative ECs* | *Proliferative Endothelial Cells* | *3252* | *168* |
| *Stromal cells* | *Smooth muscle cells* | | *214* | *151* |
| *Stromal cells* | *Stalk-like ECs* | *Stalk-like Endothelial Cells* | *438* | *99* |
| *Stromal cells* | *Stromal 1* | | *406* | *618* |
| *Stromal cells* | *Stromal 2* | | *1045* | *94* |
| *Stromal cells* | *Stromal 3* | | *394* | *171* |
| *Stromal cells* | *Tip-like ECs* | *Tip-like Endothelial Cells* | *868* | *381* |
| *T cells* | *CD4+ T cells* | *T lymphocytes* | *620* | *25* |
| *T cells* | *CD8+ T cells* | *T lymphocytes* | *1978* | *28* |
| *T cells* | *gamma delta T cells* | *T lymphocytes* | *917* | *265* |
| *T cells* | *NK cells* | *Natural Killer T lymphocytes* | *445* | *248* |
| *T cells* | *Regulatory T cells* | *T lymphocytes* | *2181* | *16* |
| *T cells* | *T follicular helper cells* | *T lymphocytes* | *482* | *200* |
| *T cells* | *T helper 17 cells* | *T lymphocytes* | *996* | *37* |

***Supplementary Table 1:*** *Cell types and subtypes as annotated in the scRNA-seq used as reference for the deconvolution process. We show the number of cells per type in that dataset and the number of genes used for deconvolution. We additionally include a column with comments for further clarification. Of Note: there are also groups of Myeloid, B cells and T cells annotated as unknown at the cell subtype level.*

| Semiquantitative pathologists grading and assessment of the deconvolution results | | | | | | |
|---|---|---|---|---|---|---|
| **Patient** | **CMS1** | **CMS2** | **CMS3** | **CMS4** | **Selected immune features** | **Selected stroma features** |
| S1_Cec A551763 | ++ | ++ | (+) | - | + CD8 T cells<br>+ CD19/CD20 B-Cells | ++ Myofibroblasts |
| S2_Col_R A595688 | - | +++ | * | (+) | + SPP1-positive MA<br>+ Pro-inflammatory MA<br>+ CD4 T-cells<br>+ Treg<br>+ IgG Plasma | ++ Myofibroblasts (perilobular)<br>+ Stroma 1 (non-neoplastic EP)<br>++ Stroma 2 (intralobular)<br>+ Stroma 3 (perilobular) |
| S3_Col_R A416371 | +++ | ++ | (+) | (+) | + SPP1-positive MA<br>+++ CD8 T cells<br>+++ CD19/CD20 B-cells<br>+ NK cells | (+) Myofibroblasts<br>80% of stroma without signature |
| S4_Col_Sig A120838 | - | +++ | - | (+) | + SPP+ MA | ++ Myofibroblasts<br>(+) Stroma 2 (intralobular)<br>+ Stroma 3<br>50% of the stroma without signature |
| S5_Rec A121573 | (+) | +++ | * | (+) | Immune poor | (+) Myofibroblasts<br>+ Stroma 1 (non-neoplastic EP)<br>(+) Stroma 2 (non-neoplastic EP and neoplastic) |
| S6_Rec A938797 | (+) | +++ | * | (+) | + SPP1-positive MA<br>+ Mast cells<br>+ CD4, CD8, T-reg, T-foll helper<br>+ CD19/20 B-cells<br>+ IgG plasma cells | +++ Myofibroblasts<br>+ Stroma 1 (non-neoplastic EP and neoplastic)<br>++ Stroma 2<br>++ Stroma 3 |
| S7_Rec/Sig A798015 | ** | ** | * | ** | + Mast cells<br>+ IgG plasma cells | ++ Myofibroblasts (non-neoplastic CT)<br>+ Stroma 2 (non-neoplastic EP)<br>++ Stroma 3 (non-neoplastic CT) |

***Supplementary Table 2: Semiquantitative grading of our set of CRC samples according to the cell type abundance of the different CMS:*** *Selected immune and stromal features are also specified for each sample. MA= macrophages; R=right, EP=epithelium, CT=connective tissue*

*(-) Highest percentage of cells in tumor/stroma tissue-associated spots <0.1;*

*(+) Percentage of cells per spot 0.2-1 in 10-30% of tumor/stroma tissue-associated spots;*

*+ Percentage of cells per spot 0.2-1 in 31-50% of tumor/stroma tissue-associated spots;*

*++ Percentage of cells per spot 1-5 in 50-100% of tumor tissue-associated spots;*

*+++ Percentage of cells per spot <5 in 50-100% of tumor tissue-associated spots;*

**CMS3 signature (percentage of cells per spot 1-5) in the non-neoplastic mucosa;*

***Low signature (percentage of cells per spot 0.2-1) in the non-neoplastic mucosa/connective tissue.*

| ID | Gene Ratio | BgRatio | pvalue | p.adjust | qvalue | geneID |
|---|---|---|---|---|---|---|
| Patient: *S2_Col_R* | | | | | | |
| ***WNT_BETA_CATENIN_SIGNALING*** | *6/168* | *40/4153* | *0.0048993* | *0.2012755* | *0.1906821* | *NKD1/NOTCH1/LEF1/CTNNB1/PTCH1/AXIN2* |
| ***HEDGEHOG_SIGNALING*** | *5/168* | *34/4153* | *0.0109591* | *0.2012755* | *0.1906821* | *AMOT/ADGRG1/PTCH1/CDK6/VEGFA* |
| ***MYC_TARGETS_V1*** | *15/168* | *196/4153* | *0.0120765* | *0.2012755* | *0.1906821* | *SNRPD1/HSP90AB1/CCT4/ACP1/CCT7/PCBP1/YWHAQ/RAD23B/YWHAE/XPO1/GSPT1/SET/SSBP1/EIF4H/RNPS1* |
| Patient: *S4_Col_Sig* | | | | | | |
| ***WNT_BETA_CATENIN_SIGNALING*** | *6/81* | *40/4153* | *0.0001033* | *0.0044404* | *0.0040219* | *NKD1/AXIN2/KAT2A/CCND2/AXIN1/HDAC5* |
| ***MTORC1_SIGNALING*** | *10/81* | *198/4153* | *0.0046955* | *0.1009540* | *0.0914394* | *PPIA/CYB5B/PHGDH/NUPR1/SQSTM1/CCT6A/EIF2S2/MCM4/HSPD1/SQLE* |
| ***MYC_TARGETS_V1*** | *9/81* | *196/4153* | *0.0132326* | *0.1896669* | *0.1717913* | *PPIA/CBX3/EIF2S2/POLD2/MCM4/HNRNPA2B1/GNL3/HSPD1/SRSF2* |
| Patient: *S5_Rec* | | | | | | |
| ***MYC_TARGETS_V1*** | *18/90* | *196/4153* | *0.0000001* | *0.0000050* | *0.0000044* | *MYC/HSPE1/PGK1/ODC1/PHB/KPNA2/PSMA7/SNRPG/RAN/HSPD1/NME1/NOP56/HPRT1/TFDP1/PRDX4/STARD7/DEK/PRPS2* |
| ***E2F_TARGETS*** | *14/90* | *198/4153* | *0.0000722* | *0.0015530* | *0.0013687* | *MYC/TFRC/DCTPP1/KPNA2/CDC25B/RAN/NME1/AURKA/NOP56/SNRPB/CSE1L/PRDX4/DEK/HMGB3* |
| ***MYC_TARGETS_V2*** | *7/90* | *58/4153* | *0.0002173* | *0.0031145* | *0.0027447* | *MYC/HSPE1/PHB/DCTPP1/HSPD1/NOP56/SORD* |
| ***G2M_CHECKPOINT*** | *11/90* | *194/4153* | *0.0028328* | *0.0285653* | *0.0251738* | *MYC/HSPA8/SLC12A2/ODC1/KPNA2/CDC25B/DKC1/AURKA/UBE2C/TFDP1/HMGB3* |
| ***MTORC1_SIGNALING*** | *11/90* | *198/4153* | *0.0033215* | *0.0285653* | *0.0251738* | *NAMPT/HSPE1/PGK1/TFRC/UFM1/HSPD1/ALDOA/AURKA/HPRT1/PSME3/SORD* |
| ***TNFA_SIGNALING_VIA_NFKB*** | *9/90* | *200/4153* | *0.0279594* | *0.2003756* | *0.1765856* | *CXCL1/CXCL3/NAMPT/MYC/RHOB/PHLDA2/TGIF1/CCL20/ETS2* |
| Patient: *S6_Rec* | | | | | | |
| ***HYPOXIA*** | *5/24* | *195/4153* | *0.0044378* | *0.1302267* | *0.1282368* | *TGFBI/ERRFI1/PPP1R15A/TPI1/GAPDH* |
| ***CHOLESTEROL_HOMEOSTASIS*** | *3/24* | *74/4153* | *0.0084017* | *0.1302267* | *0.1282368* | *ERRFI1/CXCL16/TNFRSF12A* |

***Supplementary Table 3:*** *Significant enrichment results (q<0.2) on the differentially expressed genes between the CMS2 tumors of the different patients.* ***GeneRatio*** *indicates the number of DEG belonging to the enriched category versus the total number of DEG for that patient.* ***BgRatio*** *describes the number of genes in the background belonging to the enriched category versus the total number of background genes.* ***geneID*** *details the DEG matching the enriched category.*

| ID | Gene Ratio | BgRatio | pvalue | p.adjust | qvalue | geneID |
|---|---|---|---|---|---|---|
| **Region:** ***Outer Tumor*** | | | | | | |
| ***EPITHELIAL MESENCHYMAL TRANSITION*** | *44/97* | *188/3726* | *0* | *0* | *0* | *COL6A2/SPARC/FBLN1/FN1/COL3A1/LUM/MGP/COL1A1/COL1A2/THY1/BGN/TAGLN/TIMP3/POSTN/SFRP4/VCAN/ACTA2/COL5A2/VIM/HTRA1/CCN1/DCN/FSTL1/GREM1/TPM2/COL4A1/CALD1/IGFBP4/FLNA/CTHRC1/MMP2/DPYSL3/COL4A2/COL6A3/TIMP1/MYL9/ELN/LGALS1/COL5A1/THBS2/IGFBP3/FBLN2/ITGB1/NID2* |
| ***COAGULATION*** | *17/97* | *102/3726* | *0* | *0* | *0* | *C3/SPARC/FN1/MMP11/APOC1/C1R/MMP9/TIMP3/CTSK/A2M/C1S/HTRA1/MMP2/TIMP1/SERPING1/PECAM1/CTSB* |
| ***MYOGENESIS*** | *15/97* | *148/3726* | *0.0000044* | *0.0000561* | *0.0000513* | *COL6A2/SPARC/COL3A1/COL1A1/TAGLN/IGFBP7/TPM2/AEBP1/COL15A1/COL4A2/COL6A3/LSP1/IGFBP3/WWTR1/ITGB1* |
| ***ANGIOGENESIS*** | *7/97* | *30/3726* | *0.0000081* | *0.0000773* | *0.0000706* | *COL3A1/LUM/POSTN/VCAN/COL5A2/FSTL1/TIMP1* |
| ***COMPLEMENT*** | *13/97* | *167/3726* | *0.0003201* | *0.0024328* | *0.0022239* | *C3/FN1/APOC1/C1R/TIMP2/C1S/COL4A2/TIMP1/SERPING1/PRCP/ANXA5/CTSB/CTSD* |
| **Region:** ***Intermediary Tumor*** | | | | | | |
| **No significant results** | | | | | | |
| **Region:** ***Inner Tumor*** | | | | | | |
| ***HYPOXIA*** | *19/102* | *177/3726* | *0.0000002* | *0.0000083* | *0.0000067* | *ATF3/P4HA1/VEGFA/LDHA/BHLHE40/SLC2A1/P4HA2/DDIT4/ENO1/NDRG1/ALDOA/JUN/PGK1/TPI1/MIF/ALDOC/IER3/GAPDH/SULT2B1* |
| ***CHOLESTEROL HOMEOSTASIS*** | *10/102* | *70/3726* | *0.0000153* | *0.0002581* | *0.0002081* | *SCD/ATF3/LDLR/FASN/ALDOC/ACTG1/ACSS2/CYP51A1/S100A11/LGALS3* |
| ***MTORC1 SIGNALING*** | *17/102* | *196/3726* | *0.0000165* | *0.0002581* | *0.0002081* | *INSIG1/SCD/P4HA1/LDHA/BHLHE40/SLC2A1/LDLR/DDIT4/ENO1/BTG2/ALDOA/UCHL5/PGK1/TPI1/GAPDH/CYP51A1/SERP1* |
| ***GLYCOLYSIS*** | *16/102* | *183/3726* | *0.0000273* | *0.0003207* | *0.0002586* | *P4HA1/VEGFA/LDHA/P4HA2/DDIT4/ENO1/TFF3/PKM/ELF3/SDHC/ALDOA/PGK1/TPI1/MIF/IER3/CLDN3* |
| ***TNFA SIGNALING VIA NFKB*** | *16/102* | *191/3726* | *0.0000465* | *0.0004370* | *0.0003523* | *CCL20/ATF3/AREG/VEGFA/EGR3/BHLHE40/LDLR/BTG2/EGR1/JUN/SNN/ZC3H12A/IER3/TNFAIP2/BIRC3/IFIT2* |
| ***ESTROGEN RESPONSE EARLY*** | *13/102* | *183/3726* | *0.0012558* | *0.0098369* | *0.0079312* | *AREG/EGR3/BHLHE40/SLC2A1/TFF3/FASN/ELF3/TFF1/KCNK5/SULT2B1/SLC22A5/MUC1/PEX11A* |

***Supplementary Table 4:*** *Significant enrichment results (p-value <0.01) on the differentially expressed genes between the different anatomical regions of the CMS2 tumor in the S2_Col_R_Rep1 sample.* ***GeneRatio*** *indicates the number of DEG belonging to the enriched category versus the total number of DEG for that patient.* ***BgRatio*** *describes the number of genes in the background belonging to the enriched category versus the total number of background genes.* ***geneID*** *details the DEG matching the enriched category.*

| ID | Gene Ratio | BgRatio | pvalue | p.adjust | qvalue | geneID |
|---|---|---|---|---|---|---|
| **Region:** ***Sub-Cluster 1*** | | | | | | |
| ***TNFA SIGNALING VIA NFKB*** | *49/410* | *200/4346* | *0* | *0* | *0* | *TRIB1/CXCL1/AREG/PLAUR/NFKBIE/RHOB/KYNU/CXCL3/KLF4/GADD45B/MXD1/MYC/JAG1/FOS/BCL3/LDLR/JUNB/JUN/ATF3/LIF/MCL1/ZFP36/BHLHE40/IRF1/CEBPD/CCNL1/ZC3H12A/KLF10/IER3/CXCL2/IER2/B4GALT1/RNF19B/CDKN1A/ID2/DUSP2/PHLDA2/KLF6/NFE2L2/EDN1/LAMB3/TAP1/DUSP1/NFKBIA/BTG1/TGIF1/VEGFA/BTG2/NR4A1* |
| ***CHOLESTEROL HOMEOSTASIS*** | *21/410* | *74/4346* | *0.0000024* | *0.0000585* | *0.0000427* | *PLAUR/FDFT1/PLSCR1/ANTXR2/SQLE/ACTG1/CLU/FABP5/S100A11/FDPS/JAG1/HMGCS1/LDLR/MAL2/ATF3/HMGCR/LGALS3/ERRFI1/NSDHL/CYP51A1/TMEM97* |
| ***P53 PATHWAY*** | *37/410* | *196/4346* | *0.0000224* | *0.0003660* | *0.0002673* | *GPX2/TOB1/TM4SF1/ITGB4/HINT1/IRAK1/PROCR/RAP2B/EPHA2/KLF4/MXD1/SDC1/NDRG1/TXNIP/FOS/PRMT2/JUN/EPS8L2/ATF3/LIF/STEAP3/IER3/RNF19B/CDKN1A/SFN/SLC7A11/PLXNB2/AK1/CTSD/TAP1/VAMP8/SEC61A1/BTG1/CEBPA/DDIT4/BTG2/ZFP36L1* |
| ***UV RESPONSE UP*** | *29/410* | *156/4346* | *0.0002339* | *0.0028652* | *0.0020927* | *GGH/DGAT1/ATP6V1C1/GRINA/OLFM1/EPCAM/RHOB/STK25/IGFBP2/POLR2H/FOS/DNAJA1/DNAJB1/JUNB/ATF3/IRF1/FKBP4/FURIN/CXCL2/HLA-F/SPR/WIZ/TAP1/TUBA4A/GLS/NFKBIA/BTG1/BTG2/NR4A1* |
| ***GLYCOLYSIS*** | *34/410* | *198/4346* | *0.0003366* | *0.0032983* | *0.0024090* | *PPP2CB/IDH1/HSPA5/AK3/SDC1/QSOX1/EGFR/MET/GPC4/ELF3/CHPF/HDLBP/FKBP4/MDH1/B3GNT3/IER3/CHPF2/B4GALT1/IL13RA1/ENO1/HK2/RBCK1/DSC2/NSDHL/SLC16A3/GCLC/TGFBI/AURKA/SOX9/STMN1/CLDN3/PYGB/VEGFA/DDIT4* |
| ***TGF BETA SIGNALING*** | *13/410* | *54/4346* | *0.0011700* | *0.0090233* | *0.0065906* | *ID1/SLC20A1/SPTBN1/FNTA/JUNB/FKBP1A/KLF10/FURIN/ID2/THBS1/ID3/SKIL/TGIF1* |
| ***INTERFERON ALPHA RESPONSE*** | *19/410* | *96/4346* | *0.0012890* | *0.0090233* | *0.0065906* | *LY6E/HLA-C/LGALS3BP/PLSCR1/IFI27/PROCR/IFI35/GBP4/TXNIP/PARP14/PSME2/OGFR/IRF1/LAP3/PARP9/ADAR/CD47/TAP1/IL4R* |
| ***HYPOXIA*** | *32/410* | *200/4346* | *0.0017387* | *0.0106494* | *0.0077783* | *ANXA2/GAA/PLAUR/JMJD6/HSPA5/NDRG1/GRHPR/FOS/EGFR/GPC4/JUN/ATF3/ZFP36/BHLHE40/HDLBP/IER3/ENO1/CDKN1A/IDS/HK2/SLC2A1/ERRFI1/PFKL/KLF6/TGFBI/DUSP1/ILVBL/BTG1/PLAC8/VEGFA/DDIT4/SELENBP1* |
| ***MTORC1 SIGNALING*** | *31/410* | *198/4346* | *0.0028992* | *0.0157847* | *0.0115291* | *IFRD1/ACSL3/SQLE/IDH1/HSPE1/HSPA5/HSPD1/SORD/HMGCS1/LDLR/GSR/GLA/BHLHE40/SLC1A5/CCT6A/PSME3/HMGCR/ENO1/CDKN1A/HK2/SLC2A1/SLC7A11/PFKL/CYP51A1/ACLY/TMEM97/GCLC/TUBA4A/AURKA/DDIT4/BTG2* |
| ***APOPTOSIS*** | *26/410* | *159/4346* | *0.0034054* | *0.0166862* | *0.0121876* | *SLC20A1/TSPO/PEA15/RHOB/HSPB1* |

| | | | | | | |
|---|---|---|---|---|---|---|
| | | | | | | /CLU/GADD45B/BCAP31/TXNIP/DNAJA1/GSR/JUN/ATF3/MCL1/IRF1/TIMP1/IER3/CDKN1A/LGALS3/PMAIP1/SPTAN1/ERBB2/TAP1/CDC25B/IFNGR1/BTG2 |
| ***IL2 STAT5 SIGNALING*** | *30/410* | *199/4346* | *0.0059433* | *0.0264748* | *0.0193371* | *PLEC/MYO1C/ITGAV/ITGA6/CAPG/PLSCR1/RHOB/GADD45B/MXD1/GBP4/MYC/NDRG1/NFKBIZ/CDCP1/CTSZ/AHCY/ARL4A/PRNP/LIF/BHLHE40/SLC1A5/HUWE1/FURIN/ST3GAL4/HK2/TNFRSF21/ANXA4/KLF6/IL4R/IFNGR1* |
| Region: ***Sub-Cluster 2*** | | | | | | |
| ***MYC TARGETS V1*** | *17/88* | *196/4346* | *0.0000002* | *0.0000101* | *0.0000093* | *HNRNPA1/PA2G4/TFDP1/RRM1/HPRT1/DEK/NDUFAB1/CLNS1A/NAP1L1/SLC25A3/LDHA/SYNCRIP/NPM1/RNPS1/SNRPD2/PCNA/COX5A* |
| ***OXIDATIVE PHOSPHORYLATION*** | *16/88* | *200/4346* | *0.0000017* | *0.0000359* | *0.0000333* | *COX6A1/ATP5MG/CS/NDUFAB1/COX4I1/SLC25A5/ATP5F1B/SLC25A3/LDHA/NDUFC2/MRPL11/UQCRC2/COX5A/ATP5F1C/LDHB/FDX1* |
| ***E2F TARGETS*** | *12/88* | *198/4346* | *0.0005546* | *0.0077642* | *0.0071998* | *PA2G4/HMGB2/DEK/NAP1L1/SMC1A/HMGA1/DCTPP1/SYNCRIP/MCM3/TMPO/PCNA/TP53* |
| Region: ***Sub-Cluster 3*** | | | | | | |
| ***COAGULATION*** | *12/51* | *138/4346* | *0* | *0.0000013* | *0.0000011* | *CFB/MMP3/C2/FN1/MMP1/C3/C1R/SERPINA1/PLAU/CTSK/MMP2/A2M* |
| ***KRAS SIGNALING UP*** | *8/51* | *199/4346* | *0.0019551* | *0.0235473* | *0.0198293* | *CFB/TMEM176A/PIGR/TMEM176B/CCL20/MAP7/IGFBP3/PLAU* |
| ***COMPLEMENT*** | *8/51* | *200/4346* | *0.0020183* | *0.0235473* | *0.0198293* | *CFB/C2/FN1/C3/MMP12/C1R/S100A9/SERPINA1* |
| ***INTERFERON ALPHA RESPONSE*** | *5/51* | *96/4346* | *0.0049834* | *0.0436049* | *0.0367199* | *IFITM3/IFITM2/NCOA7/IFI30/CD74* |
| ***INTERFERON GAMMA RESPONSE*** | *7/51* | *198/4346* | *0.0077083* | *0.0474365* | *0.0399466* | *CFB/SOD2/IFITM3/IFITM2/C1R/IFI30/CD74* |
| ***EPITHELIAL MESENCHYMAL TRANSITION*** | *7/51* | *200/4346* | *0.0081320* | *0.0474365* | *0.0399466* | *MMP3/FN1/MMP1/TNC/IGFBP3/CXCL8/MMP2* |

***Supplementary Table 5:*** *Significant enrichment results (p-value <0.01) on the differentially expressed genes between the different sub-clustered regions of the CMS2 tumor in the S5_Rec_Rep1 sample.* ***GeneRatio*** *indicates the number of DEG belonging to the enriched category versus the total number of DEG for that patient.* ***BgRatio*** *describes the number of genes in the background belonging to the enriched category versus the total number of background genes.* ***geneID*** *details the DEG matching the enriched category.*

| *Pathologists spot categorization* | |
|---|---|
| **Category** | **Categorization criteria** |
| ***non neoplastic epithelium*** | *>90% non neoplastic epithelial cells* |
| ***submucosa*** | *>90% submucosa* |
| ***epithelium&submucosa*** | *epithelium and submucosa: mixed spots 10-90% epithelium, 10-90% submucosa* |
| ***tumor*** | *>90% neoplastic epithelial cells* |
| ***stroma_fibroblastic_IC low*** | *fibroblast rich, collagen poor stroma, IC <10%, low nuclear density* |
| ***stroma_fibroblastic_IC med*** | *20- 50% IC, moderate nuclear density, HE: myofibroblast-like morphology* |
| ***stroma_fibroblastic_IC high*** | *51- 80% IC, high nuclear density* |
| ***stroma_desmoplastic_IC low*** | *fibroblast poor, collagen rich stroma, IC <10%, low nuclear density* |
| ***stroma_desmoplastic_IC med to high*** | *IC >10%, low to moderate nuclear density* |
| ***tumor&stroma_IC med to high*** | *mixed spots, <90% tumor, <90% stroma, IC in stroma 20-90%* |
| ***tumor&stroma_IC low*** | *mixed spots, <90% tumor, <90% stroma, IC in stroma <10%* |
| ***IC aggregate_submucosa*** | *follicular immune cell aggregates>90% IC* |
| ***IC aggregate_stroma or muscularis*** | *follicular immune cell aggregates>90% IC* |
| ***muscularis_IC med to high*** | *smooth muscle cells, IC content 20-90%* |
| ***exclude*** | *spots of poor tissue quality or not classifiable* |

***Supplementary Table 6:*** *Pathologists spot categorization and categorization criteria. Each spot was manually assigned to a category in the HE stained OCT tissue section the Loupe Browser. IC=immune cells*
