## Supplementary Note 1 for "Charting the Heterogeneity of Colorectal Cancer Consensus Molecular Subtypes using Spatial Transcriptomics"

#### Comparison of deconvolution results using the scRNA-seq data from the Korean or the Belgian cohort

In the main manuscript, we presented the results of the Cell2Location[1] deconvolution approach when using as reference the cell type annotations from a scRNAseq dataset derived from a Korean cohort of 23 patients. This Korean dataset proceeds from the study by Lee et al.[2], in which an additional scRNAseq dataset from a Belgian cohort of 6 patients was independently analyzed and annotated. Both datasets present some relevant differences, such as an enrichment of fibroblasts and myeloid cells in the Belgian dataset. Besides, a few cell types were only identified in one of the datasets. This is the case of the tuft cells, which were only detected in the Belgian dataset. Consequently and in line with the original publication, we performed the deconvolution in parallel with each dataset. In this supplementary note, we compared the results yielded by both procedures.

We first compared the deconvolution results across major cell types (Fig. 1). Overall the cell type proportions present similar trends across our set of ST samples when using the references derived from both datasets. However, there are some noticeable and relevant differences. Excluding the sample containing only non neoplastic tissue, there is a predominant increase in the proportions of tumor cells mapped into our samples when using the Belgian cohort. For some samples, the intensification of tumorigenic signals seems to be mainly at the expense of the normal epithelial cells. This is for instance the case in the S5_Rec sample. In other samples, such as in S3_Col_R, there is a marked decrease in the predicted abundance of B cells.

In order to get a better understanding of these discrepancies, we explored the proportions at the cell subtype level. We first focused on the different CMS tumor cell populations (Fig. 2). Overall, the results are highly comparable using either of the two reference datasets. The samples referred to as CMS2 tumors in the main manuscript also display a clear predominance of this tumor cell type when using the Belgian reference. Concretely, the percentages of CMS2 tumor cells as compared to the total number of tumor cells in these patients were: S2_Col_R (89%), S4_Col_Sig (92%), S5_Rec (86%) and S6_Rec (88%). In addition, the S1_Cec (30% and 68%) and S3_Col_R (45% and 51%) patients maintain a mixed proportion of CMS1 and CMS2 tumor cells. Interestingly, for the Belgian reference, the samples labeled as CMS2 tumors present a more prominent CMS1 signal, whereas the mixed CMS1-CMS2 tumors display a slight decrease in the proportion of CMS1 cell abundances. To get further insights on these differences, we explored the tumor cell abundance spatial mappings per spot in our samples (Fig. 3-9). As expected from the global tumor proportions, CMS1 cells displayed the most prominent differences between both references. In the samples labeled as CMS2 tumors, the number of predicted CMS1 cells was very limited and did not display a clear spatial distribution when using the Korean reference. However, using the Belgian reference conducted to stronger CMS1 signals overlapping to a considerable degree with the anatomical regions where the presence of CMS2 cells was predicted. In the CMS1-CMS2 mixed tumors, the predicted spatial location of CMS1 cells matched between references. Across our set of samples, the predicted anatomical location of CMS2, CMS3 and CMS4 cells matched to a large extent for both used references. There were nevertheless differences in their estimated abundance, mainly for CMS2, which presented an increased number of cells predicted per spot when using the Belgian reference. This growth in the number of predicted CMS2 tumor cells can account for the global larger proportions of tumor cells detected per sample (see Fig.1).

We subsequently compared the deconvolution results provided by the different references for immune cells. The estimated proportions per sample were highly comparable for T- (Fig. 10) and B-cells (Fig. 11). Despite some discrepancies in the abundance of $CD8^+$ T cells in some samples, such as in *S3_Col*, their predicted anatomical locations overlapped using either reference dataset (Fig. 12). The comparison between the proportions of myeloid cells was not so straightforward due to differences at the annotations level between both reference datasets (Fig 13). The main difference lies on the $SPP1^+$ macrophages which were split into $SPP1^+A$ and $SPP1^+B$ macrophages in the annotations of the scRNA-seq dataset derived from the Belgian cohort. Interestingly, $SPP1^+A$ and $SPP1^+B$ macrophages were mapped to separate anatomical locations in the tumor proximity (Fig 14). Finally, we contrasted the results for the main stromal cell types (Fig 15). The overall results are highly consistent between both reference datasets with some minor but relevant differences. For instance, in the S1_Cec_Rep1 sample, a very limited content of Stromal 1 cells was detected when using the Korean dataset as compared to the results using the Belgian dataset (Fig 16). Of note, the latter results matched with the regions annotated as stromal by the pathologist.

In summary, there was a large degree of similarity between the deconvolution results using either of the two references. In the context of our study, the most remarkable discrepancies arose from the spatial mapping of CMS1 tumor cells across our set of samples. We hypothesized that the annotations from the Korean dataset allowed the deconvolution to better distinguish between the signatures of CMS1 and CMS2 tumor cells. In samples S2_Col_R, S4_Col_Sig, S5_Rec and S6_Rec, the morphological features and the immune cold landscape of their tumors were suggestive of CMS2 carcinomas. These facts led to our choice of using in the main manuscript the deconvolution results obtained when using as reference the cell type annotations from the Korean cohort. Moreover, the number of individuals (23 versus 6) and annotated cells (65,362 versus 27,971) was larger in the Korean dataset than in the Belgian one, consequently reflecting a more comprehensive and diverse dataset for deconvolution purposes. On the other hand, some stromal populations derived from the Belgian dataset appear to better map their expected anatomical regions in a couple of our ST samples. We presume that the genetic background can have an influence on these results as our samples from German origin are expected to be more similar to the Belgian reference.

### References

1. Kleshchevnikov, V. *et al.* Cell2location maps fine-grained cell types in spatial transcriptomics. *Nat. Biotechnol.* **40**, 661–671 (2022).
2. Lee, H.-O. *et al.* Lineage-dependent gene expression programs influence the immune landscape of colorectal cancer. *Nat. Genet.* **52**, 594–603 (2020).

### Figures

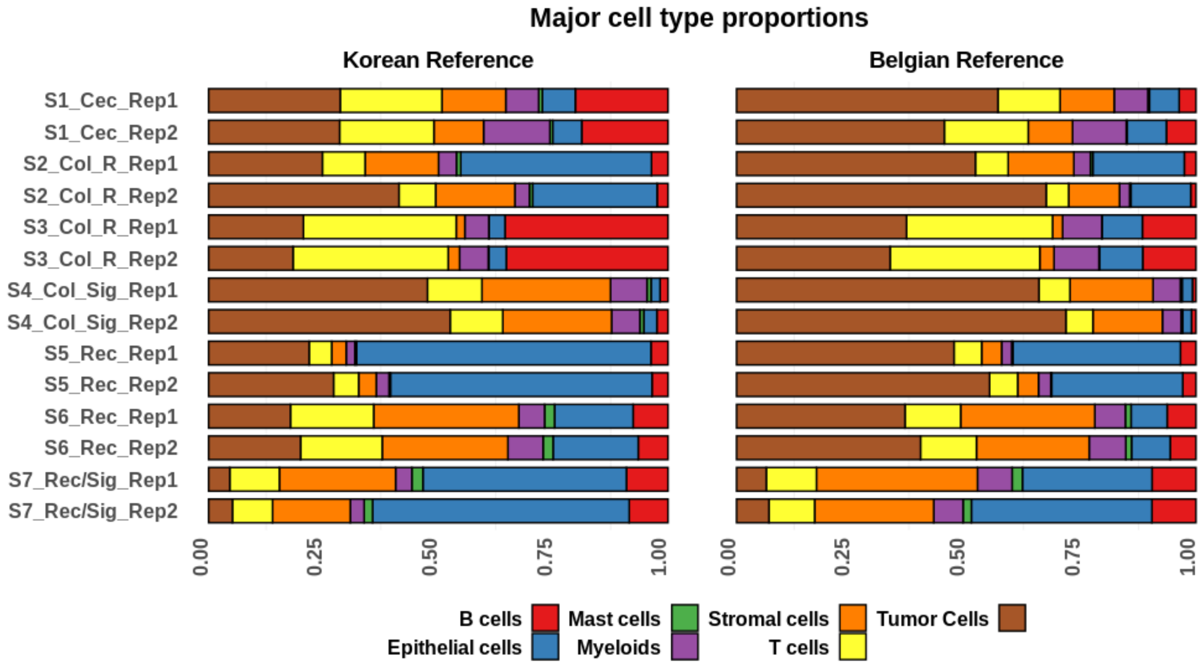


***Figure 1:*** *Proportions of major cell classes per sample as estimated by the results of the deconvolution approach when using as reference the scRNA-seq dataset derived from either the Korean or the Belgian cohort.*

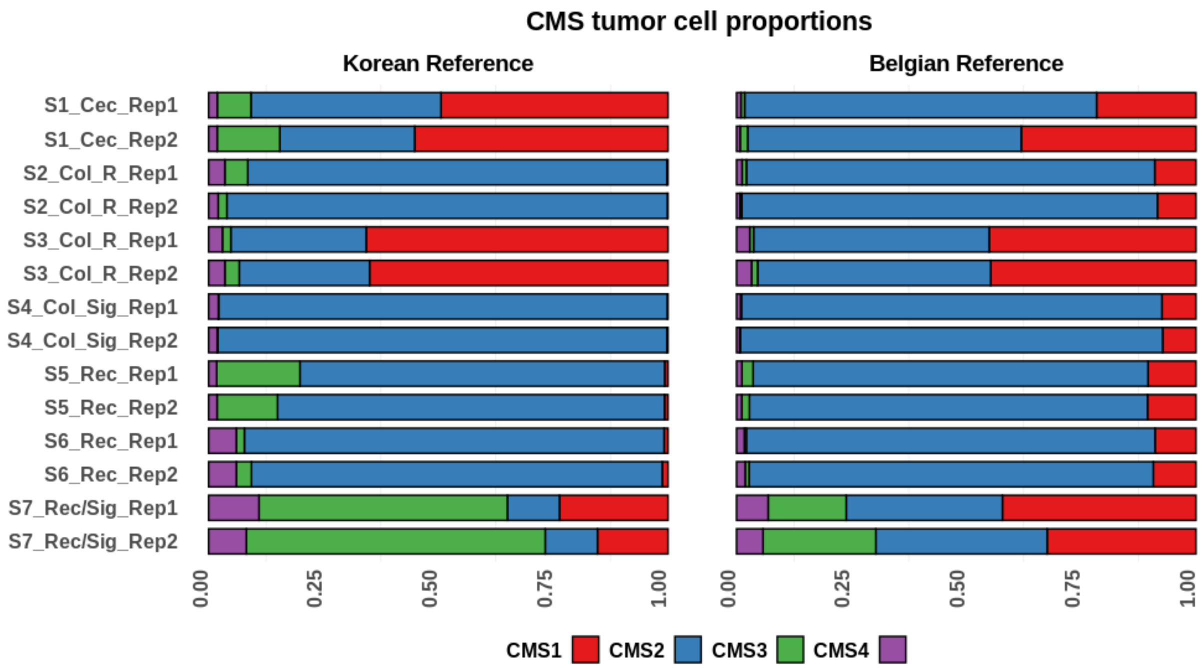


***Figure 2:*** *Proportions of CMS tumor cells per sample as estimated by the results of the deconvolution approach when using as reference the scRNA-seq dataset derived from either the Korean or the Belgian cohort.*

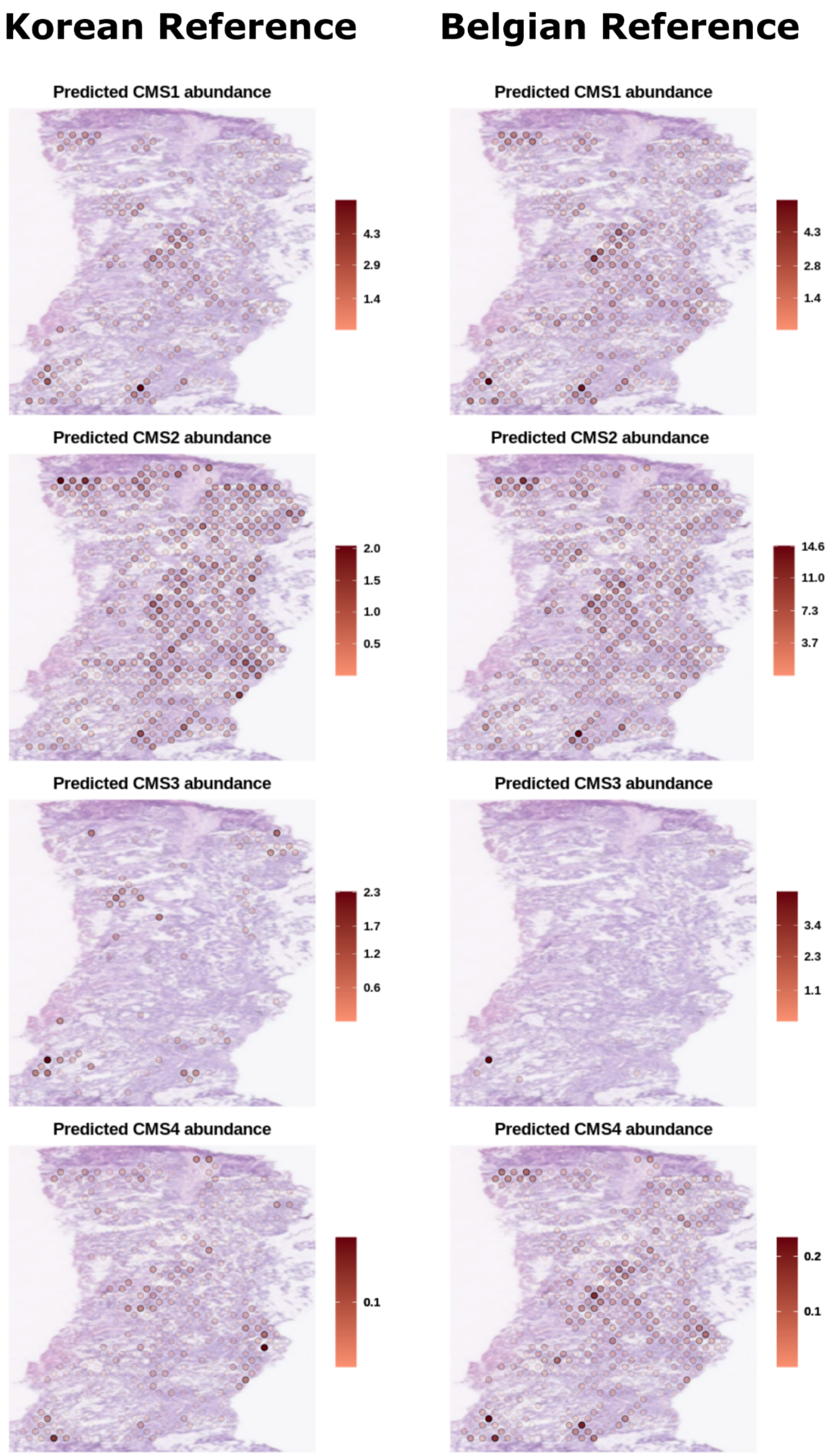


***Figure 3:*** *Spatial mapping of the different CMS tumor cell abundances as predicted by the deconvolution when using as reference either the Korean (left panels) or the Belgian (right panels) datasets in the S1_Cec_Rep1 sample.*

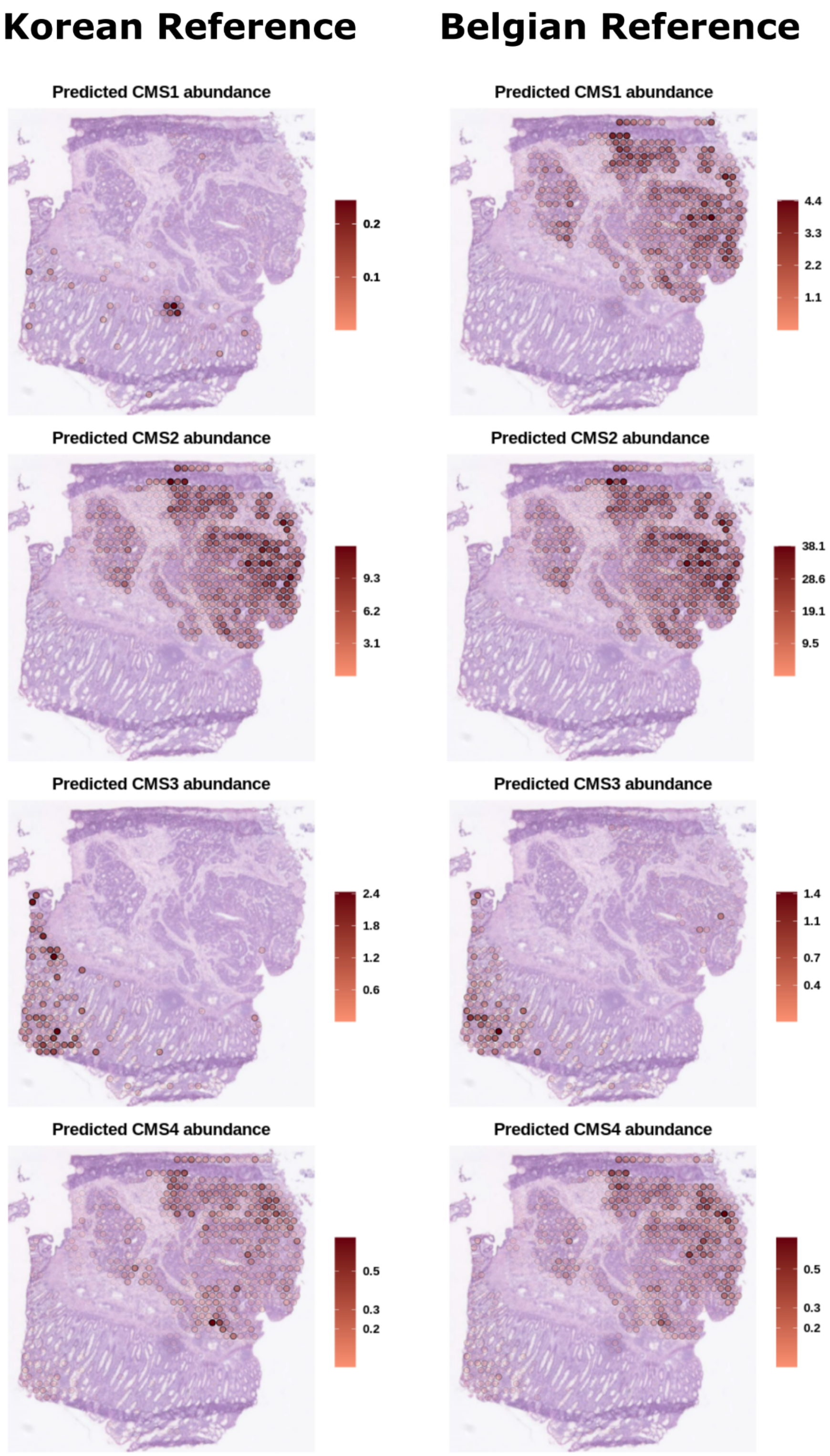


***Figure 4:*** *Spatial mapping of the different CMS tumor cell abundances as predicted by the deconvolution when using as reference either the Korean (left panels) or the Belgian (right panels) datasets in the S2_Col_R_Rep1 sample.*

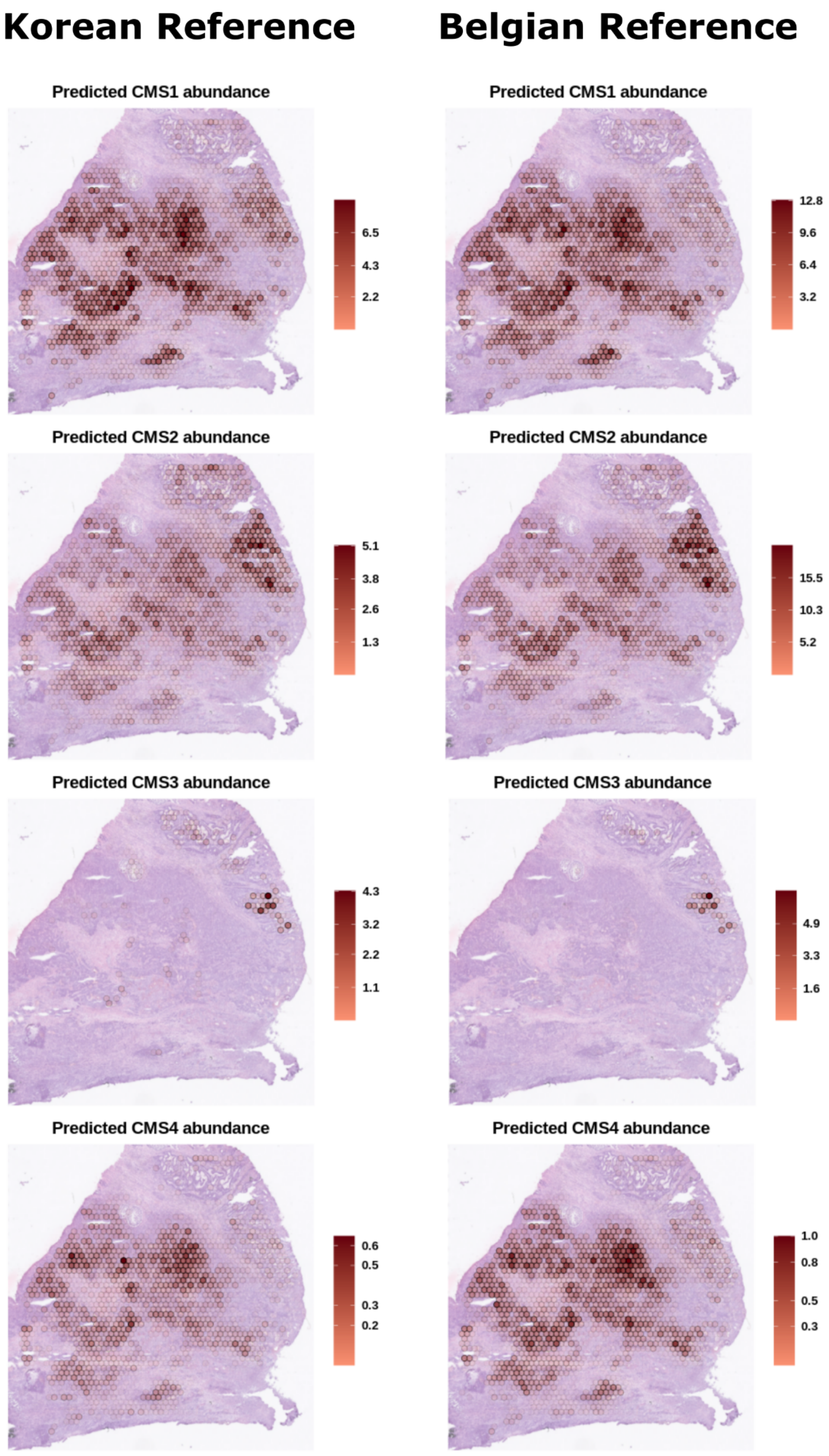


***Figure 5:*** *Spatial mapping of the different CMS tumor cell abundances as predicted by the deconvolution when using as reference either the Korean (left panels) or the Belgian (right panels) datasets in the S3_Col_Rep1 sample.*

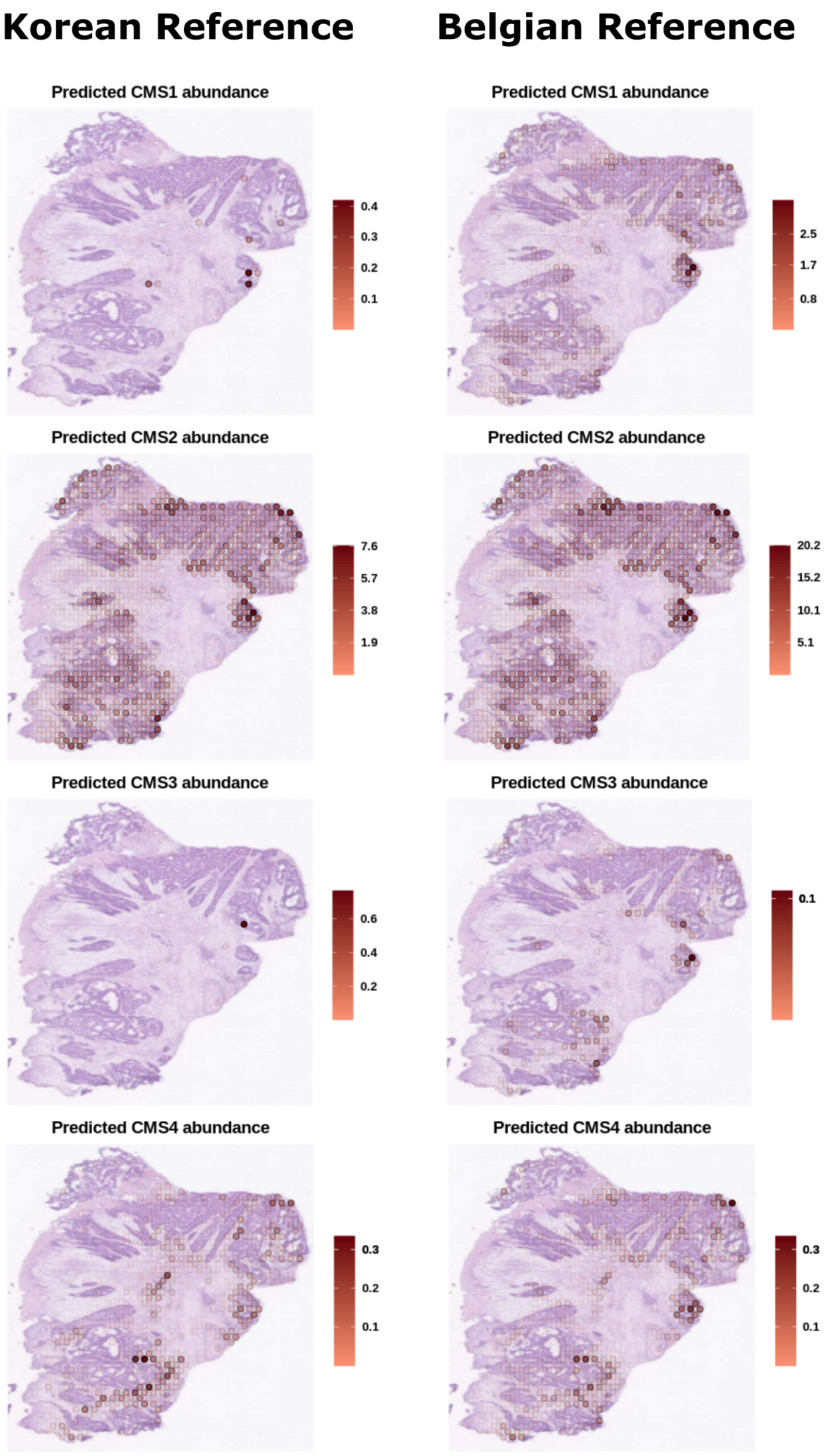


***Figure 6:*** *Spatial mapping of the different CMS tumor cell abundances as predicted by the deconvolution when using as reference either the Korean (left panels) or the Belgian (right panels) datasets in the S4_Col_Sig_Rep2 sample.*

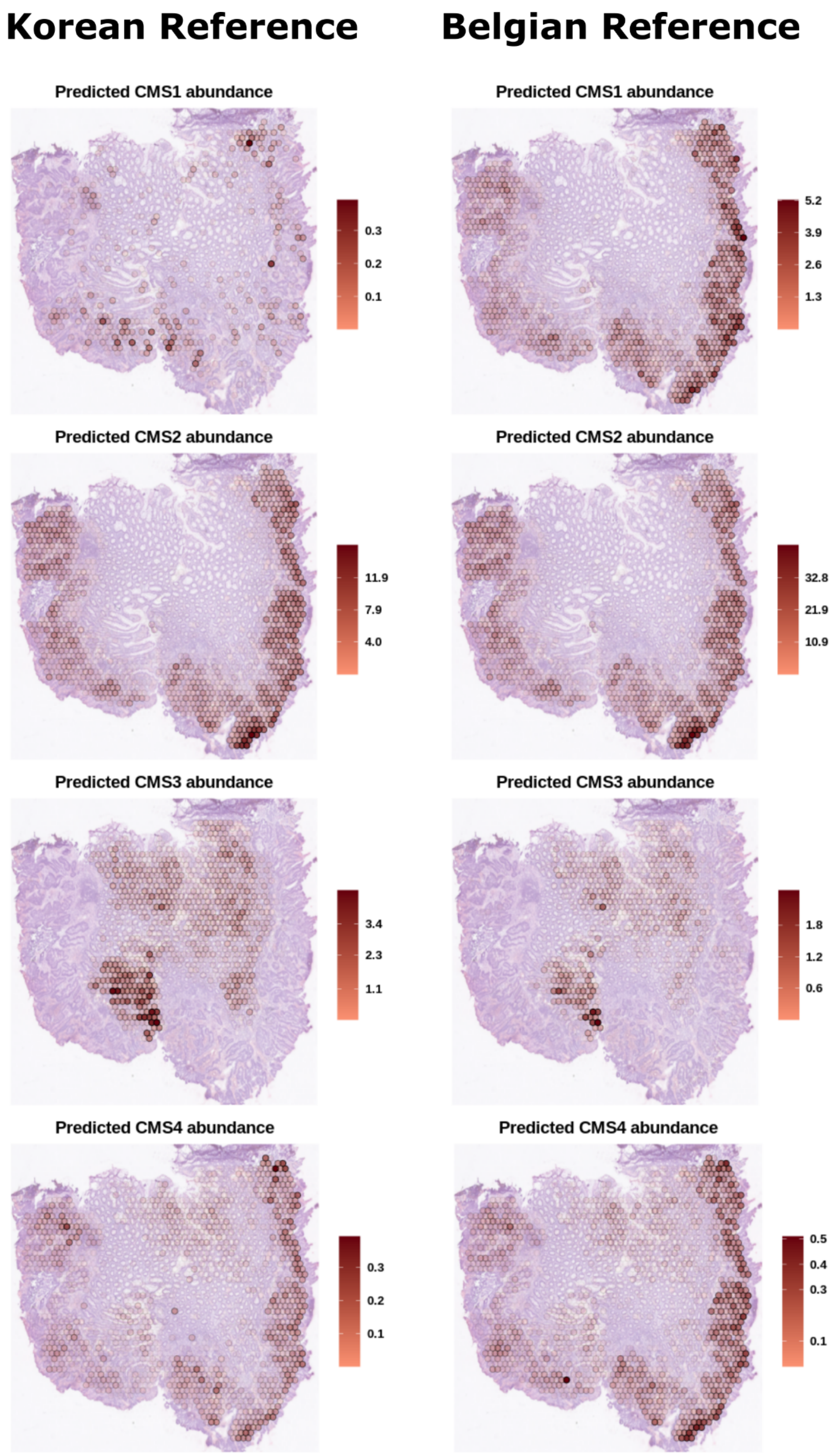


***Figure 7:*** *Spatial mapping of the different CMS tumor cell abundances as predicted by the deconvolution when using as reference either the Korean (left panels) or the Belgian (right panels) datasets in the S5_Rec_Rep1 sample.*

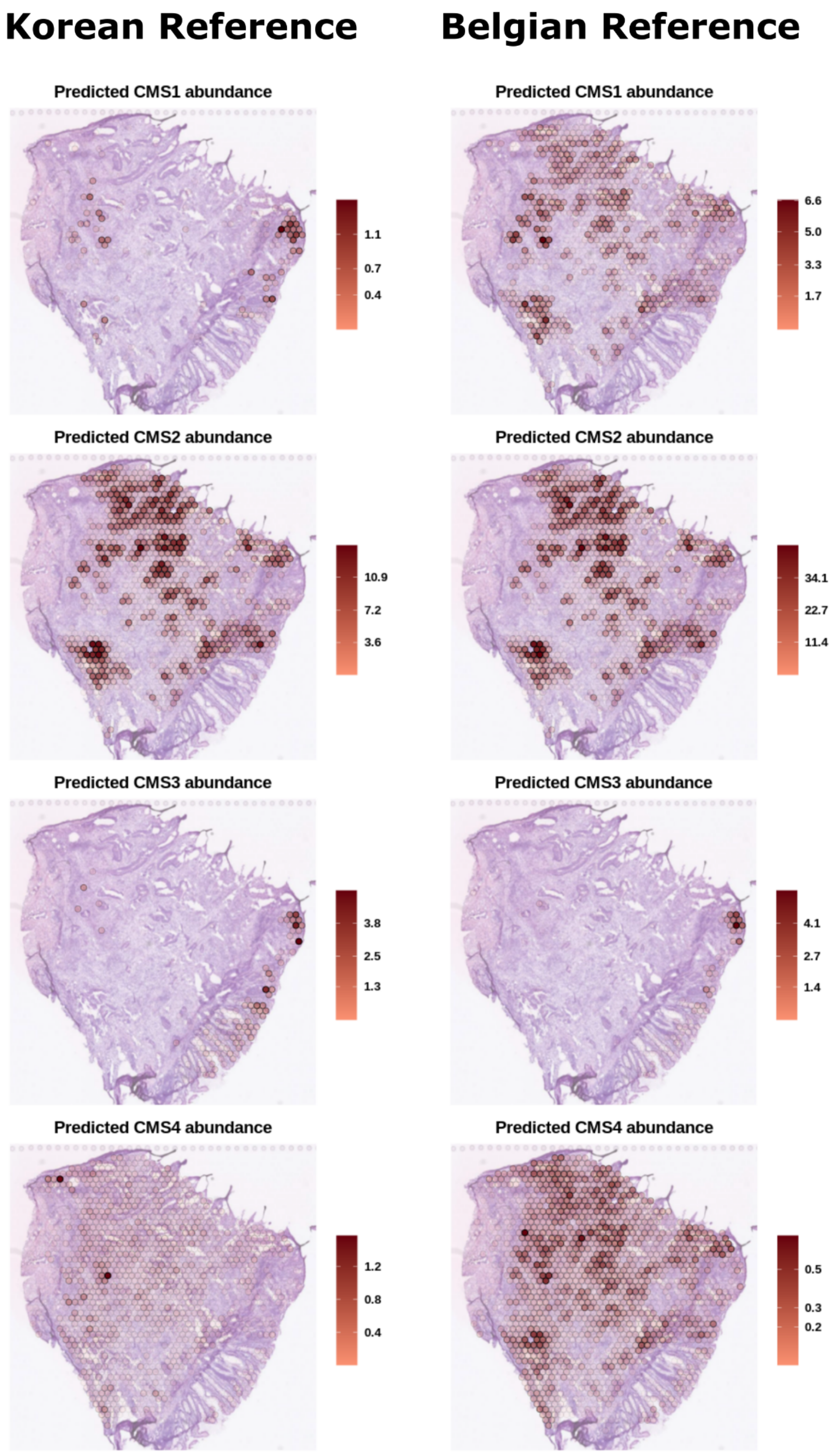


***Figure 8:*** *Spatial mapping of the different CMS tumor cell abundances as predicted by the deconvolution when using as reference either the Korean (left panels) or the Belgian (right panels) datasets in the S6_Rec_Rep2 sample.*

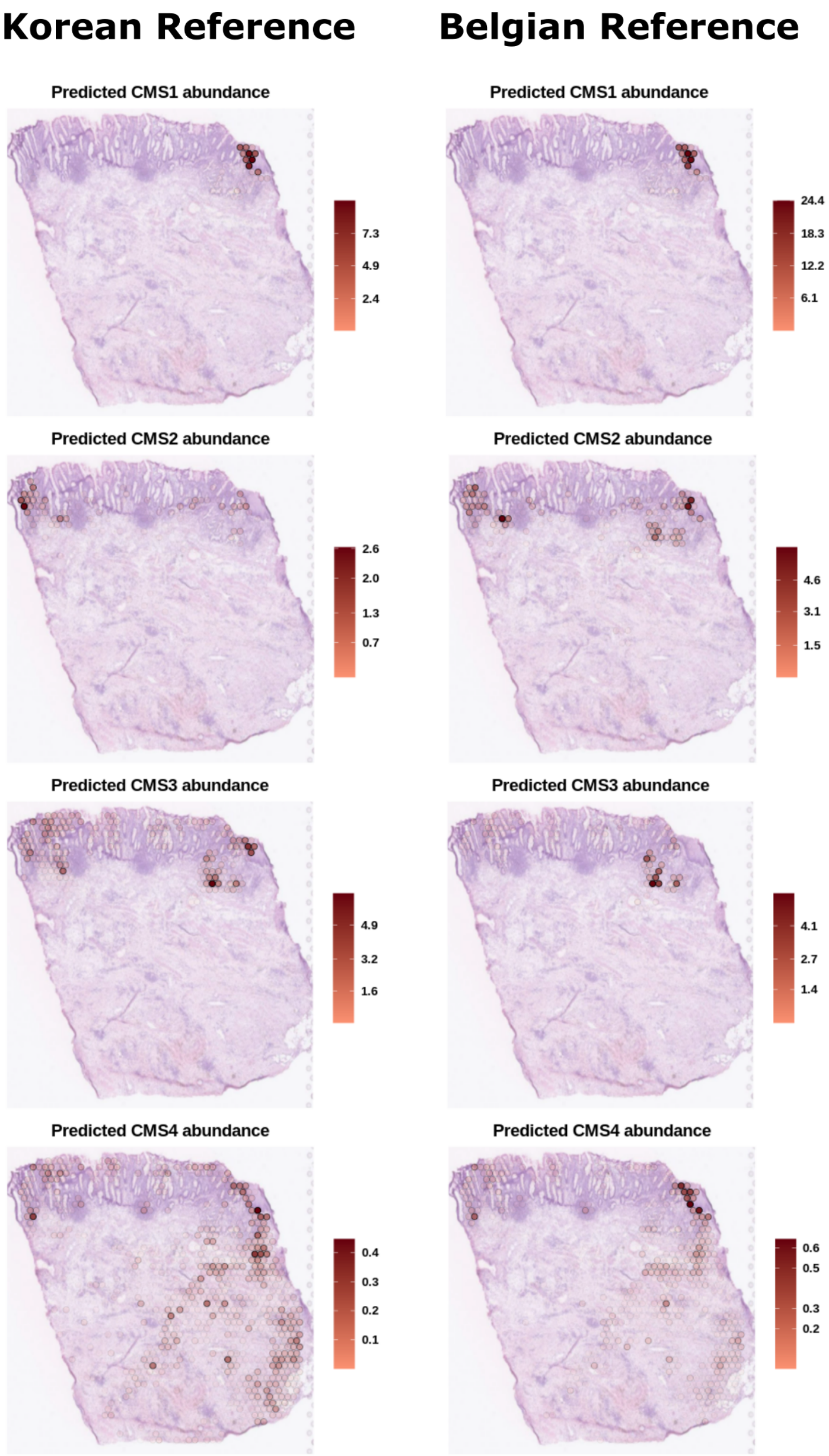


***Figure 9:*** *Spatial mapping of the different CMS tumor cell abundances as predicted by the deconvolution when using as reference either the Korean (left panels) or the Belgian (right panels) datasets in the S7_Rec/Sig_Rep1 sample.*

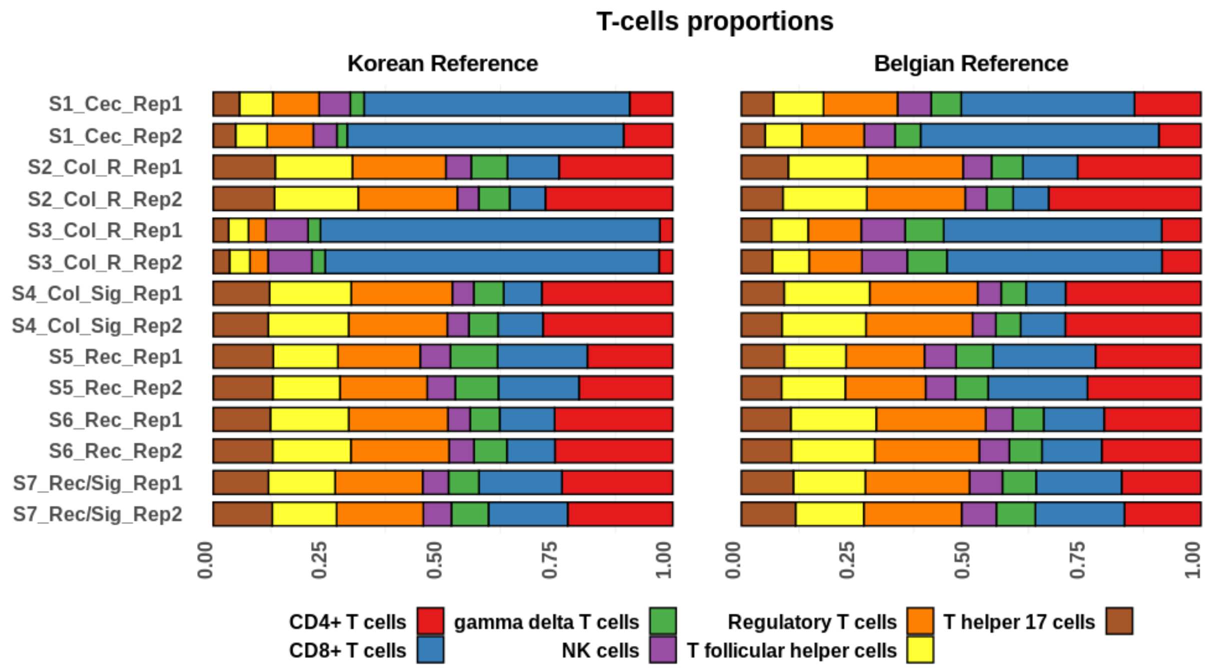


***Figure 10:*** *Proportions of T-cells subtypes per sample as estimated by the results of the deconvolution approach when using as reference the scRNA-seq dataset derived from either the Korean or the Belgian cohort.*

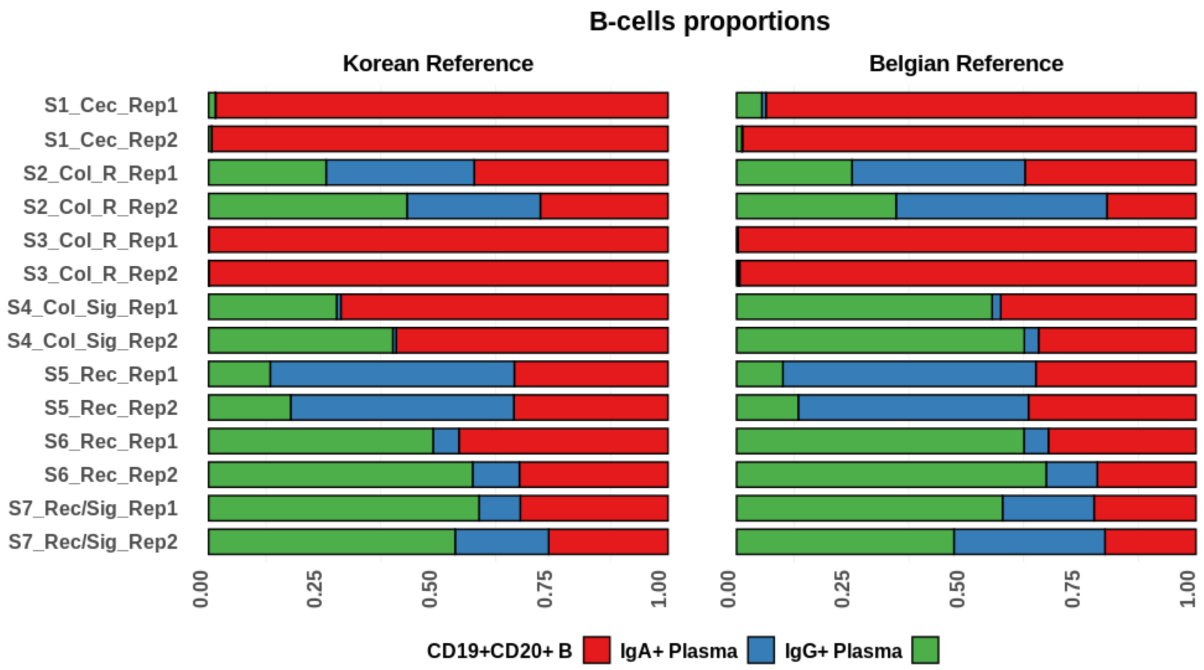


***Figure 11:*** *Proportions of B-cells subtypes per sample as estimated by the results of the deconvolution approach when using as reference the scRNA-seq dataset derived from either the Korean or the Belgian cohort.*

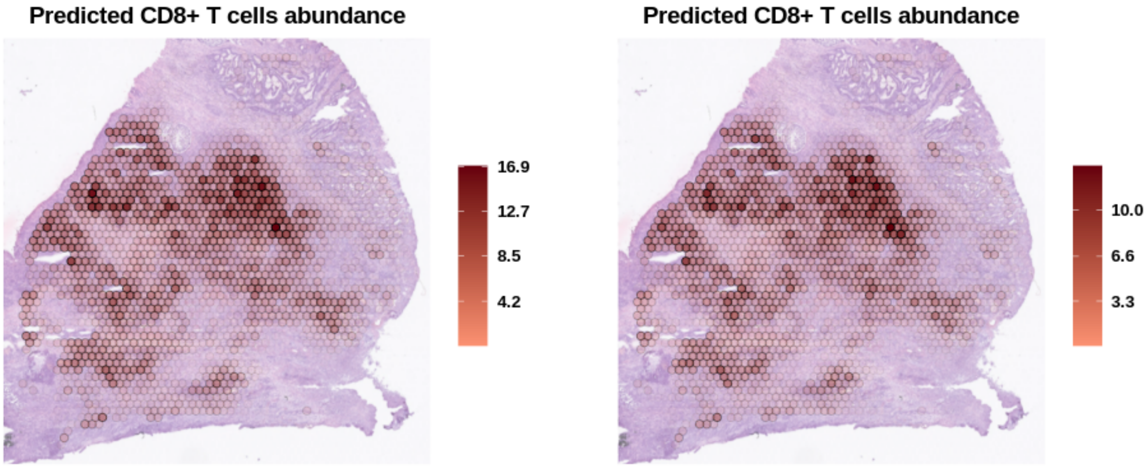


***Figure 12:*** *Spatial mapping of the CD8+ T cells abundance as predicted by the deconvolution when using as reference either the Korean (left panel) or the Belgian (right panel) datasets in the S3_Col_Rep1 sample.*

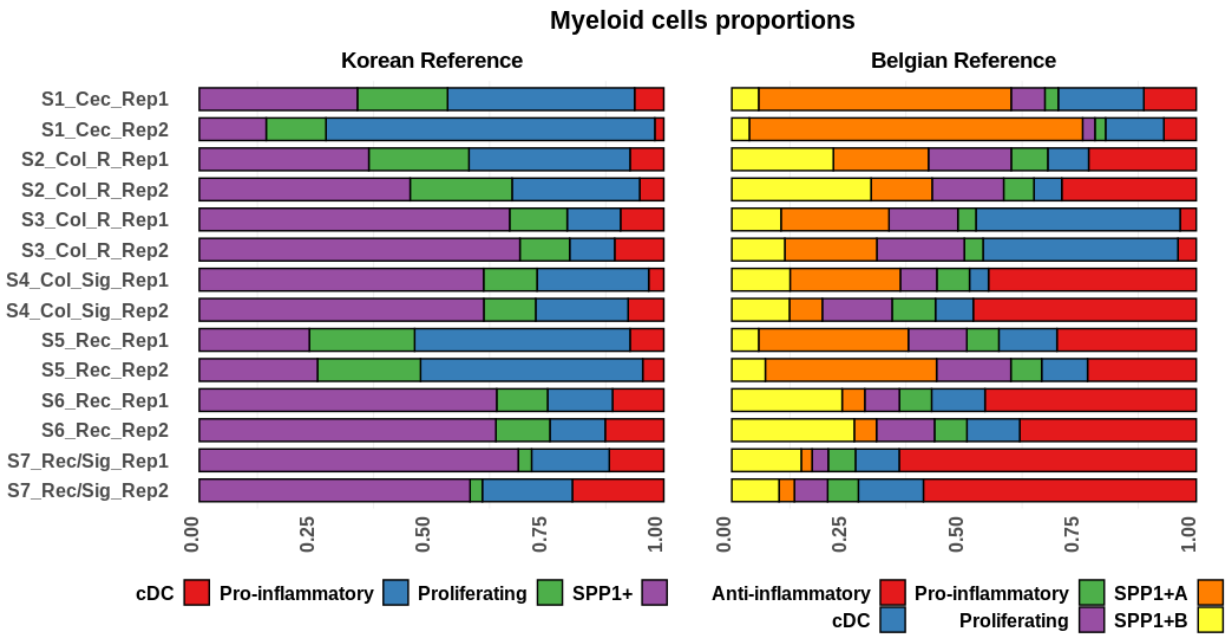


***Figure 13:*** *Proportions of myeloid cell subtypes per sample as estimated by the results of the deconvolution approach when using as reference the scRNA-seq dataset derived from either the Korean or the Belgian cohort.*

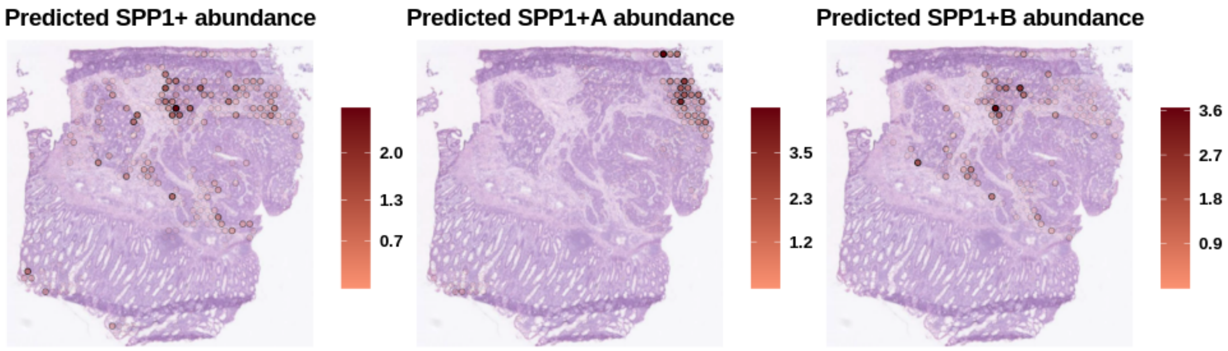


***Figure 14:*** *Spatial mapping of the cell abundance of SPP1+ macrophages as predicted by the deconvolution when using as reference either the Korean (left panel) or the Belgian (central and right panels) datasets in the S2_ Col_ R_ Rep1 sample.*

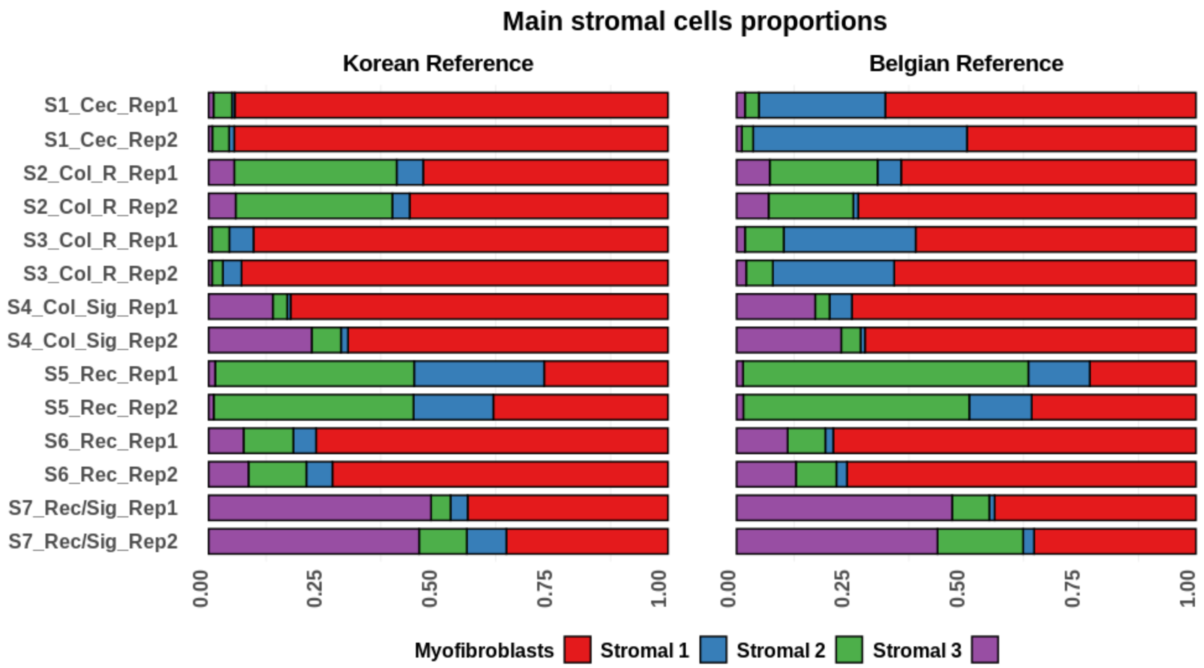


***Figure 15:*** *Proportions of the main stromal cell subtypes per sample as estimated by the results of the deconvolution approach when using as reference the scRNA-seq dataset derived from either the Korean or the Belgian cohort.*

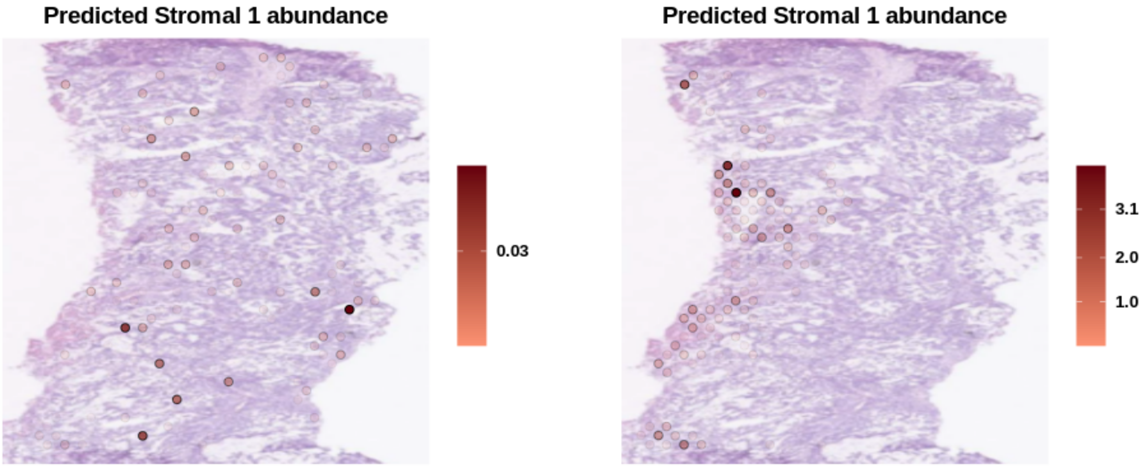


***Figure 16:*** *Spatial mapping of the cell abundance of Stromal 1 cells as predicted by the deconvolution when using as reference either the Korean (left panel) or the Belgian (right panel) datasets in the S1_ Cec_Rep1 sample.*
